# Variation in precipitation drives differences in interactions and short-term transient instability between grassland functional groups: a stage-structured community approach

**DOI:** 10.1101/2024.10.07.617067

**Authors:** Aryaman Gupta, Samuel Gascoigne, György Barabás, Man Qi, Erola Fenollosa, Rachael Thornley, Christina Hernandez, Andy Hector, Roberto Salguero-Gómez

## Abstract

Climate change is expected to increase the frequency and severity of precipitation extremes, causing droughts and flooding. Consequently, grassland communities are forecasted to become increasingly unstable. To predict grassland responses, we need empirical information together with models that reliably extrapolate community dynamics from those observations. However, such prediction is challenging because community models typically simulate long-term (asymptotic) performance, and thus potentially neglect their short-term (transient) performance. Here, we use data from a precipitation experiment performed over eight years to model both short- and long-term responses of three functional groups – grasses, legumes, and non-leguminous forbs – to precipitation extremes. We use multi-functional-group Integral Projection Models and pseudospectral theory, to track grassland community dynamics. We show that the percentage-cover-stage-structure of functional groups shapes their transient instability, and that inter-functional-group interactions are competitive under increased precipitation but facilitative under decreased precipitation. IPMs and pseudospectra enable forecasting of how functional-group-stage-structure drives responses to climatic extremes.

## Introduction

Understanding how abiotic disturbances shape communities is a pressing concern for ecology and conservation biology (Hutchinson 1958, Lovelock & Margulis 1974, Miller & Travis 1996, Sutherland et al. 2013, Kraft et al. 2014, Hou & Wang 2023). Climate change is expected to increase the frequency and severity of abiotic disturbances such as extreme precipitation (Knapp et al. 2015). These increases will place unprecedented pressure on ecological communities worldwide (Teshome et al. 2020, Pecl et al. 2017). Consequently, the composition of communities, and the population structure of species within those communities are expected to become increasingly unstable (Montoya & Raffaelli 2010, Gaüzère et al. 2018). Hence, it is crucial that community models capture these increasingly uncertain conditions.

To model communities, ecologists often use differential equations that track how variables of interest (*e.g.,* population sizes of multiple species, intra- and inter-specific competition coefficients) interact (Verhoef & Morin 2009; Breckling et al. 2011; Sieber & Malchow 2011). These models are then used to quantify community composition stability (Yang et al. 2023), tipping points (Jiang et al. 2019), or transient (*i.e.,* short-term) responses (Gorban 2020, Barabás 2024). However, these models often neglect population structure, focusing only on population size to track species performance (but see Moll & Brown 2008, Fujiwara et al. 2011, Barabás et al. 2014, de Roos 2021, Johnson et al. 2024). Yet, population structure can drive community dynamics (de Roos 2020) and can be important for conservation and management (Thomassen et al. 2018), so we need models that capture this.

In demography, Integral Projection Models (IPMs; Easterling et al. 2000) have become widely used tools to study changes in population structure (Merow et al. 2014b). Due to their usefulness, empirically parameterized IPMs now exist for over 300 species (Levin et al. 2022). IPMs are discrete-time, continuous-state models that track a species’ vital rates (*e.g.,* survival/mortality, growth/shrinkage, recruitment; Fig. 1; Merow et al. 2014a). Despite their potential to explicitly link a species’ vital rate to the population structure of coexisting species, IPMs have not been widely used in community ecology (but see Adler et al. 2010, Kayal et al. 2019, Peirce et al. 2023). This lack of adoption can be explained in part by the fact that modelling interactions between structured populations places high demands on parametrisation and computation (Rossberg and Farnsworth 2010). The problem is especially difficult to resolve when modelling realistic levels of biodiversity. Nevertheless, understanding how stage-structure and community dynamics shape each other promises to enrich our understanding of both population and community ecology (Loreau 2010).

**Figure 1.**
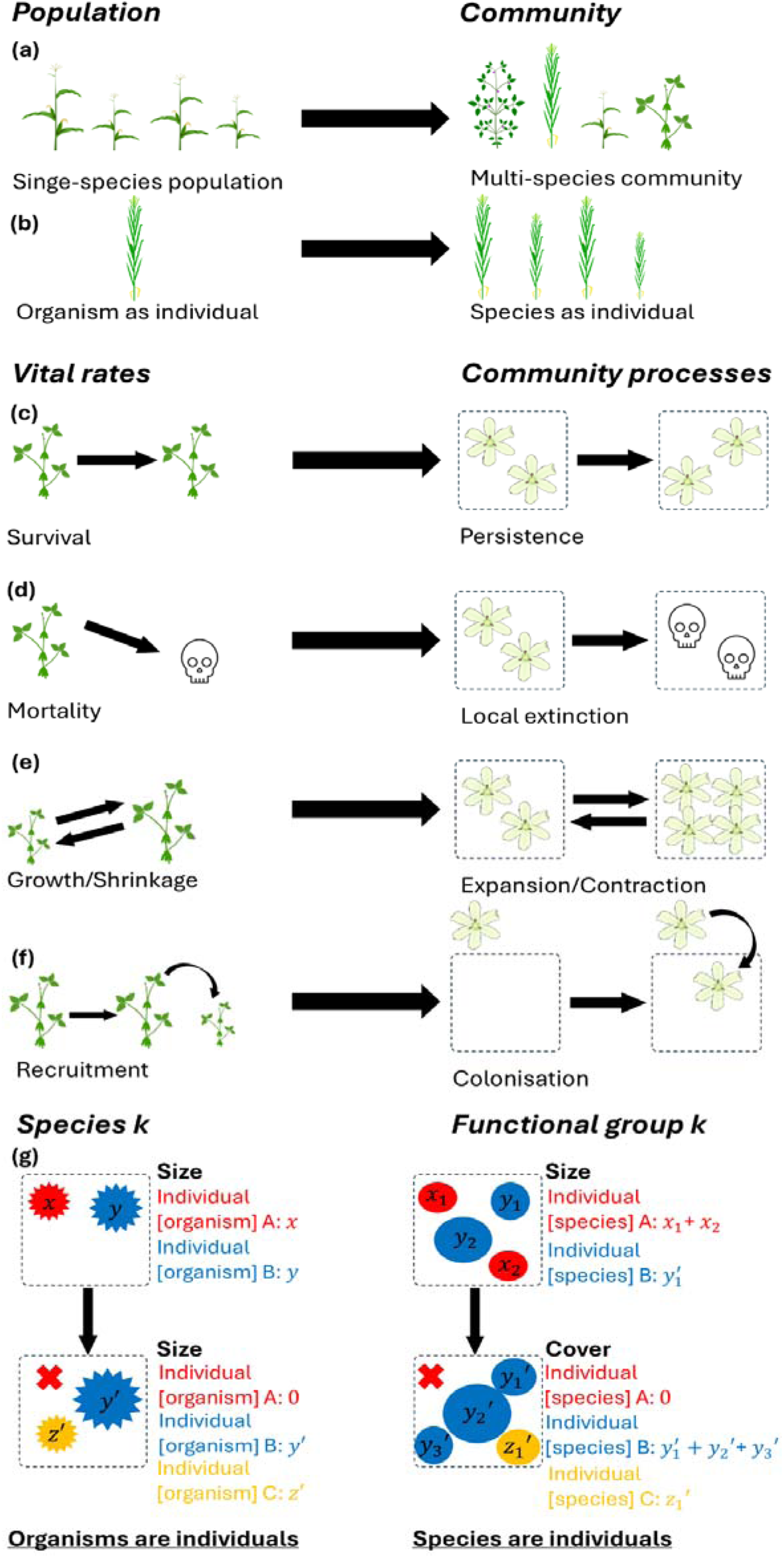
The shift in perspective from population processes to community processes. In this study, we deploy this shift to examine the impacts of experimentally manipulated precipitation on the stage-structure and transient dynamics of communities. The shift implies mapping how vital rates within a population draw analogous processes at the community level. This analogy is drawn by examining single *vs* multiple species populations (a – b), and their corresponding shifts in population to community processes (c – f). Further, instead of considering the performance of individual organisms in a species, we consider the performance of individual species in a functional group (g). (a) Instead of a single-species population, here we consider a community as the population. (b) Instead of individual organisms, here we consider species as individuals. (c) Instead of survival of individuals, we consider persistence of species. (d) Instead of mortality of individuals, we consider local extinction of species. (e) Instead of growth and shrinkage of individuals, we consider the expansion and contraction of species’ covers. (f) Instead of recruitment of new offspring, we consider colonisation of new species. (g) Instead of taking organisms as individuals to track how a species changes over time, we take species as individuals to track how a functional group changes over time. Under the (demographic) perspective of organisms as individuals, organism A experiences mortality (shifting from size x to size 0), organism B experiences growth (from size y to size y’), and organism C is recruited by the population (initial size z’). Under the perspective of species as individuals, species A experiences local extinction (shifting from cover x_1_ + x_2_ to cover 0), species B experiences growth (from cover y_1_ + y_2_ to cover y_1_’ + y_2_’ + y_3_’), and organism C colonises the plot (initial cover z_1_’). Flower icon by DBCLS https://togotv.dbcls.jp/en/pics.html is licensed under CC-BY 4.0 Unported https://creativecommons.org/licenses/by/4.0/. All other icons by Daniel Carvalho https://figshare.com/authors/Plant_Illustrations/3773596 are licensed under CC-BY 4.0 Unported https://creativecommons.org/licenses/by/4.0/.

One can model community dynamics using IPMs, while avoiding the computational costs of modelling interactions between large numbers of structured populations, by relaxing the definition of an ‘individual’. Indeed, the founder of modern plant population ecology, J. Harper, stated that ‘a single plant is a population of modules’ (1977). Similarly, here we argue that a *community* can be considered a *population of species,* or, considering species function into the community, a *population of functional groups*. From this perspective, each species is an ‘individual’ whose cover (as a community ecologist would quantify in the field) equals its ‘size’ (as a population ecologist would do in a population; Fig. 1a-b; Ghiselin 1974, Hull 1976). Species experience community processes analogous to vital rates, as reviewed in Fig. 1c-f. Thus, instead of representing the individuals changing in a single-species population, here we use IPMs to represent the species cover change in a functional group (Fig. 1g). Our IPMS are thus not size-structured but rather percentage-cover structured. The state-vector being predicted is not how many individual organisms in a species there are of a given size-range, but rather how many species there are of a given percentage-cover in a functional group. Admittedly, our proposal to treat communities as populations of species risks abstracting away important phenomena arising from intra-functional-group interactions (Rubio & Swenson 2024). However, this potential cost is offset by the benefit that our approach enables us to parametrise interactions between large numbers of species by a small number of functional-group level parameters (Chalmandrier et al. 2021). This shift in perspective is further licensed by work showing that functional-group level models can match the performance of species-level models in predicting community dynamics (Steinmann et al. 2009, Chalmandrier et al. 2021, Rubio & Swenson 2022).

Ecological models are commonly analysed at steady state (Noy-Meier 1974, Caswell 2019, Van Meerbeck et al. 2021). The properties of a system at equilibrium reveal the long-term fate of a perturbation away from that equilibrium. This way, one can partially characterise the system’s behaviour close to an equilibrium point. However, sometimes more important than asymptotic outcomes are the behaviour for short- and intermediate time scales after the perturbation, as studied in pseudospectral theory (Trefethen & Embree 2005, Barabás & Allesina 2015, Caravelli & Staniczenko 2016). This theory characterises how the eigenvalues of a matrix respond to changes in its entries and provides criteria for whether an otherwise stable system is expected to first transiently amplify a dynamical perturbation before the system returns to equilibrium. This is the first main novelty of our study – by using IPMs in conjunction with pseudospectral theory, we can gain insight into how grassland communities transiently amplify perturbations before they settle into equilibrium. The second main novelty is our species-as-individual perspective, which enables us to upscale IPMs (originally developed to study demography [Easterling et al. 2000]), to instead study how stage-structure shapes community dynamics.

Grassland responses to perturbations such as extreme precipitation are important to understand, given the global significance of their biodiversity and the ecosystem services they provide (Hopkins & Prado, 2007, Fay et al., 2008, Franklin et al. 2016, Jackson et al. 2024). Little attention has been paid to how extreme precipitation and stage-structure together influence transient community dynamics in grasslands (but see deSiervo et al. 2023). However, we understand related mechanisms of grassland community dynamics better, which can therefore serve to corroborate the results of our pseudospectral analysis of multi-functional-group IPMs. As per the Stress Gradient Hypothesis (Bertness & Callaway 1994), we expect interspecific interactions to be primarily competitive under increased precipitation and facilitative under reduced precipitation (Hallet et al. 2019, Lima et al. 2022). We can also broadly infer which precipitation conditions favour which functional groups. Under increased precipitation, grass cover would be expected to be more abundant than the other two functional groups, as grasses grow taller and faster than the legumes and forbs, thus overshadowing them (Pitt & Heady 1978, Hautier et al. 2009). Under reduced precipitation, legumes and forbs would be expected to be more abundant than grasses as they are physiologically more drought-tolerant (Hallet et al. 2017). If our models produce results in agreement with these *a priori* predictions, that would license our trust in the novel predictions it makes regarding how stage-structure shapes transient dynamics.

To examine both the transient and long-term stage-dependent dynamics of grassland communities subject to perturbations such as extreme precipitation, we construct a multi-functional-group IPM and analyse it using the theory of pseudospectra. The IPM is parameterised with eight years of abundance data from a DroughtNet site located at Wytham Woods (UK; Fig. 2a). At this site, grassland communities containing three functional groups: grasses, legumes, and non-leguminous forbs (subsequently ‘forbs’) were exposed to irrigation (+50% growing season precipitation), control, and drought (−50% precipitation) treatments, replicated in five blocks. We test the following hypotheses: (H1.1) Interactions will be more competitive under irrigation than control conditions; (H1.2) Interactions will be more facilitative under drought than control conditions; (H2.1) Under the irrigation treatment, grass cover will be more abundant than the other two functional groups (Pitt & Heady 1978); (H2.2) Under the drought treatment, legumes and forbs will be more abundant than grasses; (H3) Community stability will depend on the percentage-cover stage-structure of each functional group; and (H4) Grassland communities have a tendency to exhibit transient instability, whereby a perturbation away from equilibrium is first amplified before the system returns to its steady state.

**Figure 2.**
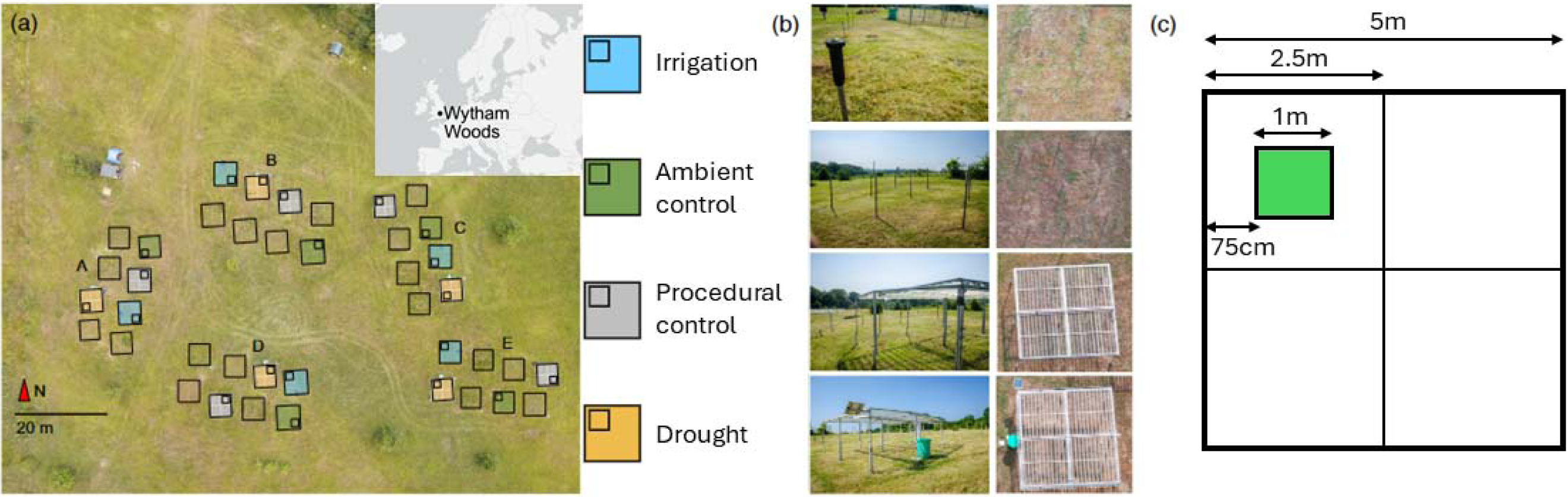
Schematic of the DroughtNet experimental treatments implemented at RainDrop, Wytham Woods. (adapted from Jackson et al. 2024). (a) Distribution of DroughtNet treatments across the site, spatially arranged into five blocks (A – E). The treatments are: irrigation (+50% precipitation; blue), ambient control (green), procedural controls (grey), and drought (−50% precipitation; orange). Each square represents the 5m × 5m permanent plot, and each smaller square within indicates the randomly chosen position of a quadrat from which community data were collected every June from 2016 to 2023. (b) Photographs of the treatment structures for each treatment, from a ground level (left) and aerial (right) view. (c) Design of each 5m × 5m permanent plot, which is subdivided into four quarters (each 2.5m × 2.5m). Percentage cover readings for each species were taken using a 1m × 1m quadrat placed at the centre of the quarter chosen for observation.

## Materials and Methods

### Experimental setup

We conducted our study at the RainDrop (Rainfall and Drought platform) long-term ecological experiment site (https://www.ecologicalcontinuitytrust.org/raindrop). Our site is located at the Upper field site at Wytham Woods, in Oxford, UK (51° 46′ 7.57″ N, 1° 19′ 49.58″ W, 84-167 m a.s.l.; Fig. 2a), which is a calcareous chalkland grassland (Gibson & Brown, 1991). See the Supplementary Materials, Fenollosa et al. (2024), and Jackson et al. (2024) for further details on the study site. The experimental site consists of five randomised blocks (A – E) of four 5 m × 5 m treatment plots (*n* = 20 plots): irrigation (+50% of ambient precipitation, via sprinklers), an ambient control (no treatment and no roof structure), procedural control (a roof structure with inverted gullies that let all natural precipitation through), and drought (−50% of ambient precipitation, via a roof structure with gullies; Fig. 2a, b). Within each plot, we monitored community responses at each plot within a 1 m × 1 m quadrat placed in the middle of one of the four quarters (determined randomly) of the plot. This placement was used to avoid edge effects (Fig. 2c), following Smith et al. (2024).

Both experimental treatments (irrigation and drought) are only applied during the growing season (April – September). The procedural control was established to evaluate whether the shelter structure has any effect on the plot beyond the intended reduction in precipitation. Potential confounding effects included changes in boundary layer dynamics that could influence vegetation via changes in temperature, humidity or light availability. See the supplementary materials for further details. However, recent analysis of the data collected at our study site shows that plots under procedural control have the same community composition and precipitation responses as plots under ambient control (Jackson et al. 2024). Thus, we subsequently merge data from procedural controls and ambient controls and refer to them as ‘control.’

### Data collection

Biotic for three functional groups (grasses, legumes, and forbs) was collected in the summer (June-July) for eight years (2016-2023). The biotic data collected at each permanent plot included species identification (including functional group: legume, grass, or forb) and abundance (measured via the proxy of species percentage cover, from 0% to 100%). Each year, for each plot and block, all species in the 1 m × 1 m quadrat were identified during the growing season peak (mid to late June). Species names were recorded following the International Plant Name Index (IPNI 2024). Abundance data were assessed as the average of the three observers’ estimates each year for each plot and block, to an accuracy of ±0.5%. Due to the three-dimensional structure of the plant community, whereby covers of different species can overlap, the sum of abundance values of all species in a quadrat can exceed 100%. Once the data were collected, we summed the abundance values of all species in each functional group separately. For each functional group *k*, we denote the overall abundance of that group by *N_k_*.

### Model construction

To test whether interspecific interactions were competitive under irrigation (H1.1) and facilitative under drought (H1.2), we constructed models of our community analogues to vital rates for each functional group × treatment. Using version 1.1-35.5 of the *lme4* package (Bates et al. 2024) in version 4.2.2 of R (R core team, 2024), we constructed generalised linear mixed-effect models of persistence, expansion, and colonisation for individual species in each of the 3 functional groups × 3 treatment conditions. These models are the equivalents to the survival, growth/shrinkage, and reproduction vital rate models for individual organisms of a single species in a demographic model (Merow et al. 2014a), respectively. These models will be used later to construct our demographic model (Fig. 1c-e). We tested the effects of inter- and intra-functional-group interactions on functional group persistence, expansion, and colonisation by individual species in each functional group. Colonisation was not dependent on the percentage cover of any particular species because new species colonise from ‘outside,’ and do not rely on existing individuals like how reproduction would in a demographic model. We tested for these effects by including overall levels of abundance (percent cover) of grasses (*N_g_*), legumes (*N_l_*), and forbs (*N_f_*) as fixed explanatory variables. Before analysis, percentage cover, *N_g_* and *N_f_* values were all log transformed to ensure linear and quadratic models could be used to fit the data. Unless stated otherwise, all logarithms in this paper have a base of *e*. Since *N_1_*was zero in two instances, *e*^-1^ was added to each *N_l_* value before they were all log-transformed. Regressions to construct models of persistence, expansion and colonisation were performed on every possible pair (*t*, *t*+1) of abundance data. Under a given treatment, persistence *s_k_*(*x*) was modelled as a continuous variable between 0 and 1 using a logit link function, where *x* is (logged) cover in year *t* for functional group k, by the formula

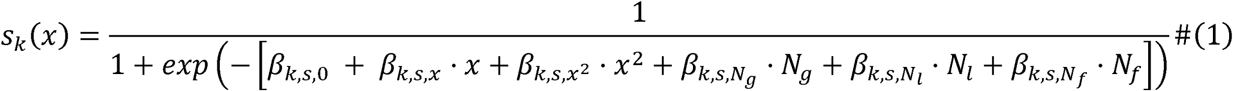

Expansion *g_k_* (*x*, *y*) was modelled using a Gaussian distribution *G* (μ*_g, k_* (x), σ_g, k_(x)), where *y* is the (logged) cover in year *t+1,* by the formula

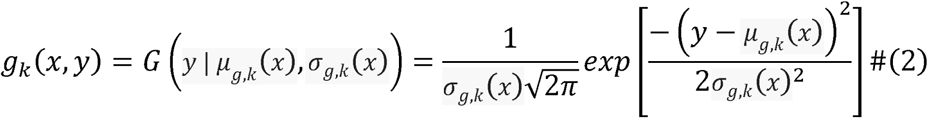

Here, the mean μ*_g, k_* (*x*) and standard deviation σ*_g, k_* (*x*) were given by

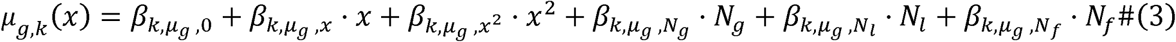

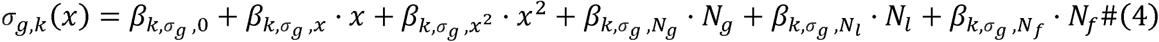

Colonisation *f_k_*(*y*) was assumed to be lognormally distributed. Therefore, colonisation was also modelled using a Gaussian distribution *G* (*μ_f_*, *σ_f_*) by the formula

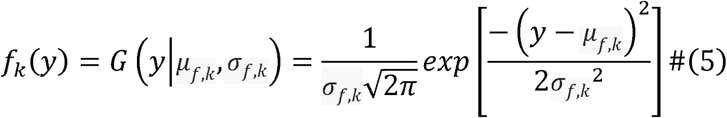

Here, the mean *μ_f, k_* and standard deviation *σ_f, k_* were given by

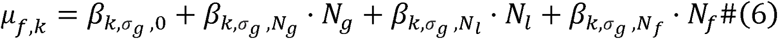

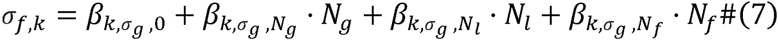

We compared a battery of models including (only for persistence/expansion) linear and squared abundance of individual species, and (for colonisation too) overall abundances of grasses, legumes, and forbs. This battery was constructed using the MuMIn package (Bartoń 2024). From this battery, models with lowest AICc (by at most two AICc scores) were identified. Among these models, the model with lowest AICc which produced biologically plausible results (*e.g.*, that percentage cover does not grow forever) were chosen (see Table S1–S45). Table 1 shows the models we selected. For excluded terms, the *β* coefficient was set to 0 in the formulas above. These models allowed us to test hypotheses H1.1 and H1.2. To find support for H1.1, coefficients of the N_k_ terms would need to be negative in models for functional groups under the irrigation treatment. To find support for H1.2, coefficients of the *N_k_* terms would need to be positive in models for functional groups under the drought treatment.

**Table 1.**
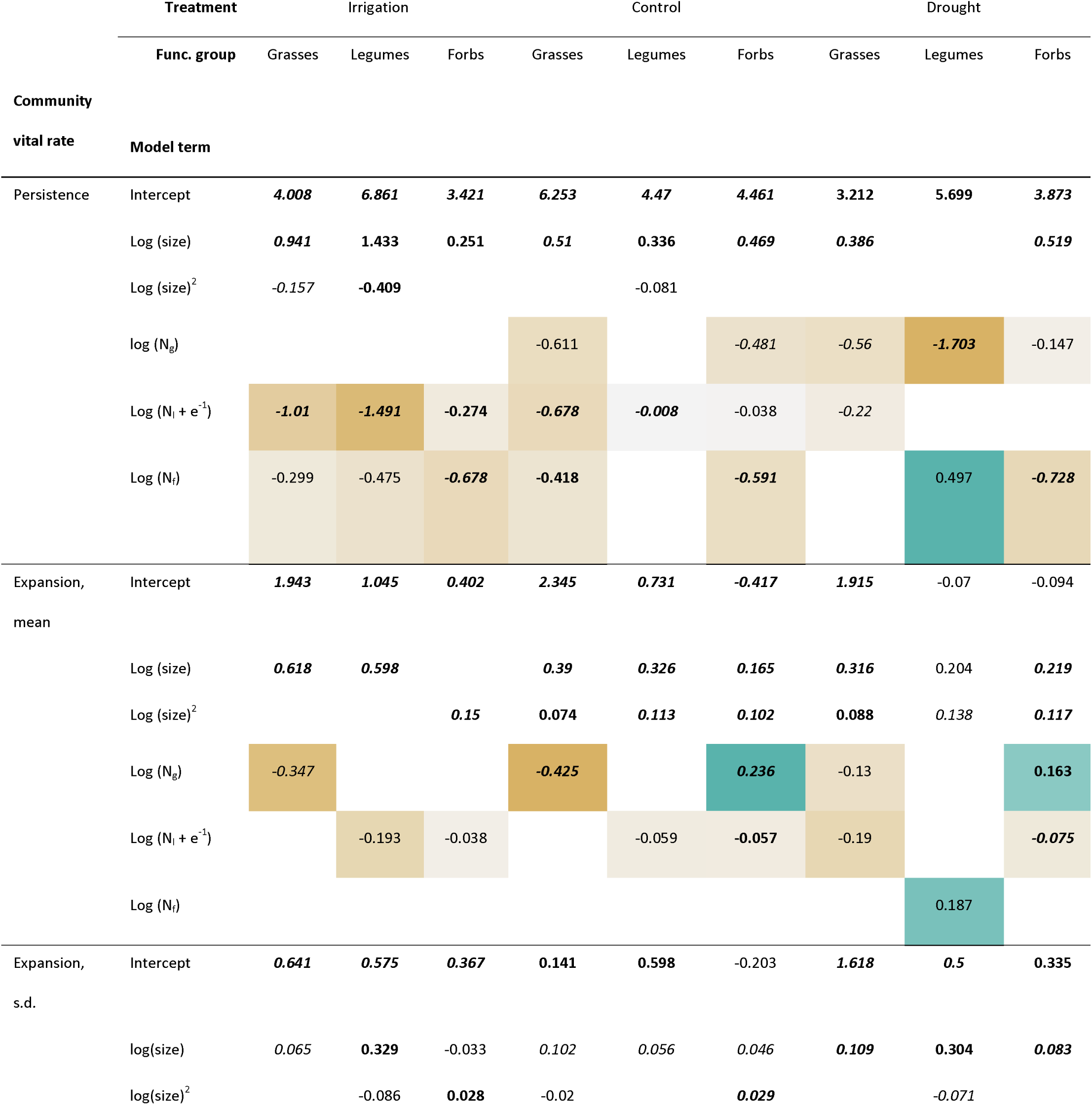
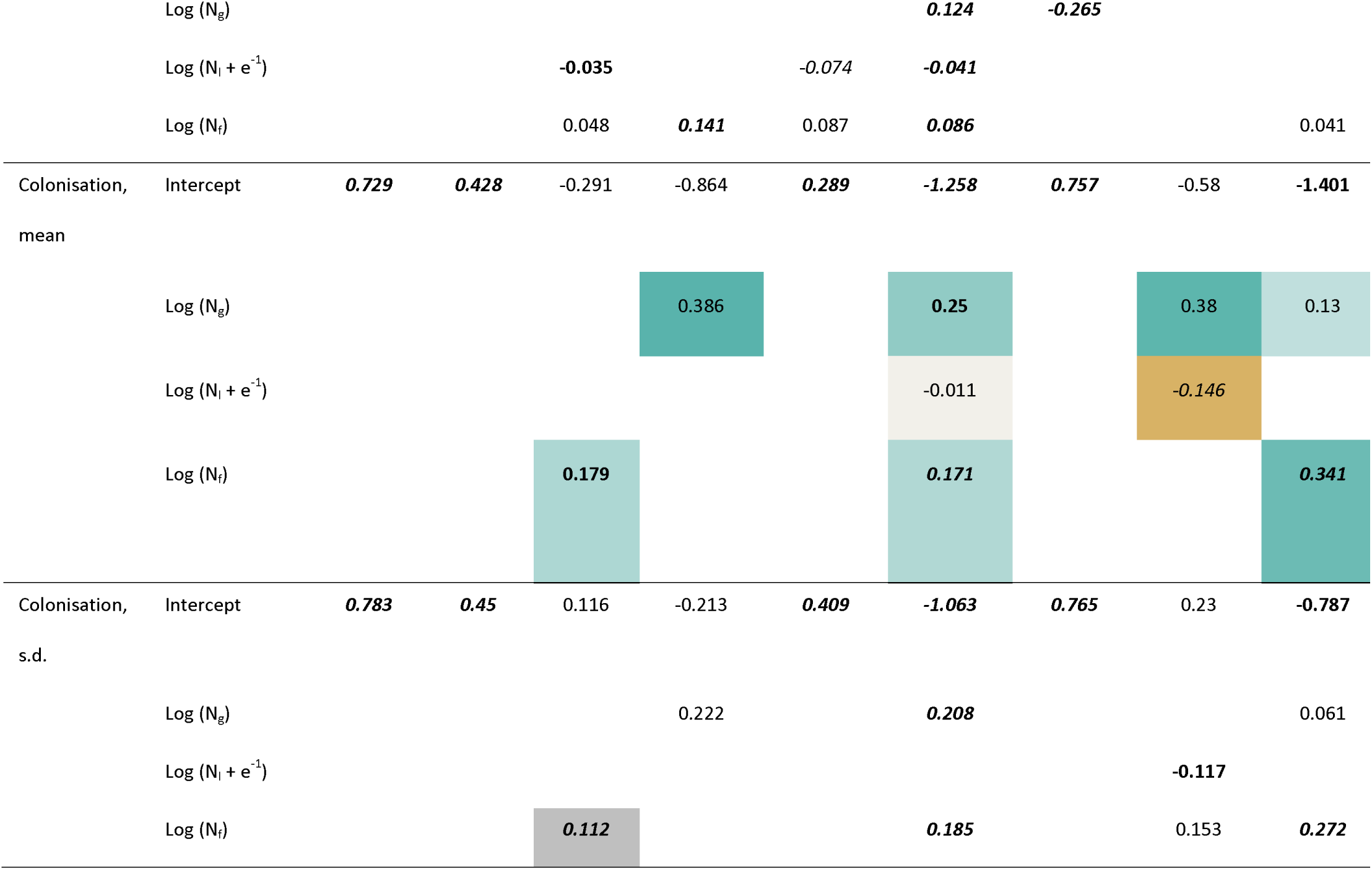
Selected models for persistence, expansion and colonisation of each treatment group functional group. This table demonstrates a shift in inter functional group interactions from competitive to facilitative as precipitation decreases. Cell colours vary from orange (for negative values) to cyan (for positive values) for overall abundance terms *N_k_* to indicate, for a given community vital rate, whether the overall abundance of *k* has a facilitative or competitive effect. The increasing presence of cyan cells further to the right of the table indicates that interactions shift from being primarily competitive to facilitative as water availability decreases (from irrigation to control to drought). Coefficients with p < 0.001 are in bold and italics. Coefficients with p < 0.01 are in bold. Marginally significant coefficients (i.e., with p < 0.1) are in italics.

To test our hypotheses about how our treatments would influence community dynamics (H2.1 and H2.2), we used the above-discussed community models to construct IPMs for each functional group × treatment. The following description of the mathematical structure of IPMs draws from Easterling et al. (2000). Here, our IPMs link the percentage-cover distribution of individual species in a functional group at time *t* to the percentage-cover distribution of individual species at time *t+1*. For a given functional group *k*, the IPM describing the relationship between the number *n_k_ (x, t)* of ‘individuals’ (species here) of cover value *x* at time *t* and the number *n_k_ (y, t + 1)* of species of cover value *y* at time *t + 1*, is

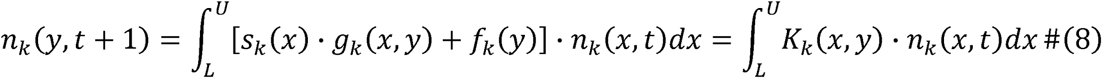

Here, *L =log* 1 = *0* and *U = log* (100) ≈ 4.605 are the lower and upper bounds of the observed percentage cover for an individual species in a functional group, and are sufficiently small and large, respectively, to avoid eviction. The first term *s_k_(x)⋅g_k_ (x, y)* describes the changes in percentage-cover (Fig. 1e) conditional on persistence (Fig. 1c), and the second term *f_k_(y)* quantifies colonisation (Fig. 1f). The resulting *kernel surface K_k_ (x, y) = s_k_(x)⋅g_k_ (x, y) + f_k_(y)* represents all possible transitions for an individual species from percentage-cover *x* at time *t* to percentage-cover *y* at time *t +1*. Here, we discretised each kernel into a 200 × 200 square matrix with a lower-limit *L* =*log* 1 = 0, and an upper limit of *U* = *log* (100) ≈ 4.605. We then generated our kernels using midpoint integration.

To simulate how the influence of each functional group on the other groups changes over time (H 2.1 and H 2.2), all three functional-group IPMs were run for 300 time-steps, to find equilibrium values of *N_g_, N_l_*, and *N_f_* for each treatment. See the supplementary materials for the technical details of how we calculated the percentage-cover stage-structure and overall abundance values of our functional groups at each timestep *t*.

### Pseudospectral analysis

After completing the IPM simulation runs, we tested our hypotheses regarding whether stage-structure influences functional-group stability (H3), and on whether transient behaviours deviate from asymptotic predictions (H4). To test these hypotheses, we performed a pseudospectral analysis (Trefethen & Embree 2005) on the final IPM (at *t* = 300) for each functional group × treatment. To help contextualise the results, below we briefly describe pseudospectra and how they are calculated.

Let *M* be a square matrix, such as an IPM after it has been discretised (Easterling et al. 2010), or a community matrix containing species competition coefficients (Kot 2001). The ɛ-pseudospectrum of *M* is the region consisting of the eigenvalues *M* has when its entries are subjected to noise of magnitude *ɛ*, given by

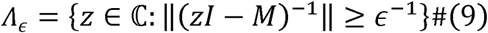

Biologically, the ɛ-pseudospectrum Λ_ɛ_ of *M* measures the extent to which the *response* of a community to some dynamical perturbation (change of cover, in this case) is altered by a structural perturbation of size ɛ to the entries of *M* (Trefethen and Embree 2005). Crucially, for a discrete-time matrix model *M* representing a stable system with all eigenvalues within the unit circle of the complex plane, its ɛ-pseudospectrum illuminates its transient behaviour after being perturbed out of equilibrium. Let ρ_ɛ_ be the maximum absolute value attained by the points in the ɛ-pseudospectrum Λ_ɛ_ of *M*, and let T_ɛ_ = (ρ_ɛ_ - 1) / ɛ. Then we have the following inequality (Trefethen & Embree 2005):

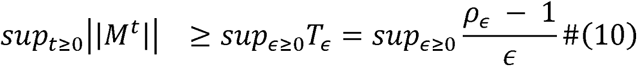

If ρ_ɛ_ < 1 + ɛ, then *T_ɛ_* < 1 for all ɛ, and thus so are all matrix norms ||*M^t^*|| for *t* ≥ 0. But if ρ_ɛ_ > 1 + ɛ, or equivalently if *T_ɛ_* > 1, then although the system is asymptotically stable, it is transiently unstable — *i.e.*, perturbations are initially amplified by the system.

For clarity, we illustrate two examples of pseudospectra in Fig. 3, corresponding to two made-up IPMs. If sup_ɛ≥0_ *T_ɛ_* < 1 for the pseudospectrum of a matrix model (Fig 3a), the ecological system does not exhibit transient instability, with any perturbations immediately decaying (Fig 3b). If sup_ɛ≥0_ *T_ɛ_* > 1 for the pseudospectrum of a matrix model (Fig. 3c), the system exhibits transient instability (Fig 3d). Biologically, this relationship means that perturbations are initially amplified by a community in the short-term, before eventually decaying. For a formal mathematical explanation, see Trefethen & Embree (2005).

**Figure 3.**
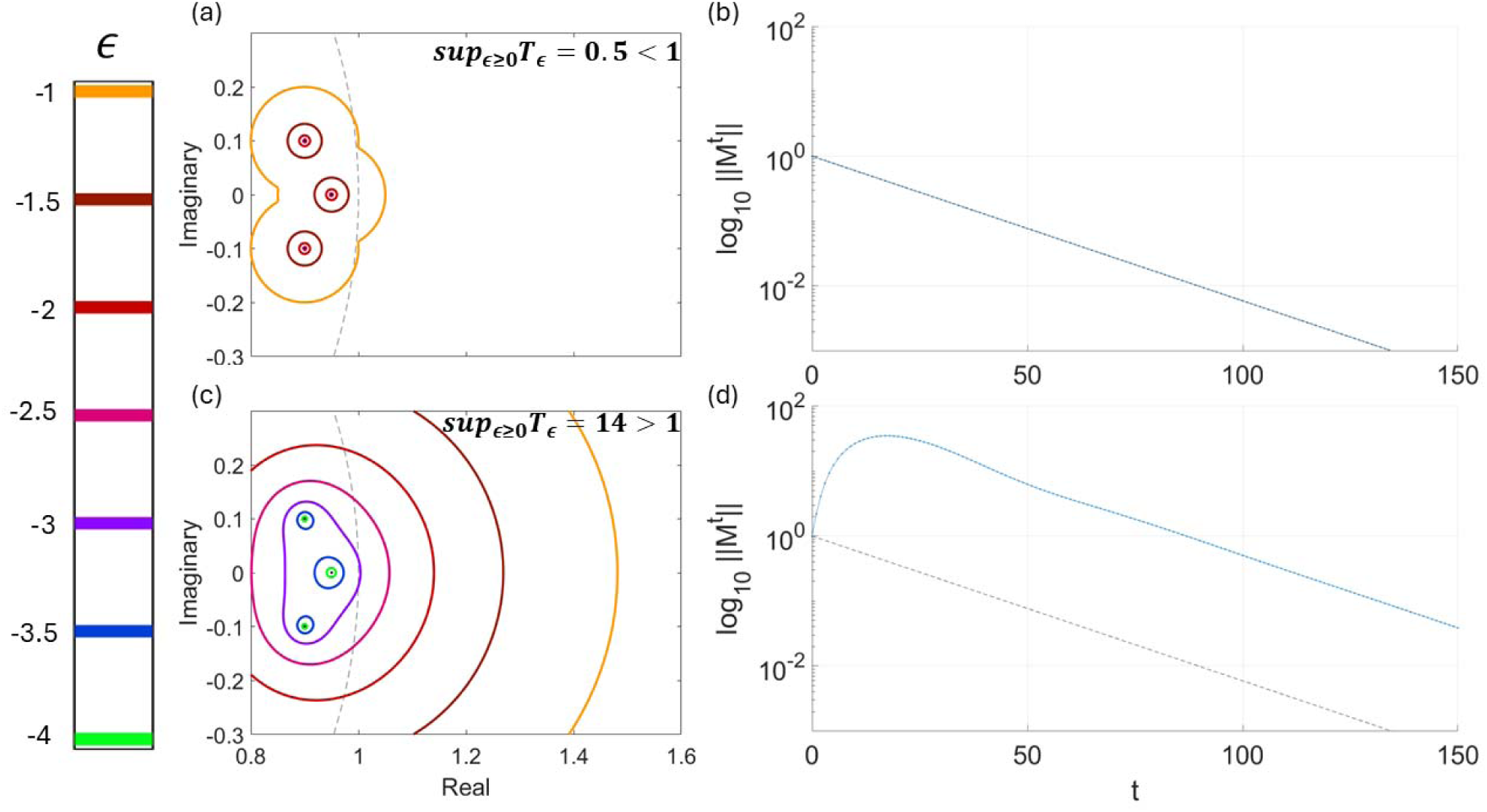
Examples of pseudospectra that do or do not exhibit transient instability. In (a), the dots represent the eigenvalues of a matrix, and the coloured lines are the boundaries of their pseudospectral envelope, representing the spectrum of possible transient rates of expansion or contraction a functional group exhibits. The grey dashed line is the boundary of the unit circle *x*^2^ + *y*^2^ = 1, which represents the threshold between species expansion and contraction. For each coloured pseudospectral envelope, the scale to the left represents the values of ɛ, which is the degree of noise that matrix *M*s entries are subject to due to structural perturbations. (a) The pseudospectrum for a matrix such that sup_ɛ≥0_ *T_ɛ_* < 1. This matrix is therefore both asymptotically and transiently stable. (b) A plot of how perturbations evolve for a transiently and asymptotically stable matrix such as the one in (a). The y axis represents the magnitude ||*M^t^*|| (at log_10_ scale) of a perturbation from a community’s steady state, and the x-axis represents time *t* following an initial perturbation. Following the blue line, any perturbation in this case decays exponentially, back to a steady-state. (c) The pseudospectrum of a matrix such that sup_ɛ≥0_ *T_ɛ_* > 1. Here, the system is transiently unstable, and amplifies any disturbance applied to it in the short-term. But then, the disturbance decays in the long-term and the system returns to equilibrium. (d) A plot of how perturbations evolve for a transiently and asymptotically stable matrix such as the one in (c). Following the blue line, the initial perturbation is initially amplified, before eventually starting to decay once *t* is approximately 20.

Finally, to determine whether stage-structure influences community stability (H3) and whether the community’s transient behaviour deviated from long-term eigenvalue stability predictions (H4), we calculated the pseudospectra for our IPMs. The eigenvalues and -pseudospectra for each IPM (for ɛ = 10^-4^ to 10^-1^) were then calculated using the MATLAB package Eigtools (Wright 2002). Further, to assess the stability of each treatment as a whole, we calculated the eigenvalues and ɛ-pseudospectra for the community matrix of the entire dynamical system represented by the IPMs for each of the three functional groups under each treatment. See the supplementary materials for how we constructed these community matrices. As the norms for all our IPMs were of order 1, the chosen values of all represent ‘small’ structural perturbations to the entries of our IPMs.

To support H3, the eigenvalues for the IPM of any functional group × treatment would have to lie neither completely within nor completely outside the unit circle. If all eigenvalues lie either within or outside the unit circle, then any stage-structure would produce the same outcome, either stability or instability respectively, regardless of the direction of the perturbation applied to the initial stage-structure. Conversely, if some eigenvalues lie within the unit circle but some ‘pull’ the pseudospectrum outside it by being close to its boundary (enabling sup_ɛ≥0_ *T_ɛ_* to be greater than 1), then stage-structure influences stability. Under this latter scenario, some perturbations of the initial stage-structures would be stable: those produced by linear combinations of the eigenvectors corresponding to eigenvalues close to the origin. Yet, other perturbations would be transiently or asymptotically unstable: linear combinations of the eigenvectors corresponding to eigenvalues close to or outside the unit circle (Vindenes et al. 2021). To find support for H4, all eigenvalues of the IPM of any functional group × treatment or any treatment would need to be grouped within the unit circle (as in Fig. 3b), but with sup_ɛ≥0_ *T*_ɛ_ > 1. This arrangement would indicate that the functional group transiently expands, despite being predicted to be stable in the long-term (Trefethen & Embree 2005).

## Results

We found that inter-functional-group interactions were exclusively competitive under the irrigation condition, thus supporting H1.1. Several facilitative interactions arose under the drought condition interspersed with competitive interactions, partially supporting H1.2 (Table 1). Under the drought condition, the overall presence of forbs positively influenced the persistence of legumes, grasses facilitated the expansion of forbs, and forbs facilitated the expansion of legumes. Further, grasses facilitated colonisation by legumes and forbs, and forbs facilitated colonisation by other forbs (Table 1).

Overall, our functional-group IPM simulations supported our hypotheses about how extreme precipitation would alter community outcomes (H2.1., H2.2). Under irrigation, grasses are dominant, with percent cover roughly an order of magnitude larger than forbs and legumes (Fig. 4a), thus supporting H2.1. Further, under drought, after seven timesteps, forbs are the dominant cover rather than grasses, and after 15 timesteps, the overall abundance of legumes also surpasses the overall abundance of grasses (Fig. 4c), thus supporting H2.2. However, although our IPMs correctly predicted the expected overall relative levels of cover of each functional group, they consistently over-predicted the overall abundance attained by each group. The overestimation can be observed in Fig. 4, since the overall abundance values of all functional groups under all treatments are significantly higher at *t* = 300 than at *t* = 0. Overall abundance values at *t* = 0 represent the empirically observed average for each functional group × treatment.

**Figure 4.**
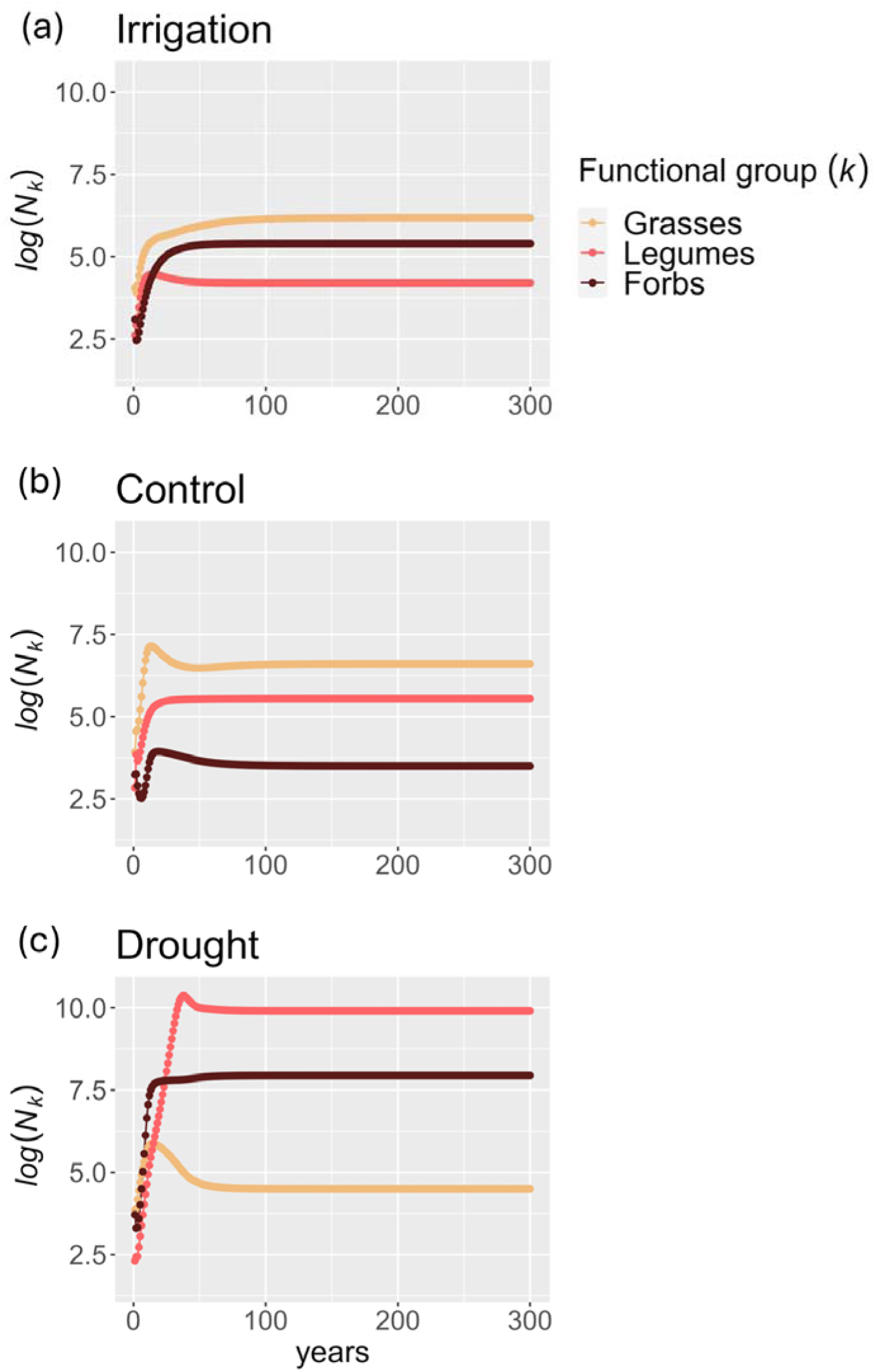
Multi-functional group IPM predictions of functional-group percentage cover for each treatment, for 300 time-steps. All functional groups reach a steady percentage-cover value (*i.e.*, log (*N_k_*) changes by less than 10^-6^) by the last 10 time-steps. (a) Under the irrigation treatment, grasses dominate over (*i.e.*, have a greater overall abundance than) forbs, which in turn dominate over legumes. (b) Under the control treatment, grasses dominate over legumes, which in turn dominate over forbs. (c) Under the drought treatment, Legumes dominate over forbs, which in turn dominate over grasses.

Our pseudospectral analysis supported H3 since, for all functional group × treatments, and for each treatment as a whole, not all eigenvalues clustered inside or outside the unit circle. In all cases, most eigenvalues clustered near the origin of coordinates, within the origin-centred circle of radius 0.7, with only the dominant eigenvalue near the unit circle (Fig. 5a–l). Thus, the pseudospectra emanating from these eigenvalues always crossed the unit circle. These pseudospectra indicate that functional-group transient and asymptotic stability depends on percentage-cover structure. If the linear combination of eigenvectors describing a perturbation from the initial percentage-cover distribution did not include the dominant eigenvector, then that functional group × treatment would be stable. Conversely, the functional group × treatment would be unstable if the linear combination describing a perturbation did include the dominant eigenvector.

**Figure 5.**
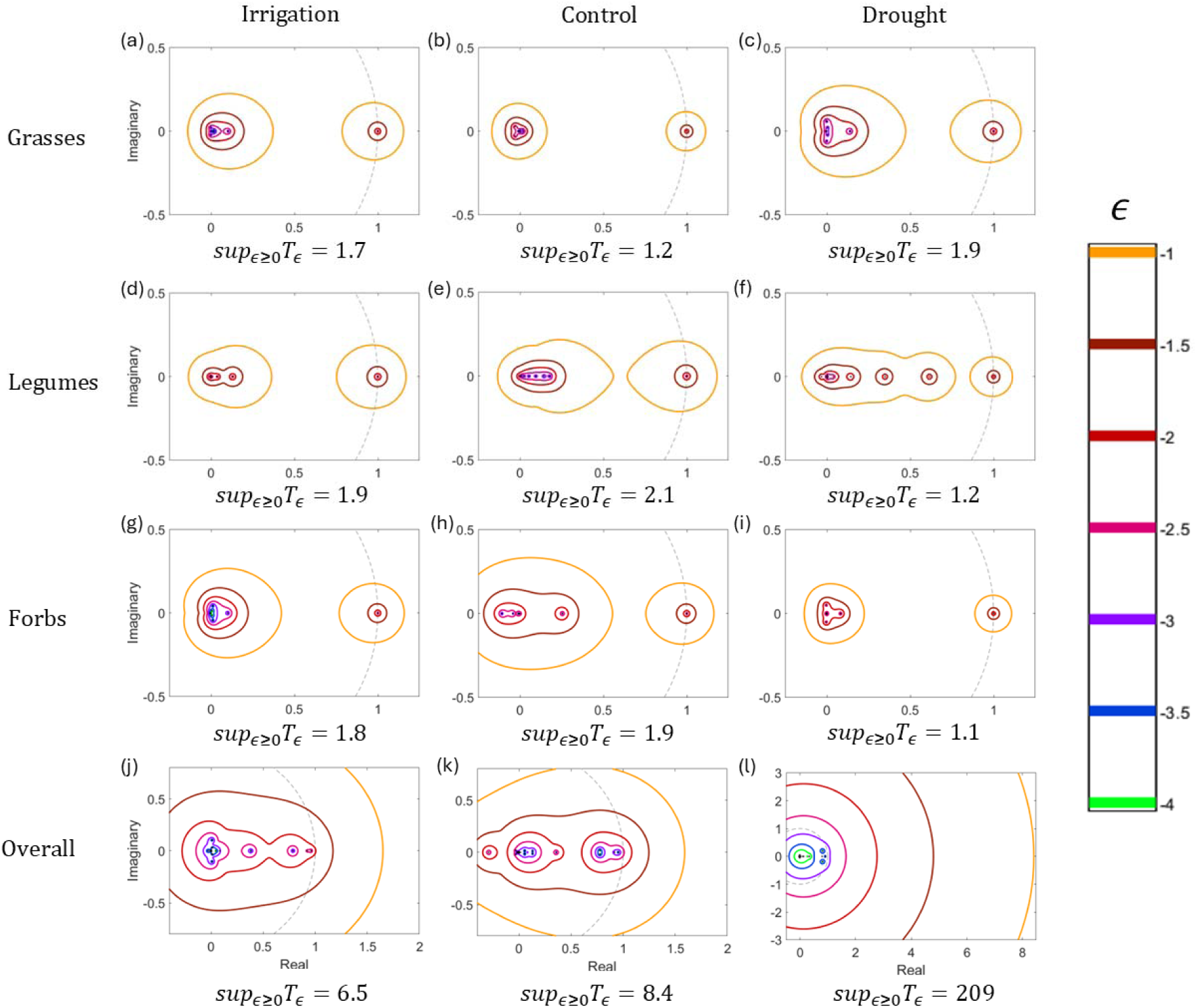
Pseudospectra and eigenvalues of the IPMs produced at the end of the simulation run for each functional group × treatment. For every functional group × treatment (a–i), and every treatment overall (j–l), no eigenvalues lie outside the unit circle. Therefore, they are all stable – *i.e.*, any perturbation is eventually suppressed as *t* → ∞. However, the pseudospectrum for every functional group × treatment, and every treatment overall, is ‘pulled outside’ the unit circle by the dominant eigenvalue, such that sup_ɛ≥0_ *T_ɛ_* > 1. This implies every functional group × treatment is transiently unstable — *i.e.*, they all transiently amplify small perturbations, before perturbations are eventually suppressed. Notably, the degree of transient instability (as given by sup_ɛ≥0_ *T_ɛ_*) for each functional group individually (a–i) are smaller than the degree of transient instability of each treatment as a whole (j–l). To see the values of T_ɛ_ plotted against ɛ for each functional group × treatment, see Figure S1. Under Irrigation (a, d, g) and control (b, e, h), Legumes are more transiently unstable than forbs, which in turn are more transiently unstable than grasses. Under drought (c, f, i), grasses are more transiently unstable than forbs, which in turn are more transiently unstable than legumes.

Our pseudospectral analysis also partially supported H4 by predicting that the transient dynamics for all functional group × treatments differ from asymptotic long-term stability predictions from eigenvalues. From Figure 5, it emerges that sup_ɛ≥0_ *T_ɛ_* > 1 for the pseudospectra of every functional group × treatment. Therefore, the pseudospectrum of every functional group × treatment predicts that it is transiently unstable. However, their eigenvalues predict that these functional groups would be (asymptotically) stable under these treatments in the long-term, since they do not cross beyond the unit circle. Therefore, the prediction of their dynamics given by eigenvalues (indicating asymptotic stability) differs from how their pseudospectra predicts that they would be transiently unstable in the short-term before reaching stability in the long-term.

## Discussion

Using eight years of percentage-cover data on the effects of irrigation and drought on grasslands, we examined how stage-structure shapes the transient dynamics of grassland communities. To do so, we first constructed interlinked functional-group integral projection models (IPMs; Easterling et al. 2000). We then examined these IPMs using pseudospectral theory to study how stage-structure and inter-group interactions shape community outcomes. We found that interactions between functional groups at our field site are almost exclusively competitive under irrigation, while some functional group interactions are facilitative under drought. Then, using these data, we parameterised multi-functional-group IPMs. Our IPM simulations correctly predicted that grasses dominate over legumes and forbs in irrigation conditions, and that legumes and forbs dominate over grasses in drought conditions. Then, by analysing the pseudospectra of these IPMs, we found that the percentage-cover structure of functional groups can determine functional group stability within a community. We also found that, for all functional group × treatments, transient dynamics differ somewhat from long-term predictions stemming from commonly applied eigenvalue analysis (Kot 2001).

Our approach includes two key advances. First, we demonstrated how IPMs can be repurposed to scale up from population to community ecology. We do this by analysing communities at a functional-group level to overcome the excessive computational costs of species-level community analysis. This scaling-up offers a powerful tool to gain mechanistic insight into how community vital rates drive emergent patterns such as disturbance-driven restructuring or range shifts. Obvious candidates for phenomena that this scaling-up can be applied to are those for which IPMs have already been applied, albeit from a demographic rather than community perspective. Examples include forest species colonisation (Zhao et al. 2021, Drees et al. 2023), coral reef responses to extreme weather (Kayal et al. 2019, Creswell et al. 2024), and species coexistence (Garcia-Cervigon et al. 2021, Peirce et al. 2023). Second, the application of pseudospectra theory to analyse the transient stability of IPMs is new, since so far, they have only been deployed for community matrix models (Caravelli and Staniczenko 2016, Kostić et al. 2015). As the usage of IPMs continues to grow across animal and plant populations (Levin et al. 2022), pseudospectra potentially offer insight into how their surrounding communities may respond to perturbations to both species numbers and interaction strengths, and into their transient dynamics in response to these key perturbations (Trefethen and Embree 2005).

Despite these advances, our model consistently over-predicted percentage cover across functional group × conditions. There are at least three possible explanations for these over-predictions: (1) our study site is presently recovering from the effects of long-term agricultural and grazing, (2) our model simulates the effects of inter- and intra-functional-group interactions and does not account for abiotic effects that would inhibit percentage-cover growth, (3) it does not take into explicit account higher-order interactions. For (1), Jackson et al. (2024) showed that at our study site, there was a treatment-independent increase in species richness with time. Therefore, the models that best fitted our data may implicitly reflect how our grassland communities would develop if the secondary succession due to their recovery continues ‘forever,’ rather than eventually ceasing. For (2), potential abiotic inhibitors include limitations of space or nutrients (Craine and Dybzinski 2013, Morris et al. 2019). Thus, without these additional limiting factors, our simulated functional groups may have been able to grow larger than their real-life counterparts before ‘purely’ community factors constrained them (Senthilnathan 2023). For (3), higher-order interactions, whereby the interaction of two species is mediated by a third, their inclusion in community models produces more realistic predictions of species abundance than models with only linear, additive interaction terms (Mayfield and Stouffer 2017). Hence, future studies can further build on our framework to disentangle community processes by (1) implementing it for a range of ecosystems at differing levels of succession and stability, (2) the inclusion of terms reflecting abiotic constraints and (3) the inclusion of higher-order interactions into our community vital rate models.

Pseudospectra offer a valuable complement to eigenvalues as a means of assessing ecosystem stability, by informing us about their transient behaviour in addition to their asymptotic stability. Indeed, eigenvalue analysis alone has become common practice in community ecology (*e.g.* by evaluating the dominant eigenvalue of Jacobian matrices; Kot 2001). Revealing how systems respond immediately to disturbances uncovers crucial features of their dynamics that cannot be observed in asymptotic, eigenvalue-based predictions. Admittedly, the individual functional-group IPMs had a limited degree of transient instability: compare, for example, the values of sup_ɛ≥0_ *T_ɛ_* across Fig 5a–i to the matrix in Fig 3a. By comparison, the values of sup_ɛ≥0_ *T_ɛ_* are much larger for the overall community matrices Fig 5j–l. This potentially indicates that individual functional-group transient instabilities can amplify each other transiently to produce a much greater transient instability in the whole community. However, it is also possible that the structural perturbations that lead to such large transient instabilities in the whole community matrices are ecologically unrealistic, and further work should try to distinguish unrealistic and realistic perturbations. Nevertheless, if a system is exposed to a perturbation and is shown (via pseudospectra) to exhibit transient instability, further perturbations may possibly drive it away from its initial steady state to a new one (Caravelli and Staniczenko 2016). An eigenvector analysis, on the other hand, would predict that such a system, being asymptotically stable, remains stable no matter how many perturbations it experiences (Caravelli and Staniczenko 2016). Thus, a pseudospectral analysis could potentially reveal important features of the transient dynamics of an ecological system that may drive outcomes that differ significantly from asymptotic stability predictions. Further, pseudospectral theory enables community stability analysis using only crude, order-of-magnitude estimates of interaction strengths, thus widening the range of data for which stability can be assessed (Barabás & Allesina 2015).

We found broad support for the stress gradient hypothesis (Bertness & Callaway 1994). In our experiment, where we manipulated the key resource in this grassland, water availability, we found that this hypothesis holds not only for interspecific interactions, but also inter-functional-group interactions. Specifically, under our irrigation treatment, functional groups tend to suppress each other’s expansion, colonisation, and persistence. On the other hand, under our drought treatment, some functional groups facilitate each other, although competitive interactions are still present. This mix of competitive and facilitative interactions under drought suggests that the stress gradient between irrigation, control and drought treatments is not strong. Under a weak stress gradient, due to weaker stress amelioration and niche differences within a community, the shift to facilitation under stress tends to be less pronounced (Qi et al. 2018). Additionally, calcareous grassland communities tend to be relatively stress-tolerant (Basto et al. 2018), further inhibiting the facilitatory shift. Nevertheless, these results do corroborate empirical studies suggesting inter-functional-group interactions shift from competitive to (partially) facilitative as water availability falls (Grant et al. 2014, Lima et al. 2022). Further, our predictions of relative levels of functional group abundance agree with prior observations of grasslands under extreme precipitation (Hallet et al. 2019).

Our findings are also pertinent to recent theoretical work indicating that the population structure of species within a community can significantly influence their stability (de Roos 2020, 2021, Muehleisen et al. 2022). Whilst these works demonstrate that stage-structured populations support much more complex, stable interaction webs than unstructured ones, we showed by contrast that for our system, stage-structure drives a limited degree of transient instability. The more transiently unstable a functional group is, the more it will either grow or contract in response to a disturbance, and the greater the magnitude of its interaction-effects with other functional groups will be (Grant et al. 2014, Shriver et al. 2019).

Increasingly, it is recognised that instability is the norm, not the exception, in ecological systems (Morozov et al. 2020, Capdevila et al. 2021). The pseudospectra we generated from our field-parameterised models corroborate that claim somewhat, since all functional group models under all treatments we examined here were transiently unstable. Although the degree of transient instability for individual functional groups was limited, our studied communities showed a greater degree of transient instability for each treatment. Hence, our results provide a clear motivation to examine transient instability more broadly, since it has the potential of significantly shaping community outcomes. This motivation is especially clear given that climate change is expected to further drive instability in ecosystems worldwide (Pecl et al. 2017). Therefore, there is a growing call for treating ecosystems as ‘dynamic regimes’ rather than ‘equilibrium regimes’ (Coulson 2020, Sánchez-Pinillos 2023, 2024). However, in work addressing this call, there has been a persistent divide between theory and empirical studies or conservation practice (Dakos and Kéfi 2022, Oro & Martínez-Abraín 2023). Perhaps the most prominent manifestation of this divide is a lack of model-validation studies, which significantly limits our understanding of both model validity and practical utility to *e.g.* conservationists (Schuwirth et al. 2019, Hale et al. 2023). Our study bridges this divide by using models parameterised by field data (rather than simulated communities) to construct IPMs on which to perform pseudospectral analysis. We encourage future studies to continue to bridge this divide, especially by validating models against real-world data. In an increasingly unstable and dynamic biosphere, the tools used to assess ecosystems must reflect this instability to ensure both accuracy of our mechanistic understanding and effectiveness of conservation practice (Sánchez-Pinillos et al. 2023).

## Supporting information

SOM

## Acknowledgements

We thank B. Blonder for feedback in earlier versions of this manuscript. We also like to thank N. Fisher, N. Havercroft, and K. Crawford for field logistic support at Wytham throughout the study, as well as M. Stone and D. Gowing for their work setting up and supporting the experiment, and to C. Lawson for her role in the data collection, alongside J. Jackson, E. Jardine, N. Hawes, and K. Maseyk. A. Hector was supported by the John Fell Fund. This work was funded by the NERC Research Experience Program (REP) grant to RSG to host AG, and a NERC Pushing the Frontiers grant to RSG (NE/X013766/1).

