## Supplementary material for "Variation in precipitation drives differences in interactions and short-term transient instability between grassland functional groups: a stage-structured community approach": SOM

### **Details of the study site**

Our site is a calcareous chalk grassland, with alkaline soil of shallow depth (300mm-500mm). According to the Köppen-Geiger climate classification, the site has a temperate maritime climate (Cfb) (Peele et al., 2007). During the years when our data were collected, 2016-2023, the mean annual temperature on the site was 11.5 ℃, with a daily temperature range between -5 ℃ and 26 ℃. The annual total precipitation was 686 mm, with a daily precipitation range of between 0-40 mm (Fenollosa et al. 2024). The study site is mowed twice a year, first mid-growing season (late July), and then at the end of the growing season (late September), as per European grassland management regulations (Török et al., 2018). Further details of our study site are found in Fenollosa et al. (2024) and Jackson et al. (2024).

### **Experimental setup**

In the drought plots, a rainwater shelter syphons off 50% of the naturally occurring precipitation via gulleys into storage containers by the side of the plot (Fig. 2b). The water in these containers is then sprayed onto the irrigation plots using sprinklers that are connected to the containers by pipes and a pump system (Fig. 2b). This design ensures that the changes in precipitation for the experimental plots are proportional to background levels of precipitation at the site. The ambient control had no structure and no experimental treatment (Fig. 2b). For the procedural control, shelters with inverted gullies are used (which let 100% precipitation fall through; Fig. 2b) to evaluate whether the shelter structure has any effect on the plot beyond the intended reduction in precipitation. Potential confounding effects included changes in boundary layer dynamics that could influence vegetation via changes in temperature, humidity or light availability.

### **Model selection**

Tables S1 – S45 provide the AICc values for each possible combination of log-transformed linear or quadratic relationships of percentage-cover, and overall cover of grasses (*N_g_*), legumes (*N_l_*), and forbs (*N_f_*) to include in our survival, persistence and expansion models for each functional group × treatment. To select from these models, the models with the least AICc value were first chosen, and IPMs were run using them. If they produced biologically unrealistic results (*e.g.* unbounded growth, or standard deviation eventually becoming negative), models with the next lowest AICc value was chosen, and this process of iteration was repeated until a suite of models was found which produced realistic results. For each table, the model that was selected through this process is indicated in bold.

#### **Table S1 – Model selection for grasses under irrigation, persistence**

| (Intercept) | I((log(size))^2) | log(Nf) | log(Ng) | log(Nl + exp(-1)) | log(size) | df | logLik | AICc | delta | weight |
| --- | --- | --- | --- | --- | --- | --- | --- | --- | --- | --- |
| 3.124855 | -0.14649 | NA | NA | -1.01695 | 0.872829 | 4 | -147.73 | 303.6108 | 0 | 0.208568 |
| **4.007948** | **-0.15646** | **-0.29884** | **NA** | **-1.01033** | **0.94093** | **5** | **-146.787** | **303.8001** | **0.189364** | **0.189726** |
| 1.870752 | -0.17499 | -0.39524 | 0.584034 | -0.99731 | 0.976156 | 6 | -145.995 | 304.3071 | 0.696363 | 0.147243 |
| 3.155395 | NA | NA | NA | -0.99321 | 0.450329 | 3 | -149.277 | 304.6437 | 1.032923 | 0.124438 |
| 4.055239 | NA | -0.29447 | NA | -0.99102 | 0.485162 | 4 | -148.344 | 304.8382 | 1.227449 | 0.112904 |
| 1.712361 | -0.1554 | NA | 0.337602 | -1.00776 | 0.883073 | 5 | -147.443 | 305.1115 | 1.500691 | 0.098487 |
| 2.493525 | NA | -0.36573 | 0.426015 | -0.97733 | 0.473792 | 5 | -147.9 | 306.0248 | 2.414031 | 0.06238 |
| 2.316088 | NA | NA | 0.199338 | -0.98548 | 0.442224 | 4 | -149.171 | 306.492 | 2.881272 | 0.049384 |
| 3.312265 | 0.089516 | NA | NA | -0.94953 | NA | 3 | -153.191 | 312.4723 | 8.861516 | 0.002483 |
| 4.020297 | 0.097706 | -0.23127 | NA | -0.94522 | NA | 4 | -152.602 | 313.353 | 9.74226 | 0.001599 |
| 2.352452 | 0.085087 | NA | 0.229015 | -0.94159 | NA | 4 | -153.049 | 314.2473 | 10.63654 | 0.001022 |
| 2.50822 | 0.091349 | -0.29875 | 0.412296 | -0.93176 | NA | 5 | -152.177 | 314.5786 | 10.96782 | 0.000866 |
| 3.546044 | NA | NA | NA | -0.94043 | NA | 2 | -156.148 | 316.3413 | 12.73053 | 0.000359 |
| 1.714954 | NA | NA | 0.433822 | -0.92911 | NA | 3 | -155.58 | 317.2493 | 13.63852 | 0.000228 |
| 3.968818 | NA | -0.13565 | NA | -0.93612 | NA | 3 | -155.93 | 317.9491 | 14.33837 | 0.000161 |
| 1.798781 | NA | -0.24937 | 0.600743 | -0.91992 | NA | 4 | -154.956 | 318.0623 | 14.45148 | 0.000152 |
| 1.243436 | -0.13395 | -0.37221 | NA | NA | 0.847412 | 4 | -163.005 | 334.1588 | 30.54807 | 4.85E-08 |
| 1.285848 | NA | -0.34814 | NA | NA | 0.445832 | 3 | -164.109 | 334.3084 | 30.69759 | 4.50E-08 |
| -0.232 | -0.14593 | -0.44311 | 0.414617 | NA | 0.867751 | 5 | -162.402 | 335.0293 | 31.41849 | 3.14E-08 |
| 0.219791 | NA | NA | NA | NA | 0.404852 | 2 | -165.589 | 335.222 | 31.61127 | 2.85E-08 |
| 0.130424 | -0.11259 | NA | NA | NA | 0.736326 | 3 | -164.655 | 335.4002 | 31.78945 | 2.61E-08 |
| 0.097767 | NA | -0.40475 | 0.334902 | NA | 0.43512 | 4 | -163.704 | 335.5578 | 31.94708 | 2.41E-08 |
| -0.33444 | NA | NA | 0.135987 | NA | 0.399203 | 3 | -165.517 | 337.1227 | 33.51192 | 1.10E-08 |
| -0.66803 | -0.11667 | NA | 0.19562 | NA | 0.73909 | 4 | -164.511 | 337.1709 | 33.56015 | 1.08E-08 |
| 1.371903 | 0.097456 | -0.29223 | NA | NA | NA | 3 | -167.611 | 341.3117 | 37.70092 | 1.36E-09 |
| 0.4606 | 0.088197 | NA | NA | NA | NA | 2 | -168.694 | 341.4328 | 37.82208 | 1.28E-09 |
| 0.243362 | 0.092634 | -0.34729 | 0.319333 | NA | NA | 4 | -167.236 | 342.622 | 39.01122 | 7.05E-10 |
| -0.11771 | 0.085656 | NA | 0.14183 | NA | NA | 3 | -168.614 | 343.3173 | 39.70654 | 4.98E-10 |
| 0.72085 | NA | NA | NA | NA | NA | 1 | -171.954 | 345.9219 | 42.31116 | 1.35E-10 |
| 1.397429 | NA | -0.21111 | NA | NA | NA | 2 | -171.343 | 346.7308 | 43.12 | 9.03E-11 |
| -0.23255 | NA | -0.30231 | 0.465369 | NA | NA | 3 | -170.503 | 347.0953 | 43.48452 | 7.53E-11 |
| -0.48767 | NA | NA | 0.292534 | NA | NA | 2 | -171.584 | 347.2133 | 43.60256 | 7.10E-11 |

#### **Table S2 – Model selection for grasses under irrigation, expansion, mean**

| (Intercept) | I((log(size))^2) | log(Nf) | log(Ng) | log(Nl + exp(-1)) | log(size) | df | logLik | AICc | delta | weight |
| --- | --- | --- | --- | --- | --- | --- | --- | --- | --- | --- |
| **1.943046** | **NA** | **NA** | **-0.3471** | **NA** | **0.616822** | **4** | **-242.726** | **493.6763** | **0** | **0.186668** |
| 2.011259 | 0.042232 | NA | -0.34986 | NA | 0.477404 | 5 | -242.093 | 494.525 | 0.848705 | 0.122117 |
| 1.845046 | NA | 0.116299 | -0.41172 | NA | 0.616254 | 5 | -242.098 | 494.5354 | 0.859089 | 0.121485 |
| 1.907469 | 0.050251 | 0.13854 | -0.42737 | NA | 0.450253 | 6 | -241.218 | 494.9134 | 1.237066 | 0.100565 |
| 0.52733 | NA | NA | NA | NA | 0.600047 | 3 | -244.653 | 495.4408 | 1.76445 | 0.077255 |
| 1.8961 | NA | NA | -0.36519 | 0.044992 | 0.61809 | 5 | -242.554 | 495.4478 | 1.771473 | 0.076984 |
| 1.956237 | 0.04672 | NA | -0.37416 | 0.05968 | 0.464267 | 6 | -241.796 | 496.0699 | 2.393626 | 0.056403 |
| 0.582871 | 0.041186 | NA | NA | NA | 0.463951 | 4 | -244.064 | 496.3529 | 2.676572 | 0.048962 |
| 1.811299 | NA | 0.111186 | -0.42355 | 0.036471 | 0.617307 | 6 | -241.986 | 496.4494 | 2.773095 | 0.046655 |
| 1.864031 | 0.053813 | 0.132864 | -0.44525 | 0.051728 | 0.439979 | 7 | -240.995 | 496.6292 | 2.952881 | 0.042644 |
| 0.390223 | NA | 0.044099 | NA | NA | 0.598647 | 4 | -244.556 | 497.3362 | 3.659866 | 0.029946 |
| 0.479663 | NA | NA | NA | 0.017761 | 0.600202 | 4 | -244.626 | 497.4775 | 3.80121 | 0.027903 |
| 0.396572 | 0.044638 | 0.061419 | NA | NA | 0.450593 | 5 | -243.878 | 498.0941 | 4.417835 | 0.0205 |
| 0.503274 | 0.043465 | NA | NA | 0.030804 | 0.456688 | 5 | -243.984 | 498.3076 | 4.631292 | 0.018425 |
| 0.3634 | NA | 0.041547 | NA | 0.012951 | 0.598841 | 5 | -244.542 | 499.4224 | 5.746107 | 0.010552 |
| 0.344968 | 0.046248 | 0.057112 | NA | 0.025027 | 0.445629 | 6 | -243.826 | 500.1288 | 6.452449 | 0.007412 |
| 2.021431 | 0.162438 | 0.189428 | -0.42801 | NA | NA | 5 | -246.423 | 503.1842 | 9.507889 | 0.001609 |
| 2.176168 | 0.160684 | NA | -0.31968 | NA | NA | 4 | -247.997 | 504.2186 | 10.54228 | 0.000959 |
| 1.951064 | 0.163969 | 0.178979 | -0.45532 | 0.079062 | NA | 6 | -245.923 | 504.3224 | 10.64605 | 0.000911 |
| 2.084298 | 0.16258 | NA | -0.35844 | 0.092057 | NA | 5 | -247.323 | 504.9859 | 11.30956 | 0.000653 |
| 0.863546 | 0.156668 | NA | NA | NA | NA | 3 | -249.547 | 505.2284 | 11.55205 | 0.000579 |
| 0.508355 | 0.156901 | 0.112231 | NA | NA | NA | 4 | -248.945 | 506.1138 | 12.43745 | 0.000372 |
| 0.689405 | 0.157646 | NA | NA | 0.063868 | NA | 4 | -249.218 | 506.661 | 12.98464 | 0.000283 |
| 0.398347 | 0.157678 | 0.102097 | NA | 0.052109 | NA | 5 | -248.729 | 507.7972 | 14.12092 | 0.00016 |
| 1.437759 | NA | NA | NA | NA | NA | 2 | -286.408 | 576.8833 | 83.20698 | 1.60E-19 |
| 1.123323 | NA | 0.099593 | NA | NA | NA | 3 | -286.092 | 578.3172 | 84.64089 | 7.79E-20 |
| 1.754381 | NA | NA | -0.07626 | NA | NA | 3 | -286.349 | 578.8315 | 85.15516 | 6.02E-20 |
| 1.41447 | NA | NA | NA | 0.008721 | NA | 3 | -286.404 | 578.9425 | 85.26616 | 5.70E-20 |
| 1.648514 | NA | 0.125857 | -0.14646 | NA | NA | 4 | -285.893 | 580.0103 | 86.33403 | 3.34E-20 |
| 1.129119 | NA | 0.100146 | NA | -0.00282 | NA | 4 | -286.091 | 580.407 | 86.73069 | 2.74E-20 |
| 1.738836 | NA | NA | -0.08202 | 0.014776 | NA | 4 | -286.337 | 580.8991 | 87.22284 | 2.14E-20 |
| 1.643626 | NA | 0.125127 | -0.14809 | 0.00523 | NA | 5 | -285.891 | 582.1217 | 88.44543 | 1.16E-20 |

#### **Table S3 – Model selection for grasses under irrigation, expansion, s.d.**

| (Intercept) | I((log(size))^2) | log(Nf) | log(Ng) | log(Nl + exp(-1)) | log(size) | df | logLik | AICc | delta | weight |
| --- | --- | --- | --- | --- | --- | --- | --- | --- | --- | --- |
| **0.640883** | **NA** | **NA** | **NA** | **NA** | **0.064674** | **3** | **-142.912** | **291.9577** | **0** | **0.124982** |
| 0.677099 | 0.016892 | NA | NA | NA | NA | 3 | -143.088 | 292.31 | 0.35231 | 0.104796 |
| 0.518499 | NA | NA | NA | 0.045602 | 0.065074 | 4 | -142.37 | 292.9655 | 1.007757 | 0.075512 |
| 0.53858 | 0.01767 | NA | NA | 0.050803 | NA | 4 | -142.42 | 293.0654 | 1.107642 | 0.071833 |
| 0.739011 | NA | NA | NA | NA | NA | 2 | -144.762 | 293.5903 | 1.632522 | 0.055252 |
| 0.763783 | NA | NA | -0.03013 | NA | 0.066131 | 4 | -142.868 | 293.961 | 2.003217 | 0.045904 |
| 0.646916 | 0.004473 | NA | NA | NA | 0.049892 | 4 | -142.891 | 294.0062 | 2.048502 | 0.044877 |
| 0.620057 | NA | 0.006698 | NA | NA | 0.064462 | 4 | -142.905 | 294.0347 | 2.076954 | 0.044243 |
| 0.788794 | 0.017234 | NA | -0.0272 | NA | NA | 4 | -143.052 | 294.3294 | 2.371679 | 0.038181 |
| 0.632675 | 0.016921 | 0.014037 | NA | NA | NA | 4 | -143.058 | 294.3405 | 2.382787 | 0.037969 |
| 0.61985 | NA | NA | NA | 0.044622 | NA | 3 | -144.254 | 294.6416 | 2.683902 | 0.032662 |
| 0.712313 | NA | NA | -0.04997 | 0.049328 | 0.067521 | 5 | -142.253 | 294.8451 | 2.887323 | 0.029503 |
| 0.734149 | 0.018362 | NA | -0.05025 | 0.054756 | NA | 5 | -142.302 | 294.9426 | 2.984889 | 0.028099 |
| 0.522859 | 0.008026 | NA | NA | 0.048011 | 0.038573 | 5 | -142.304 | 294.9463 | 2.988566 | 0.028047 |
| 0.525051 | NA | -0.00234 | NA | 0.045873 | 0.06515 | 5 | -142.37 | 295.0781 | 3.120378 | 0.026258 |
| 0.526457 | 0.017671 | 0.004253 | NA | 0.050314 | NA | 5 | -142.418 | 295.1742 | 3.216469 | 0.025026 |
| 0.698997 | NA | 0.012674 | NA | NA | NA | 3 | -144.738 | 295.6095 | 3.651769 | 0.020131 |
| 0.743556 | NA | NA | -0.00109 | NA | NA | 3 | -144.762 | 295.6576 | 3.699818 | 0.019654 |
| 0.752596 | NA | 0.013276 | -0.03751 | NA | 0.066066 | 5 | -142.844 | 296.0267 | 4.068927 | 0.016341 |
| 0.771156 | 0.004564 | NA | -0.03043 | NA | 0.051062 | 5 | -142.846 | 296.0313 | 4.073599 | 0.016303 |
| 0.620763 | 0.004958 | 0.008622 | NA | NA | 0.048017 | 5 | -142.88 | 296.0983 | 4.140577 | 0.015766 |
| 0.771541 | 0.017429 | 0.021122 | -0.03928 | NA | NA | 5 | -142.991 | 296.3214 | 4.3637 | 0.014102 |
| 0.695133 | NA | NA | -0.01904 | 0.046028 | NA | 4 | -144.237 | 296.6982 | 4.740432 | 0.011681 |
| 0.608356 | NA | 0.004034 | NA | 0.044157 | NA | 4 | -144.251 | 296.7275 | 4.769748 | 0.011511 |
| 0.723221 | 0.008475 | NA | -0.0516 | 0.051993 | 0.039618 | 6 | -142.179 | 296.8346 | 4.876864 | 0.010911 |
| 0.707409 | NA | 0.00643 | -0.05334 | 0.048836 | 0.067476 | 6 | -142.247 | 296.972 | 5.014272 | 0.010186 |
| 0.723713 | 0.018471 | 0.01402 | -0.05784 | 0.053738 | NA | 6 | -142.275 | 297.0271 | 5.06941 | 0.009909 |
| 0.521845 | 0.008044 | 0.000366 | NA | 0.047974 | 0.038502 | 6 | -142.304 | 297.0846 | 5.126817 | 0.009629 |
| 0.731527 | NA | 0.014301 | -0.00907 | NA | NA | 4 | -144.734 | 297.693 | 5.735269 | 0.007103 |
| 0.759395 | 0.005473 | 0.015698 | -0.03921 | NA | 0.047986 | 6 | -142.813 | 298.1035 | 6.145719 | 0.005785 |
| 0.689081 | NA | 0.007953 | -0.02324 | 0.045421 | NA | 5 | -144.228 | 298.7954 | 6.837713 | 0.004093 |
| 0.71624 | 0.009012 | 0.01006 | -0.05698 | 0.051391 | 0.037779 | 7 | -142.165 | 298.9701 | 7.012402 | 0.003751 |

#### **Table S4 – Model selection for grasses under irrigation, colonisation, mean**

| (Intercept) | log(Nf) | log(Ng) | log(Nl + exp(-1)) | df | logLik | AICc | delta | weight |
| --- | --- | --- | --- | --- | --- | --- | --- | --- |
| -2.08258 | NA | 0.675944 | NA | 3 | -76.9644 | 160.4186 | 0 | 0.277519 |
| **0.729227** | **NA** | **NA** | **NA** | **2** | **-78.1397** | **160.5193** | **0.100766** | **0.263883** |
| 0.269965 | 0.146273 | NA | NA | 3 | -77.848 | 162.1859 | 1.767326 | 0.114689 |
| 0.366744 | NA | NA | 0.118967 | 3 | -77.9529 | 162.3955 | 1.976971 | 0.103276 |
| -2.12797 | NA | 0.647679 | 0.053488 | 4 | -76.9271 | 162.6876 | 2.269023 | 0.089244 |
| -2.09094 | -0.00459 | 0.681418 | NA | 4 | -76.9642 | 162.7617 | 2.343098 | 0.085999 |
| 0.073947 | 0.124501 | NA | 0.086769 | 4 | -77.7543 | 164.3419 | 3.923299 | 0.039026 |
| -2.1566 | -0.01471 | 0.664094 | 0.055633 | 5 | -76.9249 | 165.1265 | 4.707884 | 0.026363 |

#### **Table S5 – Model selection for grasses under irrigation, colonisation, s.d.**

| (Intercept) | log(Nf) | log(Ng) | log(Nl + exp(-1)) | df | logLik | AICc | delta | weight |
| --- | --- | --- | --- | --- | --- | --- | --- | --- |
| **0.782758** | **NA** | **NA** | **NA** | **2** | **-57.0701** | **118.3802** | **0** | **0.426444** |
| 0.886922 | -0.03318 | NA | NA | 3 | -57.0371 | 120.5639 | 2.183675 | 0.143114 |
| 0.570663 | NA | 0.050987 | NA | 3 | -57.0556 | 120.601 | 2.220817 | 0.140481 |
| 0.840977 | NA | NA | -0.01911 | 3 | -57.0595 | 120.6088 | 2.228524 | 0.139941 |
| 0.460595 | -0.06042 | 0.123049 | NA | 4 | -56.9748 | 122.783 | 4.402732 | 0.047187 |
| 0.912355 | -0.03035 | NA | -0.01126 | 4 | -57.0336 | 122.9005 | 4.520299 | 0.044493 |
| 0.592415 | NA | 0.064532 | -0.02563 | 4 | -57.0375 | 122.9083 | 4.528096 | 0.04432 |
| 0.480989 | -0.05727 | 0.12843 | -0.01728 | 5 | -56.9668 | 125.2101 | 6.829907 | 0.014021 |

#### **Table S6 – Model selection for legumes under irrigation, persistence**

| (Intercept) | I((log(size))^2) | log(Nf) | log(Ng) | log(Nl + exp(-1)) | log(size) | df | logLik | AICc | delta | weight |
| --- | --- | --- | --- | --- | --- | --- | --- | --- | --- | --- |
| 5.487029 | -0.38151 | NA | NA | -1.50491 | 1.297155 | 4 | -60.4016 | 129.111 | 0 | 0.352733 |
| **6.860951** | **-0.40944** | **-0.47447** | **NA** | **-1.49085** | **1.433169** | **5** | **-59.4581** | **129.3814** | **0.270397** | **0.308127** |
| 3.950385 | -0.4068 | -0.65394 | 0.849021 | -1.50254 | 1.438615 | 6 | -58.9089 | 130.4741 | 1.363157 | 0.178419 |
| 4.31167 | -0.3784 | NA | 0.282852 | -1.50224 | 1.286641 | 5 | -60.3301 | 131.1253 | 2.014357 | 0.128835 |
| 6.913465 | NA | -0.49561 | NA | -1.44702 | 0.33916 | 4 | -64.611 | 137.5296 | 8.418668 | 0.00524 |
| 5.369037 | NA | NA | NA | -1.44326 | 0.29475 | 3 | -65.8061 | 137.7953 | 8.684374 | 0.004588 |
| 5.097032 | NA | NA | NA | -1.25188 | NA | 2 | -66.9346 | 137.96 | 8.849081 | 0.004226 |
| 3.747027 | NA | -0.687 | 0.920087 | -1.45883 | 0.352803 | 5 | -63.8854 | 138.236 | 9.125002 | 0.003681 |
| 6.314582 | NA | -0.41397 | NA | -1.22079 | NA | 3 | -66.0644 | 138.312 | 9.201055 | 0.003544 |
| 3.437144 | NA | -0.5839 | 0.826176 | -1.2186 | NA | 4 | -65.4475 | 139.2027 | 10.09176 | 0.00227 |
| 4.977191 | -0.03831 | NA | NA | -1.18113 | NA | 3 | -66.733 | 139.6493 | 10.5383 | 0.001816 |
| 4.020907 | NA | NA | 0.321921 | -1.43858 | 0.29426 | 4 | -65.7015 | 139.7107 | 10.5997 | 0.001761 |
| 3.759879 | NA | NA | 0.320169 | -1.24835 | NA | 3 | -66.8261 | 139.8354 | 10.72444 | 0.001654 |
| 6.171053 | -0.03013 | -0.39688 | NA | -1.16659 | NA | 4 | -65.9433 | 140.1943 | 11.08338 | 0.001383 |
| 3.383315 | -0.02605 | -0.56512 | 0.806484 | -1.17223 | NA | 5 | -65.3582 | 141.1814 | 12.07047 | 0.000844 |
| 3.66955 | -0.03793 | NA | 0.313808 | -1.17888 | NA | 4 | -66.6287 | 141.5651 | 12.45418 | 0.000697 |
| 2.541188 | -0.43705 | -0.50449 | NA | NA | 1.14575 | 4 | -68.991 | 146.2896 | 17.17863 | 6.56E-05 |
| 1.0435 | -0.39804 | NA | NA | NA | 0.971962 | 3 | -70.2704 | 146.7239 | 17.61298 | 5.28E-05 |
| 1.503151 | -0.4283 | -0.56017 | 0.297562 | NA | 1.116371 | 5 | -68.8672 | 148.1996 | 19.08865 | 2.53E-05 |
| 1.083593 | -0.39835 | NA | -0.00985 | NA | 0.973112 | 4 | -70.2702 | 148.8481 | 19.73717 | 1.83E-05 |
| 1.435184 | -0.12578 | NA | NA | NA | NA | 2 | -73.6713 | 151.4334 | 22.32247 | 5.01E-06 |
| 2.751415 | -0.11598 | -0.42137 | NA | NA | NA | 3 | -72.6387 | 151.4606 | 22.34962 | 4.95E-06 |
| 1.219816 | -0.11517 | -0.50814 | 0.437893 | NA | NA | 4 | -72.368 | 153.0437 | 23.9327 | 2.24E-06 |
| 0.974163 | -0.1259 | NA | 0.111699 | NA | NA | 3 | -73.652 | 153.4872 | 24.37622 | 1.80E-06 |
| 2.654593 | NA | -0.49223 | NA | NA | NA | 2 | -74.7043 | 153.4995 | 24.38854 | 1.78E-06 |
| 1.08876 | NA | NA | NA | NA | NA | 1 | -76.1887 | 154.4074 | 25.29644 | 1.13E-06 |
| 1.098082 | NA | -0.5785 | 0.443927 | NA | NA | 3 | -74.4052 | 154.9937 | 25.8827 | 8.45E-07 |
| 2.668344 | NA | -0.48545 | NA | NA | -0.03167 | 3 | -74.6885 | 155.5602 | 26.44925 | 6.37E-07 |
| 1.169051 | NA | NA | NA | NA | -0.07177 | 2 | -76.1062 | 156.3033 | 27.19238 | 4.39E-07 |
| 0.710857 | NA | NA | 0.091521 | NA | NA | 2 | -76.1748 | 156.4404 | 27.32948 | 4.10E-07 |
| 1.098709 | NA | -0.57183 | 0.448324 | NA | -0.03576 | 4 | -74.385 | 157.0777 | 27.96671 | 2.98E-07 |
| 0.740365 | NA | NA | 0.104306 | NA | -0.07356 | 3 | -76.0883 | 158.3599 | 29.2489 | 1.57E-07 |

#### **Table S7 – Model selection for legumes under irrigation, expansion, mean**

| (Intercept) | I((log(size))^2) | log(Nf) | log(Ng) | log(Nl + exp(-1)) | log(size) | | df | logLik | AICc | delta | weight |
| --- | --- | --- | --- | --- | --- | --- | --- | --- | --- | --- | --- |
| **1.044617** | **NA** | **NA** | **NA** | **-0.19373** | | **0.597585** | **4** | **-136.395** | **281.2059** | **0** | **0.15725** |
| 0.57963 | NA | NA | NA | NA | 0.521598 | | 3 | -137.716 | 281.6797 | 0.473764 | 0.124084 |
| -0.1343 | NA | NA | 0.312217 | -0.23326 | 0.597022 | | 5 | -135.647 | 281.9261 | 0.720184 | 0.1097 |
| 0.953468 | -0.10646 | NA | NA | -0.19676 | 0.895506 | | 5 | -135.733 | 282.0981 | 0.892178 | 0.10066 |
| 0.48439 | -0.10302 | NA | NA | NA | 0.80872 | | 4 | -137.113 | 282.6426 | 1.436633 | 0.076671 |
| 0.790964 | NA | 0.096759 | NA | -0.2041 | 0.584579 | | 5 | -136.14 | 282.9122 | 1.706264 | 0.067001 |
| -0.10566 | -0.09507 | NA | 0.283077 | -0.23227 | 0.863121 | | 6 | -135.119 | 283.1322 | 1.926259 | 0.060022 |
| -0.16888 | NA | NA | 0.183468 | NA | 0.512157 | | 4 | -137.447 | 283.3109 | 2.104971 | 0.054891 |
| 0.38274 | NA | 0.068397 | NA | NA | 0.50953 | | 4 | -137.591 | 283.5979 | 2.39199 | 0.047553 |
| -0.21403 | NA | 0.066404 | 0.287231 | -0.23721 | 0.588142 | | 6 | -135.531 | 283.955 | 2.749068 | 0.039778 |
| 0.744189 | -0.10075 | 0.081697 | NA | -0.20535 | 0.86855 | | 6 | -135.552 | 283.9968 | 2.79092 | 0.038954 |
| -0.13968 | -0.09646 | NA | 0.154452 | NA | 0.782505 | | 5 | -136.923 | 284.4768 | 3.270879 | 0.030643 |
| 0.334218 | -0.09919 | 0.053397 | NA | NA | 0.788625 | | 5 | -137.036 | 284.7044 | 3.498467 | 0.027347 |
| -0.17285 | -0.09201 | 0.055183 | 0.26325 | -0.23559 | 0.847181 | | 7 | -135.039 | 285.2813 | 4.075394 | 0.020494 |
| -0.22753 | NA | 0.048483 | 0.163633 | NA | 0.504624 | | 5 | -137.387 | 285.4054 | 4.199473 | 0.019261 |
| -0.18517 | -0.09442 | 0.037094 | 0.13989 | NA | 0.771017 | | 6 | -136.887 | 286.6681 | 5.462187 | 0.010245 |
| 0.826739 | 0.15132 | NA | NA | NA | NA | | 3 | -141.166 | 288.5787 | 7.372752 | 0.003941 |
| 1.152627 | 0.166683 | NA | NA | -0.12677 | NA | | 4 | -140.623 | 289.6621 | 8.456178 | 0.002293 |
| -0.24086 | 0.148479 | NA | 0.259602 | NA | NA | | 4 | -140.656 | 289.7291 | 8.523133 | 0.002217 |
| 0.462914 | 0.145611 | 0.122432 | NA | NA | NA | | 4 | -140.78 | 289.9776 | 8.771652 | 0.001958 |
| -0.22299 | 0.168635 | NA | 0.365181 | -0.17585 | NA | | 5 | -139.682 | 289.9955 | 8.78962 | 0.001941 |
| 0.763848 | 0.162144 | 0.147541 | NA | -0.14609 | NA | | 5 | -140.069 | 290.7698 | 9.563881 | 0.001318 |
| -0.35301 | 0.144519 | 0.094553 | 0.218551 | NA | NA | | 5 | -140.437 | 291.5063 | 10.30036 | 0.000912 |
| -0.35635 | 0.164918 | 0.113193 | 0.321403 | -0.18479 | NA | | 6 | -139.364 | 291.6226 | 10.41664 | 0.00086 |
| 0.428393 | NA | 0.231847 | NA | NA | NA | | 3 | -148.903 | 304.0536 | 22.84766 | 1.72E-06 |
| 1.139726 | NA | NA | NA | NA | NA | | 2 | -150.112 | 304.3462 | 23.14029 | 1.49E-06 |
| -0.31302 | NA | NA | 0.351324 | NA | NA | | 3 | -149.323 | 304.8931 | 23.68722 | 1.13E-06 |
| -0.54345 | NA | 0.197653 | 0.260396 | NA | NA | | 4 | -148.487 | 305.3916 | 24.18571 | 8.81E-07 |
| 0.886515 | NA | NA | NA | 0.089748 | NA | | 3 | -149.842 | 305.9316 | 24.7257 | 6.72E-07 |
| 0.321805 | NA | 0.218377 | NA | 0.052427 | NA | | 4 | -148.813 | 306.0428 | 24.83693 | 6.36E-07 |
| -0.31527 | NA | NA | 0.318314 | 0.04918 | NA | | 4 | -149.248 | 306.9122 | 25.70626 | 4.12E-07 |
| -0.53928 | NA | 0.193077 | 0.245402 | 0.025474 | NA | | 5 | -148.467 | 307.5665 | 26.36057 | 2.97E-07 |

#### **Table S8 – Model selection for legumes under irrigation, expansion, s.d.**

| (Intercept) | I((log(size))^2) | log(Nf) | log(Ng) | log(Nl + exp(-1)) | log(size) | df | logLik | AICc | delta | weight |
| --- | --- | --- | --- | --- | --- | --- | --- | --- | --- | --- |
| **0.574748** | **-0.08649** | **NA** | **NA** | **NA** | **0.328548** | **4** | **-82.0032** | **172.423** | **0** | **0.103674** |
| 0.654708 | NA | NA | NA | NA | 0.087492 | 3 | -83.2611 | 172.7697 | 0.346688 | 0.087174 |
| 0.748657 | NA | NA | NA | NA | NA | 2 | -84.41 | 172.9425 | 0.519519 | 0.079957 |
| 0.782958 | -0.08802 | NA | NA | -0.08734 | 0.367069 | 5 | -81.1982 | 173.028 | 0.605023 | 0.076611 |
| 1.242255 | -0.0935 | NA | -0.1652 | NA | 0.356588 | 5 | -81.3517 | 173.335 | 0.912035 | 0.065709 |
| 0.858314 | NA | NA | NA | -0.08483 | 0.120765 | 4 | -82.5205 | 173.4577 | 1.034714 | 0.061799 |
| 1.21395 | NA | NA | -0.13708 | NA | 0.094545 | 4 | -82.8189 | 174.0545 | 1.631504 | 0.045856 |
| 0.713829 | 0.016838 | NA | NA | NA | NA | 3 | -84.0358 | 174.319 | 1.895982 | 0.040176 |
| 0.666317 | -0.08882 | -0.03256 | NA | NA | 0.340801 | 5 | -81.9184 | 174.4684 | 2.045399 | 0.037284 |
| 1.187343 | NA | NA | -0.10609 | NA | NA | 3 | -84.1471 | 174.5417 | 2.118692 | 0.035942 |
| 1.252739 | -0.09307 | NA | -0.12556 | -0.07159 | 0.381434 | 6 | -80.8435 | 174.5806 | 2.157555 | 0.03525 |
| 0.826364 | NA | NA | NA | -0.02754 | NA | 3 | -84.3169 | 174.8812 | 2.458166 | 0.030331 |
| 0.709768 | NA | -0.01913 | NA | NA | 0.090866 | 4 | -83.2323 | 174.8813 | 2.458278 | 0.030329 |
| 0.71791 | NA | 0.010021 | NA | NA | NA | 3 | -84.4018 | 175.0511 | 2.628097 | 0.02786 |
| 0.836306 | -0.08947 | -0.02083 | NA | -0.08515 | 0.373941 | 6 | -81.1635 | 175.2207 | 2.797653 | 0.025596 |
| 1.224706 | NA | NA | -0.09703 | -0.07255 | 0.120939 | 5 | -82.3119 | 175.2553 | 2.832291 | 0.025156 |
| 1.259354 | -0.09427 | -0.01394 | -0.15973 | NA | 0.360907 | 6 | -81.3367 | 175.567 | 3.144039 | 0.021525 |
| 0.877844 | NA | -0.00745 | NA | -0.08403 | 0.121766 | 5 | -82.5162 | 175.6639 | 3.240885 | 0.020508 |
| 0.864593 | 0.023945 | NA | NA | -0.05865 | NA | 4 | -83.6763 | 175.7692 | 3.346214 | 0.019456 |
| 1.19615 | 0.018121 | NA | -0.11728 | NA | NA | 4 | -83.7141 | 175.8448 | 3.421831 | 0.018734 |
| 1.217063 | NA | -0.00257 | -0.13602 | NA | 0.094945 | 5 | -82.8184 | 176.2684 | 3.845416 | 0.015158 |
| 0.721932 | 0.016965 | -0.00273 | NA | NA | NA | 4 | -84.0352 | 176.487 | 4.064046 | 0.013589 |
| 1.157622 | NA | 0.025493 | -0.11782 | NA | NA | 4 | -84.097 | 176.6107 | 4.187689 | 0.012774 |
| 1.188046 | NA | NA | -0.0958 | -0.01533 | NA | 4 | -84.1206 | 176.6579 | 4.234888 | 0.012476 |
| 1.263071 | -0.09354 | -0.00849 | -0.12251 | -0.07108 | 0.383885 | 7 | -80.8379 | 176.8801 | 4.457054 | 0.011164 |
| 0.78012 | NA | 0.017883 | NA | -0.0306 | NA | 4 | -84.2919 | 177.0004 | 4.577394 | 0.010512 |
| 1.221199 | NA | 0.002921 | -0.09813 | -0.07272 | 0.120549 | 6 | -82.3112 | 177.516 | 5.093034 | 0.008123 |
| 1.200889 | 0.023468 | NA | -0.08928 | -0.04665 | NA | 5 | -83.5039 | 177.6395 | 5.216477 | 0.007637 |
| 0.84477 | 0.023714 | 0.007523 | NA | -0.05963 | NA | 5 | -83.6719 | 177.9753 | 5.552285 | 0.006457 |
| 1.180787 | 0.017579 | 0.012952 | -0.12291 | NA | NA | 5 | -83.7014 | 178.0344 | 5.611433 | 0.006268 |
| 1.154535 | NA | 0.028884 | -0.10671 | -0.01888 | NA | 5 | -84.0576 | 178.7469 | 6.323863 | 0.00439 |
| 1.179918 | 0.022884 | 0.017799 | -0.09616 | -0.04806 | NA | 6 | -83.4801 | 179.8539 | 7.430892 | 0.002524 |

#### **Table S9 – Model selection for legumes under irrigation, colonisation, mean**

| (Intercept) | log(Nf) | log(Ng) | log(Nl + exp(-1)) | df | logLik | AICc | delta | weight |
| --- | --- | --- | --- | --- | --- | --- | --- | --- |
| **0.427788** | **NA** | **NA** | **NA** | **2** | **-12.4288** | **29.71472** | **0** | **0.464619** |
| -0.21364 | 0.211161 | NA | NA | 3 | -11.7508 | 31.34779 | 1.633076 | 0.205342 |
| -0.3846 | NA | 0.197864 | NA | 3 | -12.2664 | 32.37892 | 2.664207 | 0.122623 |
| 0.458758 | NA | NA | -0.01181 | 3 | -12.4274 | 32.70095 | 2.986235 | 0.104386 |
| -0.71456 | 0.198787 | 0.131159 | NA | 4 | -11.6764 | 34.68607 | 4.971351 | 0.038688 |
| -0.30565 | 0.215217 | NA | 0.030394 | 4 | -11.7411 | 34.81558 | 5.100866 | 0.036263 |
| -0.94927 | NA | 0.277857 | 0.090103 | 4 | -12.2107 | 35.75474 | 6.040025 | 0.022674 |
| -1.42761 | 0.204548 | 0.228884 | 0.112254 | 5 | -11.584 | 38.62264 | 8.907925 | 0.005405 |

#### **Table S10 – Model selection for legumes under irrigation, colonisation, s.d.**

| (Intercept) | log(Nf) | log(Ng) | log(Nl + exp(-1)) | df | logLik | AICc | delta | weight |
| --- | --- | --- | --- | --- | --- | --- | --- | --- |
| **0.452952** | **NA** | **NA** | **NA** | **2** | **1.776799** | **1.303546** | **0** | **0.384106** |
| 0.083261 | NA | NA | 0.141008 | 3 | 2.899545 | 2.047065 | 0.743519 | 0.264848 |
| 0.311555 | 0.046549 | NA | NA | 3 | 1.946913 | 3.952328 | 2.648783 | 0.102159 |
| 0.770955 | NA | -0.07745 | NA | 3 | 1.908917 | 4.028321 | 2.724775 | 0.09835 |
| -0.15515 | 0.067124 | NA | 0.154172 | 4 | 3.29795 | 4.737433 | 3.433887 | 0.068991 |
| -0.27544 | NA | 0.070785 | 0.166972 | 4 | 2.984236 | 5.364862 | 4.061316 | 0.050414 |
| 0.678639 | 0.055616 | -0.09611 | NA | 4 | 2.148737 | 7.03586 | 5.732314 | 0.021862 |
| -0.42638 | 0.064544 | 0.055332 | 0.173962 | 5 | 3.35148 | 8.751586 | 7.44804 | 0.009271 |

#### **Table S11 – Model selection for forbs under irrigation, persistence**

| (Intercept) | I((log(size))^2) | log(Nf) | log(Ng) | log(Nl + exp(-1)) | log(size) | df | logLik | AICc | delta | weight |
| --- | --- | --- | --- | --- | --- | --- | --- | --- | --- | --- |
| 3.821004 | 0.108906 | -0.82922 | NA | -0.26985 | 0.191941 | 5 | -251.085 | 512.3181 | 0 | 0.168396 |
| 2.292898 | NA | -0.79895 | 0.406474 | -0.32807 | 0.276588 | 5 | -251.086 | 512.3206 | 0.002515 | 0.168184 |
| 2.778272 | 0.096427 | -0.92029 | 0.360219 | -0.31848 | 0.222769 | 6 | -250.17 | 512.548 | 0.22989 | 0.15011 |
| **3.42127** | **NA** | **-0.67772** | **NA** | **-0.27368** | **0.250657** | **4** | **-252.283** | **512.6652** | **0.347152** | **0.141562** |
| 3.800954 | 0.138702 | -0.81224 | NA | -0.26417 | NA | 4 | -252.284 | 512.6677 | 0.349632 | 0.141387 |
| 2.997597 | 0.132809 | -0.87987 | 0.275839 | -0.29998 | NA | 5 | -251.731 | 513.6102 | 1.292103 | 0.088258 |
| 3.223914 | NA | -0.58792 | NA | -0.26816 | NA | 3 | -254.503 | 515.0648 | 2.74671 | 0.042647 |
| 2.316704 | NA | -0.67722 | 0.320458 | -0.30929 | NA | 4 | -253.74 | 515.5794 | 3.261351 | 0.032971 |
| 3.363025 | 0.113288 | -0.91998 | NA | NA | 0.18176 | 4 | -254.451 | 517.0006 | 4.682535 | 0.016201 |
| 3.354631 | 0.141712 | -0.90245 | NA | NA | NA | 3 | -255.532 | 517.1225 | 4.804403 | 0.015243 |
| 2.94552 | NA | -0.76542 | NA | NA | 0.244221 | 3 | -255.758 | 517.5754 | 5.257292 | 0.012154 |
| 3.072429 | 0.110037 | -0.94775 | 0.093273 | NA | 0.189141 | 5 | -254.374 | 518.8974 | 6.579354 | 0.006275 |
| 3.235141 | 0.140854 | -0.91354 | 0.038284 | NA | NA | 4 | -255.518 | 519.136 | 6.817882 | 0.00557 |
| 2.537237 | NA | -0.81223 | 0.136391 | NA | 0.251987 | 4 | -255.59 | 519.2792 | 6.961131 | 0.005185 |
| 2.760953 | NA | -0.67533 | NA | NA | NA | 2 | -257.872 | 519.7742 | 7.456144 | 0.004048 |
| 2.524513 | NA | -0.70064 | 0.077973 | NA | NA | 3 | -257.817 | 521.6927 | 9.374657 | 0.001551 |
| 1.524412 | NA | NA | NA | -0.34779 | NA | 2 | -261.821 | 527.6724 | 15.35428 | 7.80E-05 |
| 1.494988 | NA | NA | NA | -0.35593 | 0.120855 | 3 | -261.259 | 528.5776 | 16.25951 | 4.96E-05 |
| 1.900435 | NA | NA | -0.10139 | -0.33308 | NA | 3 | -261.733 | 529.5262 | 17.20809 | 3.09E-05 |
| 1.536244 | -0.02156 | NA | NA | -0.34397 | NA | 3 | -261.741 | 529.5418 | 17.22374 | 3.06E-05 |
| 1.513285 | -0.05106 | NA | NA | -0.34971 | 0.160187 | 4 | -260.879 | 529.8566 | 17.53856 | 2.62E-05 |
| 1.843533 | NA | NA | -0.09385 | -0.34223 | 0.119169 | 4 | -261.184 | 530.4671 | 18.14902 | 1.93E-05 |
| 1.828846 | -0.01632 | NA | -0.07972 | -0.33327 | NA | 4 | -261.691 | 531.482 | 19.16386 | 1.16E-05 |
| 1.609941 | -0.04892 | NA | -0.02629 | -0.34608 | 0.158082 | 5 | -260.874 | 531.896 | 19.57789 | 9.44E-06 |
| 2.039769 | NA | NA | -0.36147 | NA | NA | 2 | -266.794 | 537.618 | 25.29988 | 5.40E-07 |
| 0.556735 | NA | NA | NA | NA | NA | 1 | -268.244 | 538.4984 | 26.18029 | 3.48E-07 |
| 2.003249 | NA | NA | -0.36176 | NA | 0.08745 | 3 | -266.491 | 539.0416 | 26.72354 | 2.65E-07 |
| 1.973557 | -0.01626 | NA | -0.34126 | NA | NA | 3 | -266.752 | 539.5628 | 27.24473 | 2.04E-07 |
| 0.60053 | -0.04242 | NA | NA | NA | NA | 2 | -267.927 | 539.8833 | 27.5652 | 1.74E-07 |
| 0.519089 | NA | NA | NA | NA | 0.087901 | 2 | -267.944 | 539.9184 | 27.60035 | 1.71E-07 |
| 0.566224 | -0.06887 | NA | NA | NA | 0.141953 | 3 | -267.237 | 540.5339 | 28.21584 | 1.26E-07 |
| 1.825052 | -0.04067 | NA | -0.31162 | NA | 0.119607 | 4 | -266.272 | 540.6421 | 28.32396 | 1.19E-07 |

#### **Table S12 – Model selection for forbs under irrigation, expansion, mean**

| (Intercept) | I((log(size))^2) | log(Nf) | log(Ng) | log(Nl + exp(-1)) | log(size) | df | logLik | AICc | delta | weight |
| --- | --- | --- | --- | --- | --- | --- | --- | --- | --- | --- |
| 0.07814 | 0.126839685 | 0.12239 | NA | -0.05157 | NA | 5 | -195.651 | 401.5383 | 0 | 0.143891 |
| 0.06455 | 0.112379681 | 0.122216 | NA | -0.05177 | 0.061359 | 6 | -194.621 | 401.5736 | 0.035394 | 0.141367 |
| 0.03544 | 0.128952984 | 0.092852 | NA | NA | NA | 4 | -197.358 | 402.8723 | 1.334084 | 0.073848 |
| 0.021772 | 0.11459699 | 0.092563 | NA | NA | 0.060953 | 5 | -196.354 | 402.9452 | 1.406908 | 0.071208 |
| 0.306626 | 0.147126336 | NA | NA | NA | NA | 3 | -198.486 | 403.0664 | 1.528153 | 0.067019 |
| 0.292064 | 0.132663209 | NA | NA | NA | 0.061167 | 4 | -197.485 | 403.1263 | 1.588062 | 0.065041 |
| **0.401734** | **0.149831244** | **NA** | **NA** | **-0.03803** | **NA** | **4** | **-197.507** | **403.1702** | **1.631969** | **0.063629** |
| 0.387646 | 0.135301013 | NA | NA | -0.03825 | 0.061518 | 5 | -196.486 | 403.2079 | 1.669665 | 0.062441 |
| 0.189016 | 0.128092033 | 0.126308 | -0.03454 | -0.04538 | NA | 6 | -195.539 | 403.4092 | 1.870968 | 0.056462 |
| 0.304998 | 0.131273356 | 0.110199 | -0.08026 | NA | NA | 5 | -196.625 | 403.4859 | 1.947626 | 0.054339 |
| 0.126629 | 0.113478577 | 0.124401 | -0.01922 | -0.04832 | 0.059653 | 7 | -194.587 | 403.6175 | 2.079264 | 0.050878 |
| 0.254474 | 0.118005405 | 0.10748 | -0.06888 | NA | 0.054937 | 6 | -195.821 | 403.9733 | 2.435085 | 0.042585 |
| 0.522385 | 0.150977335 | NA | -0.05408 | NA | NA | 4 | -198.139 | 404.4351 | 2.896888 | 0.033805 |
| 0.463945 | 0.136595186 | NA | -0.04286 | NA | 0.057445 | 5 | -197.269 | 404.7738 | 3.235581 | 0.028539 |
| 0.457466 | 0.15074531 | NA | -0.01588 | -0.03498 | NA | 5 | -197.483 | 405.202 | 3.663704 | 0.023039 |
| 0.388987 | 0.135330823 | NA | -0.00038 | -0.03818 | 0.061484 | 6 | -196.486 | 405.3037 | 3.765434 | 0.021897 |
| -0.33056 | NA | 0.27632 | NA | -0.05857 | 0.150005 | 5 | -205.744 | 421.7243 | 20.18603 | 5.95E-06 |
| -0.46099 | NA | 0.268048 | 0.043064 | -0.06615 | 0.151884 | 6 | -205.581 | 423.4937 | 21.95548 | 2.46E-06 |
| -0.38792 | NA | 0.246137 | NA | NA | 0.151528 | 4 | -207.784 | 423.7243 | 22.18603 | 2.19E-06 |
| -0.31592 | NA | 0.252498 | -0.02249 | NA | 0.150443 | 5 | -207.731 | 425.6975 | 24.15928 | 8.16E-07 |
| -0.44242 | NA | 0.336387 | NA | -0.06059 | NA | 4 | -212.534 | 433.2247 | 31.68649 | 1.89E-08 |
| -0.50323 | NA | 0.332919 | 0.019864 | -0.0641 | NA | 5 | -212.501 | 435.2378 | 33.69954 | 6.92E-09 |
| -0.50295 | NA | 0.305785 | NA | NA | NA | 3 | -214.606 | 435.3055 | 33.76724 | 6.69E-09 |
| -0.36224 | NA | 0.317252 | -0.04346 | NA | NA | 4 | -214.417 | 436.99 | 35.45173 | 2.88E-09 |
| 0.352814 | NA | NA | NA | NA | 0.203718 | 3 | -217.376 | 440.8457 | 39.30745 | 4.19E-10 |
| -0.04758 | NA | NA | 0.1285 | -0.04757 | 0.206169 | 5 | -215.675 | 441.5863 | 40.04809 | 2.90E-10 |
| 0.040937 | NA | NA | 0.076957 | NA | 0.202814 | 4 | -216.725 | 441.6072 | 40.06891 | 2.87E-10 |
| 0.407565 | NA | NA | NA | -0.02164 | 0.20552 | 4 | -217.1 | 442.3571 | 40.81885 | 1.97E-10 |
| 0.447402 | NA | NA | NA | NA | NA | 2 | -229.612 | 463.2712 | 61.73298 | 5.66E-15 |
| 0.107841 | NA | NA | 0.083675 | NA | NA | 3 | -228.912 | 463.9169 | 62.37862 | 4.10E-15 |
| 0.038329 | NA | NA | 0.124752 | -0.03783 | NA | 4 | -228.306 | 464.7687 | 63.23048 | 2.68E-15 |
| 0.479978 | NA | NA | NA | -0.01268 | NA | 3 | -229.526 | 465.1453 | 63.60708 | 2.22E-15 |

#### **Table S13 – Model selection for forbs under irrigation, expansion, s.d.**

| (Intercept) | I((log(size))^2) | log(Nf) | log(Ng) | log(Nl + exp(-1)) | log(size) | df | logLik | AICc | delta | weight |
| --- | --- | --- | --- | --- | --- | --- | --- | --- | --- | --- |
| 0.492926 | 0.036814 | NA | NA | -0.03007 | -0.03294 | 5 | -29.479 | 69.19416 | 0 | 0.119297 |
| 0.485382 | 0.029033 | NA | NA | -0.03019 | NA | 4 | -30.5366 | 69.23013 | 0.035972 | 0.117171 |
| **0.366867** | **0.027871** | **0.047684** | **NA** | **-0.03535** | **-0.03301** | **6** | **-28.4564** | **69.24483** | **0.05067** | **0.116313** |
| 0.359557 | 0.020093 | 0.04759 | NA | -0.03546 | NA | 5 | -29.5264 | 69.28898 | 0.094826 | 0.113773 |
| 0.277094 | NA | 0.08149 | NA | -0.03688 | NA | 4 | -31.14 | 70.43693 | 1.242772 | 0.064086 |
| 0.443629 | 0.02923 | 0.050385 | -0.02377 | -0.03108 | -0.03511 | 7 | -28.2681 | 70.9807 | 1.786544 | 0.04883 |
| 0.549891 | 0.038081 | NA | -0.01614 | -0.02697 | -0.03437 | 6 | -29.3914 | 71.1148 | 1.920641 | 0.045663 |
| 0.406905 | 0.020628 | 0.049263 | -0.01475 | -0.03281 | NA | 6 | -29.4528 | 71.23765 | 2.043494 | 0.042943 |
| 0.511607 | 0.029463 | NA | -0.00747 | -0.02876 | NA | 5 | -30.5176 | 71.2714 | 2.07724 | 0.042224 |
| 0.417784 | 0.03474 | NA | NA | NA | -0.03322 | 4 | -31.6993 | 71.55538 | 2.361225 | 0.036635 |
| 0.409876 | 0.026886 | NA | NA | NA | NA | 3 | -32.7565 | 71.60682 | 2.412664 | 0.035705 |
| 0.602844 | 0.038974 | NA | -0.04614 | NA | -0.03723 | 5 | -30.8009 | 71.83803 | 2.643874 | 0.031807 |
| 0.268877 | NA | 0.085903 | NA | -0.03703 | -0.01102 | 5 | -30.9958 | 72.22787 | 3.033717 | 0.026174 |
| 0.564974 | 0.029654 | NA | -0.03887 | NA | NA | 4 | -32.1139 | 72.38458 | 3.190422 | 0.024201 |
| 0.295427 | NA | 0.082535 | -0.00599 | -0.03583 | NA | 5 | -31.1279 | 72.49196 | 3.297807 | 0.022936 |
| 0.525857 | 0.032141 | 0.039502 | -0.05571 | NA | -0.03815 | 6 | -30.099 | 72.53001 | 3.335854 | 0.022504 |
| 0.337661 | 0.029385 | 0.027439 | NA | NA | -0.03328 | 5 | -31.3448 | 72.92581 | 3.731654 | 0.018463 |
| 0.330198 | 0.021546 | 0.027281 | NA | NA | NA | 4 | -32.409 | 72.97481 | 3.780655 | 0.018017 |
| 0.490773 | 0.022928 | 0.037614 | -0.04781 | NA | NA | 5 | -31.4833 | 73.20278 | 4.008618 | 0.016076 |
| 0.292269 | NA | 0.087386 | -0.00772 | -0.03567 | -0.01136 | 6 | -30.9757 | 74.28344 | 5.08928 | 0.009365 |
| 0.240242 | NA | 0.062859 | NA | NA | NA | 3 | -34.2276 | 74.54899 | 5.354831 | 0.008201 |
| 0.374233 | NA | 0.073778 | -0.04138 | NA | NA | 4 | -33.5388 | 75.23442 | 6.040259 | 0.005821 |
| 0.232607 | NA | 0.066818 | NA | NA | -0.01006 | 4 | -34.1103 | 76.37747 | 7.183313 | 0.003287 |
| 0.500544 | NA | NA | NA | -0.02528 | NA | 3 | -35.3702 | 76.83421 | 7.640052 | 0.002616 |
| 0.370497 | NA | 0.079001 | -0.04307 | NA | -0.01213 | 5 | -33.3682 | 76.97263 | 7.778478 | 0.002441 |
| 0.435601 | NA | NA | NA | NA | NA | 2 | -36.8921 | 77.83097 | 8.636811 | 0.001589 |
| 0.429687 | NA | NA | 0.020015 | -0.02931 | NA | 4 | -35.231 | 78.61886 | 9.424703 | 0.001072 |
| 0.498346 | NA | NA | NA | -0.02555 | 0.006238 | 4 | -35.3216 | 78.80008 | 9.605919 | 0.000979 |
| 0.483552 | NA | NA | -0.01182 | NA | NA | 3 | -36.8308 | 79.75525 | 10.5611 | 0.000607 |
| 0.433692 | NA | NA | NA | NA | 0.00411 | 3 | -36.8712 | 79.83615 | 10.64199 | 0.000583 |
| 0.427045 | NA | NA | 0.02013 | -0.02961 | 0.00634 | 5 | -35.1807 | 80.59767 | 11.40352 | 0.000398 |
| 0.48215 | NA | NA | -0.01196 | NA | 0.004251 | 4 | -36.8084 | 81.77357 | 12.57942 | 0.000221 |

#### **Table S14 – Model selection for forbs under irrigation, colonisation, mean**

| (Intercept) | log(Nf) | log(Ng) | log(Nl + exp(-1)) | df | logLik | AICc | delta | weight |
| --- | --- | --- | --- | --- | --- | --- | --- | --- |
| **-0.2914** | **0.178925** | **NA** | **NA** | **3** | **-99.7923** | **205.797** | **0** | **0.297998** |
| -0.47799 | 0.16185 | NA | 0.085702 | 4 | -98.9079 | 206.173 | 0.375983 | 0.246928 |
| -0.66855 | 0.161774 | 0.10457 | NA | 4 | -99.4887 | 207.3346 | 1.537581 | 0.138144 |
| -0.76205 | 0.149468 | 0.081982 | 0.080362 | 5 | -98.7222 | 207.9848 | 2.187815 | 0.099801 |
| -0.03818 | NA | NA | 0.109461 | 3 | -101.204 | 208.6205 | 2.823445 | 0.072629 |
| 0.268377 | NA | NA | NA | 2 | -102.626 | 209.3582 | 3.561157 | 0.050225 |
| -0.61977 | NA | 0.150074 | 0.096359 | 4 | -100.559 | 209.4745 | 3.677483 | 0.047387 |
| -0.49151 | NA | 0.184445 | NA | 3 | -101.642 | 209.4957 | 3.698643 | 0.046888 |

#### **Table S15 – Model selection for forbs under irrigation, colonisation, s.d.**

| (Intercept) | log(Nf) | log(Ng) | log(Nl + exp(-1)) | df | logLik | AICc | delta | weight |
| --- | --- | --- | --- | --- | --- | --- | --- | --- |
| **-0.11691** | **0.162892** | **NA** | **NA** | **3** | **-56.9149** | **120.0422** | **0** | **0.434437** |
| -0.22797 | 0.15273 | NA | 0.051008 | 4 | -56.2642 | 120.8855 | 0.843316 | 0.284972 |
| -0.08889 | 0.164166 | -0.00777 | NA | 4 | -56.9114 | 122.18 | 2.137795 | 0.14918 |
| -0.14994 | 0.156131 | -0.02252 | 0.052475 | 5 | -56.2351 | 123.0108 | 2.968676 | 0.098466 |
| 0.18706 | NA | NA | 0.073429 | 3 | -60.4345 | 127.0814 | 7.039232 | 0.012864 |
| 0.392703 | NA | NA | NA | 2 | -61.721 | 127.5473 | 7.505135 | 0.010191 |
| -0.00131 | NA | 0.048608 | 0.069185 | 4 | -60.2992 | 128.9555 | 8.913332 | 0.00504 |
| 0.090776 | NA | 0.073286 | NA | 3 | -61.41 | 129.0323 | 8.990138 | 0.00485 |

#### **Table S16 – Model selection for grasses under control, persistence**

| (Intercept) | I((log(size))^2) | log(Nf) | log(Ng) | log(Nl + exp(-1)) | log(size) | df | logLik | AICc | delta | weight |
| --- | --- | --- | --- | --- | --- | --- | --- | --- | --- | --- |
| **6.253228** | **NA** | **-0.41792** | **-0.61073** | **-0.677673566** | **0.510313** | **5** | **-310.31** | **630.729** | **0** | **0.340088** |
| 4.046939 | NA | -0.46167 | NA | -0.707951238 | 0.498471 | 4 | -311.509 | 631.0906 | 0.361608 | 0.283837 |
| 6.331535 | 0.013507 | -0.42011 | -0.6272 | -0.678728611 | 0.480332 | 6 | -310.278 | 632.7097 | 1.980722 | 0.126323 |
| 4.046544 | -0.0003 | -0.4616 | NA | -0.70791079 | 0.499143 | 5 | -311.509 | 633.127 | 2.398071 | 0.102532 |
| 5.444339 | NA | NA | -0.7628 | -0.66108201 | 0.495845 | 4 | -312.834 | 633.7402 | 3.011175 | 0.075461 |
| 2.484742 | NA | NA | NA | -0.692600076 | 0.480358 | 3 | -314.72 | 635.4831 | 4.754124 | 0.031568 |
| 5.486988 | 0.007759 | NA | -0.77285 | -0.6616212 | 0.478715 | 5 | -312.823 | 635.7544 | 5.0254 | 0.027564 |
| 2.481078 | -0.00916 | NA | NA | -0.691524481 | 0.500519 | 4 | -314.703 | 637.4794 | 6.750429 | 0.011635 |
| 7.103238 | 0.147864 | -0.42566 | -0.76021 | -0.685308491 | NA | 5 | -316.652 | 643.4122 | 12.68318 | 0.000599 |
| 4.336559 | 0.137243 | -0.47378 | NA | -0.720727734 | NA | 4 | -318.523 | 645.1192 | 14.39025 | 0.000255 |
| 6.227156 | 0.141215 | NA | -0.90407 | -0.666390261 | NA | 4 | -319.324 | 646.7214 | 15.99239 | 0.000115 |
| 2.722921 | 0.128256 | NA | NA | -0.702014514 | NA | 3 | -322.002 | 650.0475 | 19.31852 | 2.17E-05 |
| 5.764781 | NA | -0.40987 | -1.05367 | NA | 0.496959 | 4 | -324.496 | 657.0653 | 26.33633 | 6.50E-07 |
| 5.814225 | 0.010513 | -0.41139 | -1.06463 | NA | 0.473854 | 5 | -324.475 | 659.0599 | 28.33088 | 2.40E-07 |
| 4.898968 | NA | NA | -1.17149 | NA | 0.483874 | 3 | -326.917 | 659.8777 | 29.14875 | 1.59E-07 |
| 4.924392 | 0.005742 | NA | -1.17782 | NA | 0.471314 | 4 | -326.91 | 661.8936 | 31.16459 | 5.81E-08 |
| 4.283448 | NA | -0.39364 | NA | -0.676029778 | NA | 3 | -328.981 | 664.0055 | 33.27651 | 2.02E-08 |
| 1.811524 | NA | -0.49143 | NA | NA | 0.480712 | 3 | -329.356 | 664.7549 | 34.02593 | 1.39E-08 |
| 5.757138 | NA | -0.36184 | -0.41222 | -0.65047091 | NA | 4 | -328.381 | 664.8349 | 34.10596 | 1.34E-08 |
| 1.80273 | -0.01337 | -0.48868 | NA | NA | 0.509759 | 4 | -329.32 | 666.7118 | 35.98286 | 5.22E-09 |
| 5.071255 | NA | NA | -0.55022 | -0.636688593 | NA | 3 | -330.447 | 666.9382 | 36.20917 | 4.67E-09 |
| 2.94959 | NA | NA | NA | -0.667340639 | NA | 2 | -331.537 | 667.0961 | 36.36712 | 4.31E-09 |
| 0.203131 | NA | NA | NA | NA | 0.466367 | 2 | -332.961 | 669.944 | 39.21499 | 1.04E-09 |
| 6.444964 | 0.142633 | -0.41343 | -1.17249 | NA | NA | 4 | -331.162 | 670.3974 | 39.66838 | 8.27E-10 |
| 0.204737 | -0.02041 | NA | NA | NA | 0.510324 | 3 | -332.871 | 671.7848 | 41.05579 | 4.13E-10 |
| 5.549179 | 0.136792 | NA | -1.28601 | NA | NA | 3 | -333.693 | 673.43 | 42.70107 | 1.82E-10 |
| 2.039414 | 0.127511 | -0.49755 | NA | NA | NA | 3 | -337.232 | 680.5074 | 49.77841 | 5.28E-12 |
| 0.411347 | 0.120239 | NA | NA | NA | NA | 2 | -341.07 | 686.1612 | 55.43217 | 3.12E-13 |
| 5.582846 | NA | -0.35498 | -0.91201 | NA | NA | 3 | -342.932 | 691.908 | 61.17899 | 1.76E-14 |
| 4.86341 | NA | NA | -1.02441 | NA | NA | 2 | -344.917 | 693.8548 | 63.12586 | 6.67E-15 |
| 2.174474 | NA | -0.43488 | NA | NA | NA | 2 | -346.97 | 697.9621 | 67.23309 | 8.55E-16 |
| 0.736632 | NA | NA | NA | NA | NA | 1 | -350.089 | 702.1861 | 71.45708 | 1.03E-16 |

#### **Table S17 – Model selection for grasses under control, expansion, mean**

| (Intercept) | I((log(size))^2) | log(Nf) | log(Ng) | log(Nl + exp(-1)) | log(size) | df | logLik | AICc | delta | weight |
| --- | --- | --- | --- | --- | --- | --- | --- | --- | --- | --- |
| 2.700579908 | 0.070355 | -0.14237 | -0.39858 | NA | 0.399754 | 6 | -468.644 | 949.5156 | 0 | 0.361764 |
| **2.345211743** | **0.073569** | **NA** | **-0.42451** | **NA** | **0.389544** | **5** | **-470.014** | **950.1894** | **0.673807** | **0.258291** |
| 2.748943579 | 0.071338 | -0.14258 | -0.4212 | 0.01424 | 0.397234 | 7 | -468.585 | 951.4743 | 1.958666 | 0.135864 |
| 2.391464249 | 0.074523 | NA | -0.44641 | 0.013764 | 0.387095 | 6 | -469.959 | 952.1455 | 2.629812 | 0.097133 |
| 2.562872441 | NA | -0.15641 | -0.36863 | NA | 0.609053 | 5 | -471.587 | 953.3367 | 3.821074 | 0.053541 |
| 2.163768751 | NA | NA | -0.39574 | NA | 0.608336 | 4 | -473.221 | 954.5494 | 5.033722 | 0.029199 |
| 2.576564808 | NA | -0.15653 | -0.37518 | 0.004199 | 0.609172 | 6 | -471.582 | 955.392 | 5.876355 | 0.01916 |
| 1.217169906 | 0.062342 | -0.17268 | NA | NA | 0.414812 | 5 | -473.088 | 956.339 | 6.823315 | 0.011933 |
| 2.173907319 | NA | NA | -0.40071 | 0.003179 | 0.608426 | 5 | -473.218 | 956.5979 | 7.082271 | 0.010484 |
| 1.312798894 | 0.060932 | -0.16754 | NA | -0.03802 | 0.419256 | 6 | -472.576 | 957.3798 | 7.864198 | 0.007091 |
| 0.661392677 | 0.065653 | NA | NA | NA | 0.40346 | 4 | -475.08 | 958.2688 | 8.753125 | 0.004547 |
| 1.193693337 | NA | -0.18319 | NA | NA | 0.600863 | 4 | -475.369 | 958.8457 | 9.330017 | 0.003408 |
| 0.786184577 | 0.063975 | NA | NA | -0.0423 | 0.40878 | 5 | -474.451 | 959.0634 | 9.547777 | 0.003056 |
| 1.2993644 | NA | -0.17728 | NA | -0.04178 | 0.601122 | 5 | -474.756 | 959.6745 | 10.15884 | 0.002251 |
| 0.600857675 | NA | NA | NA | NA | 0.599298 | 3 | -477.59 | 961.2437 | 11.7281 | 0.001027 |
| 0.739806336 | NA | NA | NA | -0.04652 | 0.599643 | 4 | -476.836 | 961.7791 | 12.26347 | 0.000786 |
| 2.638973689 | 0.182877 | NA | -0.44978 | NA | NA | 4 | -478.381 | 964.8707 | 15.35509 | 0.000168 |
| 2.941072205 | 0.182587 | -0.11847 | -0.42875 | NA | NA | 5 | -477.472 | 965.1055 | 15.58988 | 0.000149 |
| 2.726485904 | 0.183404 | NA | -0.49262 | 0.027125 | NA | 5 | -478.177 | 966.5157 | 17.00009 | 7.36E-05 |
| 3.032580828 | 0.183126 | -0.11916 | -0.47256 | 0.027816 | NA | 6 | -477.255 | 966.7385 | 17.22286 | 6.58E-05 |
| 1.350572047 | 0.178504 | -0.15019 | NA | NA | NA | 4 | -482.386 | 972.8806 | 23.36501 | 3.05E-06 |
| 0.862271471 | 0.178619 | NA | NA | NA | NA | 3 | -483.828 | 973.7202 | 24.2046 | 2.01E-06 |
| 1.42809159 | 0.178372 | -0.14589 | NA | -0.03036 | NA | 5 | -482.075 | 974.3119 | 24.7963 | 1.49E-06 |
| 0.965526721 | 0.178466 | NA | NA | -0.03427 | NA | 4 | -483.432 | 974.9727 | 25.4571 | 1.07E-06 |
| 1.472597804 | NA | NA | NA | NA | NA | 2 | -569.349 | 1142.731 | 193.2152 | 4.00E-43 |
| 1.984258076 | NA | -0.1575 | NA | NA | NA | 3 | -568.345 | 1142.754 | 193.2382 | 3.96E-43 |
| 2.391893792 | NA | NA | -0.23083 | NA | NA | 3 | -568.44 | 1142.944 | 193.4283 | 3.60E-43 |
| 2.75534626 | NA | -0.14234 | -0.20598 | NA | NA | 4 | -567.626 | 1143.36 | 193.8445 | 2.92E-43 |
| 1.599417804 | NA | NA | NA | -0.04231 | NA | 3 | -568.967 | 1143.998 | 194.4828 | 2.12E-43 |
| 2.08126623 | NA | -0.15209 | NA | -0.03823 | NA | 4 | -568.032 | 1144.172 | 194.6562 | 1.95E-43 |
| 2.337781423 | NA | NA | -0.20452 | -0.01691 | NA | 4 | -568.39 | 1144.888 | 195.3726 | 1.36E-43 |
| 2.702998805 | NA | -0.1419 | -0.18115 | -0.01601 | NA | 5 | -567.582 | 1145.325 | 195.8098 | 1.09E-43 |

#### **Table S18 – Model selection for grasses under control, expansion, s.d.**

| (Intercept) | I((log(size))^2) | log(Nf) | log(Ng) | log(Nl + exp(-1)) | log(size) | df | logLik | AICc | delta | weight |
| --- | --- | --- | --- | --- | --- | --- | --- | --- | --- | --- |
| 0.148751 | NA | 0.144297 | NA | NA | 0.040827 | 4 | -270.952 | 550.0114 | 0 | 0.18764 |
| **0.141062** | **-0.02042** | **0.140855** | **NA** | **NA** | **0.101766** | **5** | **-270.229** | **550.6202** | **0.608857** | **0.138393** |
| 0.202468 | NA | 0.146042 | NA | NA | NA | 3 | -272.592 | 551.2481 | 1.236707 | 0.101106 |
| 0.126647 | NA | 0.143061 | NA | 0.008739 | 0.040773 | 5 | -270.872 | 551.9069 | 1.895507 | 0.072731 |
| 0.173789 | 0.008079 | 0.146373 | NA | NA | NA | 4 | -271.909 | 551.9263 | 1.914865 | 0.072031 |
| 0.236321 | NA | 0.146009 | -0.02358 | NA | 0.041351 | 5 | -270.906 | 551.9749 | 1.963478 | 0.070301 |
| 0.122206 | -0.02014 | 0.139842 | NA | 0.007496 | 0.10089 | 6 | -270.171 | 552.5687 | 2.557294 | 0.052242 |
| 0.196944 | -0.02012 | 0.141997 | -0.01502 | NA | 0.101199 | 6 | -270.211 | 552.6491 | 2.63766 | 0.050184 |
| 0.179682 | NA | 0.14477 | NA | 0.00898 | NA | 4 | -272.509 | 553.1251 | 3.113748 | 0.039553 |
| 0.249389 | NA | 0.146965 | -0.01253 | NA | NA | 4 | -272.579 | 553.2658 | 3.254399 | 0.036867 |
| 0.283199 | NA | 0.145605 | -0.04599 | 0.014375 | 0.041759 | 6 | -270.732 | 553.6925 | 3.681138 | 0.029784 |
| 0.14995 | 0.008119 | 0.145052 | NA | 0.009338 | NA | 5 | -271.819 | 553.8002 | 3.788806 | 0.028223 |
| 0.257826 | 0.008294 | 0.148049 | -0.02265 | NA | NA | 5 | -271.868 | 553.8976 | 3.886225 | 0.026881 |
| 0.236532 | -0.01931 | 0.141829 | -0.03353 | 0.011656 | 0.099137 | 7 | -270.097 | 554.4985 | 4.487084 | 0.019905 |
| 0.291867 | NA | 0.146608 | -0.03269 | 0.01299 | NA | 5 | -272.438 | 555.0384 | 5.027002 | 0.015196 |
| 0.307318 | 0.008585 | 0.147674 | -0.04635 | 0.015044 | NA | 6 | -271.679 | 555.5855 | 5.574065 | 0.011559 |
| 0.615722 | NA | NA | NA | NA | 0.04206 | 3 | -275.019 | 556.1022 | 6.090798 | 0.008927 |
| 0.594404 | -0.02312 | NA | NA | NA | 0.111026 | 4 | -274.109 | 556.3252 | 6.313799 | 0.007986 |
| 0.676902 | NA | NA | NA | NA | NA | 2 | -276.722 | 557.4771 | 7.465684 | 0.004489 |
| 0.57819 | NA | NA | NA | 0.012566 | 0.041966 | 4 | -274.858 | 557.8228 | 7.811441 | 0.003777 |
| 0.561747 | -0.02268 | NA | NA | 0.011069 | 0.109634 | 5 | -273.983 | 558.1286 | 8.11724 | 0.003241 |
| 0.60889 | NA | NA | 0.00173 | NA | 0.04202 | 4 | -275.019 | 558.145 | 8.133614 | 0.003215 |
| 0.649683 | 0.007966 | NA | NA | NA | NA | 3 | -276.073 | 558.2109 | 8.199549 | 0.00311 |
| 0.551369 | -0.02332 | NA | 0.01085 | NA | 0.111381 | 5 | -274.099 | 558.3605 | 8.349123 | 0.002886 |
| 0.638351 | NA | NA | NA | 0.012861 | NA | 3 | -276.555 | 559.1745 | 9.163074 | 0.001921 |
| 0.624648 | NA | NA | 0.013121 | NA | NA | 3 | -276.709 | 559.4816 | 9.470219 | 0.001648 |
| 0.657763 | NA | NA | -0.02223 | 0.015323 | 0.042454 | 5 | -274.825 | 559.8126 | 9.801177 | 0.001396 |
| 0.609847 | 0.008025 | NA | NA | 0.013222 | NA | 4 | -275.896 | 559.899 | 9.887646 | 0.001337 |
| 0.592132 | -0.02248 | NA | -0.00845 | 0.01213 | 0.109223 | 6 | -273.979 | 560.1849 | 10.17348 | 0.001159 |
| 0.635364 | 0.007932 | NA | 0.003625 | NA | NA | 4 | -276.072 | 560.2521 | 10.24074 | 0.001121 |
| 0.669198 | NA | NA | -0.00854 | 0.013922 | NA | 4 | -276.55 | 561.2082 | 11.19685 | 0.000695 |
| 0.686662 | 0.00824 | NA | -0.02149 | 0.0159 | NA | 5 | -275.866 | 561.8939 | 11.8825 | 0.000493 |

#### **Table S19 – Model selection for grasses under control, colonisation, mean**

| (Intercept) | log(Nf) | log(Ng) | log(Nl + exp(-1)) | df | logLik | AICc | delta | weight |
| --- | --- | --- | --- | --- | --- | --- | --- | --- |
| **-0.86403** | **NA** | **0.385529** | **NA** | **3** | **-139.431** | **285.0805** | **0** | **0.256266** |
| 0.676877 | NA | NA | NA | 2 | -140.596 | 285.3005 | 0.219966 | 0.229575 |
| 0.488775 | NA | NA | 0.063755 | 3 | -140.014 | 286.2456 | 1.165035 | 0.143122 |
| -0.74482 | NA | 0.329563 | 0.035412 | 4 | -139.273 | 286.9135 | 1.832994 | 0.102485 |
| 0.463509 | 0.064283 | NA | NA | 3 | -140.453 | 287.1232 | 2.042694 | 0.092284 |
| -0.93731 | 0.036691 | 0.373394 | NA | 4 | -139.385 | 287.1362 | 2.055661 | 0.091687 |
| 0.416295 | 0.025417 | NA | 0.059727 | 4 | -139.993 | 288.3537 | 3.273166 | 0.049881 |
| -0.79205 | 0.018863 | 0.327811 | 0.032573 | 5 | -139.262 | 289.0795 | 3.998934 | 0.0347 |

#### **Table S20 – Model selection for grasses under control, colonisation, s.d.**

| (Intercept) | log(Nf) | log(Ng) | log(Nl + exp(-1)) | df | logLik | AICc | delta | weight |
| --- | --- | --- | --- | --- | --- | --- | --- | --- |
| -0.21245 | **NA** | **0.222441** | **NA** | **3** | **-73.7291** | **153.6764** | **0** | **0.289056** |
| 0.676614 | NA | NA | NA | 2 | -74.9566 | 154.0213 | 0.344859 | 0.243275 |
| 0.60469 | NA | NA | 0.024378 | 3 | -74.6879 | 155.594 | 1.91763 | 0.110809 |
| -0.19201 | NA | 0.212844 | 0.006073 | 4 | -73.7144 | 155.7958 | 2.119418 | 0.100174 |
| -0.20311 | -0.00468 | 0.223989 | NA | 4 | -73.7267 | 155.8204 | 2.143997 | 0.098951 |
| 0.637205 | 0.011873 | NA | NA | 3 | -74.9411 | 156.1004 | 2.42396 | 0.086025 |
| 0.617377 | -0.00445 | NA | 0.025083 | 4 | -74.686 | 157.7389 | 4.062502 | 0.037916 |
| -0.17017 | -0.00872 | 0.213654 | 0.007385 | 5 | -73.7068 | 157.9691 | 4.292715 | 0.033793 |

#### **Table S21 – Model selection for legumes under control, persistence**

| (Intercept) | I((log(size))^2) | log(Nf) | log(Ng) | log(Nl + exp(-1)) | log(size) | df | logLik | AICc | delta | weight |
| --- | --- | --- | --- | --- | --- | --- | --- | --- | --- | --- |
| **4.469706** | **-0.0812** | **NA** | **NA** | **-1.00824** | **0.336324** | **4** | **-147.4** | **302.942** | **0** | **0.148848** |
| 4.590801 | NA | NA | NA | -1.0037 | NA | 2 | -149.476 | 302.9951 | 0.053129 | 0.144946 |
| 4.696086 | NA | NA | NA | -1.08468 | 0.147559 | 3 | -148.506 | 303.0963 | 0.154277 | 0.137797 |
| 2.350687 | -0.08791 | NA | 0.547846 | -1.03201 | 0.353337 | 5 | -146.937 | 304.0891 | 1.147107 | 0.083879 |
| 2.944322 | NA | NA | 0.430459 | -1.02813 | NA | 3 | -149.182 | 304.4498 | 1.507795 | 0.070037 |
| 3.022462 | NA | NA | 0.435722 | -1.10769 | 0.148113 | 4 | -148.208 | 304.558 | 1.616038 | 0.066347 |
| 4.788265 | NA | -0.05338 | NA | -1.00992 | NA | 3 | -149.454 | 304.9923 | 2.050285 | 0.053398 |
| 4.586503 | -0.08066 | -0.03132 | NA | -1.01236 | 0.335228 | 5 | -147.392 | 304.9987 | 2.056681 | 0.053228 |
| 4.61452 | 0.004715 | NA | NA | -1.015 | NA | 3 | -149.467 | 305.0193 | 2.077276 | 0.052683 |
| 4.929037 | NA | -0.06273 | NA | -1.09248 | 0.148205 | 4 | -148.475 | 305.0914 | 2.149397 | 0.050817 |
| 2.534318 | -0.08694 | -0.08488 | 0.58428 | -1.04684 | 0.351515 | 6 | -146.884 | 306.069 | 3.127037 | 0.031168 |
| 3.145485 | NA | -0.09904 | 0.475887 | -1.04452 | NA | 4 | -149.108 | 306.3576 | 3.415593 | 0.026981 |
| 3.248649 | NA | -0.10826 | 0.483946 | -1.12613 | 0.149125 | 5 | -148.12 | 306.4535 | 3.5115 | 0.025718 |
| 2.970637 | 0.002975 | NA | 0.427385 | -1.03498 | NA | 4 | -149.179 | 306.4997 | 3.55767 | 0.025131 |
| 4.822812 | 0.005182 | -0.05564 | NA | -1.02264 | NA | 4 | -149.442 | 307.0273 | 4.085252 | 0.019304 |
| 3.179817 | 0.003572 | -0.10021 | 0.472659 | -1.05296 | NA | 5 | -149.102 | 308.419 | 5.476963 | 0.009626 |
| 1.169823 | -0.13297 | NA | NA | NA | 0.316395 | 3 | -156.985 | 320.0558 | 17.1138 | 2.86E-05 |
| 1.278027 | -0.05219 | NA | NA | NA | NA | 2 | -158.999 | 322.0403 | 19.09831 | 1.06E-05 |
| 1.015753 | -0.13345 | 0.047008 | NA | NA | 0.317812 | 4 | -156.968 | 322.0788 | 19.13677 | 1.04E-05 |
| 0.85043 | -0.13418 | NA | 0.079878 | NA | 0.318573 | 4 | -156.972 | 322.0862 | 19.14423 | 1.04E-05 |
| 1.107958 | NA | NA | NA | NA | NA | 1 | -160.276 | 322.5665 | 19.6245 | 8.15E-06 |
| 1.190618 | -0.05224 | 0.026687 | NA | NA | NA | 3 | -158.993 | 324.0717 | 21.12966 | 3.84E-06 |
| 1.267721 | -0.05221 | NA | 0.002581 | NA | NA | 3 | -158.999 | 324.083 | 21.14099 | 3.82E-06 |
| 0.75661 | -0.13443 | 0.04207 | 0.06885 | NA | 0.319544 | 5 | -156.958 | 324.1313 | 21.18925 | 3.73E-06 |
| 1.450117 | NA | NA | -0.08517 | NA | NA | 2 | -160.26 | 324.5633 | 21.6213 | 3.00E-06 |
| 1.048877 | NA | 0.018015 | NA | NA | NA | 2 | -160.274 | 324.5897 | 21.64769 | 2.96E-06 |
| 1.108159 | NA | NA | NA | NA | -0.00017 | 2 | -160.276 | 324.5948 | 21.65281 | 2.96E-06 |
| 1.209252 | -0.0522 | 0.027053 | -0.00497 | NA | NA | 4 | -158.993 | 326.1288 | 23.1868 | 1.37E-06 |
| 1.395327 | NA | 0.024294 | -0.09136 | NA | NA | 3 | -160.256 | 326.5968 | 23.65483 | 1.09E-06 |
| 1.44998 | NA | NA | -0.08537 | NA | 0.000841 | 3 | -160.26 | 326.6059 | 23.66394 | 1.08E-06 |
| 1.049198 | NA | 0.018029 | NA | NA | -0.00032 | 3 | -160.274 | 326.6324 | 23.69038 | 1.07E-06 |
| 1.395255 | NA | 0.024272 | -0.09153 | NA | 0.000706 | 4 | -160.256 | 328.654 | 25.71203 | 3.89E-07 |

#### **Table S22 – Model selection for legumes under control, expansion, mean**

| (Intercept) | I((log(size))^2) | log(Nf) | log(Ng) | log(Nl + exp(-1)) | log(size) | df | logLik | AICc | delta | weight |
| --- | --- | --- | --- | --- | --- | --- | --- | --- | --- | --- |
| 0.556798 | 0.109686 | NA | NA | NA | 0.320588 | 4 | -276.436 | 561.0624 | 0 | 0.307612 |
| 0.868072 | 0.104611 | -0.09782 | NA | NA | 0.341483 | 5 | -276.014 | 562.3144 | 1.251978 | 0.16449 |
| **0.731415** | **0.113056** | **NA** | **NA** | **-0.05933** | **0.32565** | **5** | **-276.131** | **562.5483** | **1.48589** | **0.146335** |
| 0.955207 | 0.110855 | NA | -0.09969 | NA | 0.319255 | 5 | -276.309 | 562.9051 | 1.842663 | 0.122426 |
| 1.04939 | 0.107975 | -0.099 | NA | -0.06032 | 0.346884 | 6 | -275.697 | 563.7974 | 2.735044 | 0.07836 |
| 1.155384 | 0.105797 | -0.09222 | -0.07635 | NA | 0.339266 | 6 | -275.94 | 564.2844 | 3.222012 | 0.061426 |
| 0.952357 | 0.113384 | NA | -0.06019 | -0.05268 | 0.324277 | 6 | -276.088 | 564.58 | 3.517567 | 0.052988 |
| 1.161685 | 0.108281 | -0.09653 | -0.03275 | -0.05668 | 0.345606 | 7 | -275.684 | 565.9099 | 4.847504 | 0.027251 |
| 0.682666 | 0.196126 | NA | NA | NA | NA | 3 | -280.891 | 567.8963 | 6.833918 | 0.010093 |
| 0.81717 | 0.199732 | NA | NA | -0.04519 | NA | 4 | -280.721 | 569.6321 | 8.569738 | 0.004237 |
| 1.139581 | 0.197056 | NA | -0.11448 | NA | NA | 4 | -280.731 | 569.6515 | 8.589115 | 0.004197 |
| 0.773251 | 0.196284 | -0.02774 | NA | NA | NA | 4 | -280.857 | 569.905 | 8.842638 | 0.003697 |
| 0.995021 | NA | -0.15532 | NA | NA | 0.65116 | 4 | -281.022 | 570.2347 | 9.172296 | 0.003135 |
| 0.496311 | NA | NA | NA | NA | 0.641565 | 3 | -282.065 | 570.2443 | 9.181872 | 0.00312 |
| 1.139616 | 0.199677 | NA | -0.08798 | -0.03555 | NA | 5 | -280.634 | 571.5541 | 10.49174 | 0.001621 |
| 0.908009 | 0.199892 | -0.0278 | NA | -0.04521 | NA | 5 | -280.687 | 571.6603 | 10.59793 | 0.001537 |
| 1.186313 | 0.197132 | -0.02035 | -0.10954 | NA | NA | 5 | -280.712 | 571.712 | 10.64964 | 0.001498 |
| 1.082326 | NA | -0.15675 | NA | -0.0284 | 0.658393 | 5 | -280.954 | 572.1951 | 11.13269 | 0.001176 |
| 0.567202 | NA | NA | NA | -0.02435 | 0.647688 | 4 | -282.016 | 572.2218 | 11.15939 | 0.001161 |
| 0.699925 | NA | NA | -0.05103 | NA | 0.642634 | 4 | -282.034 | 572.2576 | 11.19516 | 0.00114 |
| 1.054374 | NA | -0.1543 | -0.01569 | NA | 0.651426 | 5 | -281.019 | 572.3253 | 11.26293 | 0.001102 |
| 1.190708 | 0.199809 | -0.02225 | -0.08208 | -0.03621 | NA | 6 | -280.612 | 573.6278 | 12.56542 | 0.000575 |
| 0.696569 | NA | NA | -0.03532 | -0.02038 | 0.647431 | 5 | -282.002 | 574.2906 | 13.22823 | 0.000413 |
| 1.056402 | NA | -0.15728 | 0.007555 | -0.02927 | 0.658484 | 6 | -280.953 | 574.3106 | 13.24819 | 0.000409 |
| 0.237067 | NA | NA | NA | 0.312548 | NA | 3 | -337.501 | 681.1158 | 120.0534 | 2.62E-27 |
| 0.635293 | NA | NA | -0.1086 | 0.324325 | NA | 4 | -337.423 | 683.0357 | 121.9733 | 1.00E-27 |
| 0.16896 | NA | 0.020947 | NA | 0.312346 | NA | 4 | -337.49 | 683.1696 | 122.1072 | 9.39E-28 |
| 0.569671 | NA | 0.028768 | -0.11622 | 0.324873 | NA | 5 | -337.401 | 685.0897 | 124.0273 | 3.60E-28 |
| 1.238067 | NA | NA | NA | NA | NA | 2 | -342.975 | 690.0063 | 128.9439 | 3.08E-29 |
| 0.565982 | NA | NA | 0.167432 | NA | NA | 3 | -342.78 | 691.6728 | 130.6104 | 1.34E-29 |
| 1.147625 | NA | 0.027555 | NA | NA | NA | 3 | -342.956 | 692.0257 | 130.9633 | 1.12E-29 |
| 0.528996 | NA | 0.016185 | 0.163411 | NA | NA | 4 | -342.773 | 693.7367 | 132.6743 | 4.77E-30 |

#### **Table S23 – Model selection for legumes under control, expansion, s.d.**

| (Intercept) | I((log(size))^2) | log(Nf) | log(Ng) | log(Nl + exp(-1)) | log(size) | df | logLik | AICc | delta | weight |
| --- | --- | --- | --- | --- | --- | --- | --- | --- | --- | --- |
| **0.598447** | **NA** | **0.087334** | **NA** | **-0.07379** | **0.05569** | **5** | **-154.569** | **319.4251** | **0** | **0.077678** |
| 0.885448 | NA | NA | NA | -0.07605 | 0.061654 | 4 | -155.637 | 319.4642 | 0.039061 | 0.076175 |
| 0.847533 | -0.0261 | NA | NA | -0.06798 | 0.13601 | 5 | -154.643 | 319.5732 | 0.14808 | 0.072134 |
| 0.380286 | NA | 0.101413 | NA | NA | NA | 3 | -156.905 | 319.9245 | 0.499394 | 0.060514 |
| 0.647462 | -0.02996 | NA | NA | NA | 0.13021 | 4 | -155.879 | 319.948 | 0.522928 | 0.059806 |
| 0.605228 | -0.02223 | 0.075444 | NA | -0.06722 | 0.119829 | 6 | -153.862 | 320.1276 | 0.70253 | 0.054669 |
| 0.371638 | NA | 0.09105 | NA | NA | 0.036899 | 4 | -156.049 | 320.2878 | 0.862724 | 0.050461 |
| 0.403167 | -0.02598 | 0.076768 | NA | NA | 0.113811 | 5 | -155.079 | 320.445 | 1.019925 | 0.046647 |
| 0.663986 | NA | NA | NA | NA | 0.042524 | 3 | -157.195 | 320.5028 | 1.077707 | 0.045319 |
| 0.713151 | NA | NA | NA | NA | NA | 2 | -158.339 | 320.7338 | 1.308709 | 0.040375 |
| 0.521189 | NA | 0.102365 | NA | -0.04497 | NA | 4 | -156.28 | 320.7504 | 1.325263 | 0.040042 |
| 0.801259 | NA | NA | 0.022983 | -0.07863 | 0.061822 | 5 | -155.618 | 321.5227 | 2.097562 | 0.027216 |
| 0.601322 | NA | 0.087393 | -0.00084 | -0.0737 | 0.05568 | 6 | -154.569 | 321.5418 | 2.116715 | 0.026956 |
| 0.854022 | NA | NA | NA | -0.04399 | NA | 3 | -157.748 | 321.6101 | 2.184985 | 0.026052 |
| 0.742017 | -0.02626 | NA | 0.028743 | -0.07115 | 0.136665 | 6 | -154.613 | 321.6299 | 2.204766 | 0.025795 |
| 0.371565 | 0.004572 | 0.100125 | NA | NA | NA | 4 | -156.761 | 321.7121 | 2.287007 | 0.024756 |
| 0.556389 | 0.009521 | 0.100043 | NA | -0.062 | NA | 5 | -155.738 | 321.764 | 2.338938 | 0.024121 |
| 0.565642 | NA | 0.10482 | -0.04896 | NA | NA | 4 | -156.813 | 321.816 | 2.390891 | 0.023503 |
| 0.745867 | -0.02968 | NA | -0.02462 | NA | 0.129881 | 5 | -155.855 | 321.9971 | 2.572006 | 0.021468 |
| 0.596213 | NA | 0.0949 | -0.05938 | NA | 0.037905 | 5 | -155.912 | 322.111 | 2.685889 | 0.02028 |
| 0.579638 | -0.0223 | 0.07488 | 0.007464 | -0.06805 | 0.12012 | 7 | -153.86 | 322.2608 | 2.835754 | 0.018816 |
| 0.572073 | -0.02528 | 0.080061 | -0.04489 | NA | 0.112507 | 6 | -155.001 | 322.4056 | 2.98049 | 0.017502 |
| 0.698585 | 0.005144 | NA | NA | NA | NA | 3 | -158.158 | 322.4292 | 3.004123 | 0.017297 |
| 0.814205 | NA | NA | -0.03765 | NA | 0.043312 | 4 | -157.139 | 322.4692 | 3.044093 | 0.016954 |
| 0.883349 | 0.010097 | NA | NA | -0.06207 | NA | 4 | -157.146 | 322.4832 | 3.0581 | 0.016836 |
| 0.805178 | NA | NA | -0.02293 | NA | NA | 3 | -158.318 | 322.7502 | 3.325083 | 0.014732 |
| 0.560165 | NA | 0.103125 | -0.0113 | -0.04375 | NA | 5 | -156.275 | 322.838 | 3.412863 | 0.014099 |
| 0.582329 | 0.005005 | 0.103892 | -0.05589 | NA | NA | 5 | -156.641 | 323.5696 | 4.144472 | 0.00978 |
| 0.795408 | NA | NA | 0.015985 | -0.04572 | NA | 4 | -157.739 | 323.6687 | 4.24363 | 0.009307 |
| 0.589725 | 0.009511 | 0.100697 | -0.00968 | -0.06094 | NA | 6 | -155.735 | 323.8742 | 4.449068 | 0.008398 |
| 0.820875 | 0.005393 | NA | -0.03064 | NA | NA | 4 | -158.122 | 324.4338 | 5.008756 | 0.006348 |
| 0.820937 | 0.010108 | NA | 0.017029 | -0.06394 | NA | 5 | -157.136 | 324.5592 | 5.134064 | 0.005963 |

#### **Table S24 – Model selection for legumes under control, colonisation, mean**

| (Intercept) | log(Nf) | log(Ng) | log(Nl + exp(-1)) | df | logLik | AICc | delta | weight |
| --- | --- | --- | --- | --- | --- | --- | --- | --- |
| **0.288878** | **NA** | **NA** | **NA** | **2** | **-26.1461** | **56.67934** | **0** | **0.302181** |
| -0.27656 | 0.181088 | NA | NA | 3 | -25.4004 | 57.60082 | 0.921482 | 0.19062 |
| -1.18743 | NA | 0.367064 | NA | 3 | -25.4031 | 57.6062 | 0.926861 | 0.190108 |
| 0.224877 | NA | NA | 0.026346 | 3 | -25.9979 | 58.79574 | 2.1164 | 0.104881 |
| -1.28303 | 0.139568 | 0.282479 | NA | 4 | -24.9856 | 59.35051 | 2.671168 | 0.079475 |
| -1.53865 | NA | 0.468912 | -0.02405 | 4 | -25.333 | 60.04524 | 3.365902 | 0.056153 |
| -0.27031 | 0.173574 | NA | 0.007085 | 4 | -25.3906 | 60.16045 | 3.481108 | 0.05301 |
| -1.84008 | 0.157509 | 0.429581 | -0.0373 | 5 | -24.8192 | 61.78125 | 5.101907 | 0.023572 |

#### **Table S25 – Model selection for legumes under control, colonisation, s.d.**

| (Intercept) | log(Nf) | log(Ng) | log(Nl + exp(-1)) | df | logLik | AICc | delta | weight |
| --- | --- | --- | --- | --- | --- | --- | --- | --- |
| **0.407827** | **NA** | **NA** | **NA** | **2** | **-10.1296** | **24.64636** | **0** | **0.300418** |
| 0.046916 | 0.115586 | NA | NA | 3 | -9.34943 | 25.49886 | 0.852498 | 0.196159 |
| -0.50133 | NA | 0.226049 | NA | 3 | -9.40715 | 25.61429 | 0.967929 | 0.185158 |
| 0.363751 | NA | NA | 0.018144 | 3 | -9.94907 | 26.69814 | 2.051775 | 0.107693 |
| -0.56326 | 0.090415 | 0.171253 | NA | 4 | -8.95777 | 27.29485 | 2.648483 | 0.079913 |
| 0.052236 | 0.109193 | NA | 0.006027 | 4 | -9.33111 | 28.04154 | 3.395175 | 0.055014 |
| -0.66616 | NA | 0.273847 | -0.01129 | 4 | -9.3676 | 28.11451 | 3.468147 | 0.053043 |
| -0.85731 | 0.099885 | 0.248905 | -0.01969 | 5 | -8.8389 | 29.82065 | 5.174285 | 0.022602 |

#### **Table S26 – Model selection for forbs under control, persistence**

| (Intercept) | I((log(size))^2) | log(Nf) | log(Ng) | log(Nl + exp(-1)) | log(size) | df | logLik | AICc | delta | weight |
| --- | --- | --- | --- | --- | --- | --- | --- | --- | --- | --- |
| 4.650234 | NA | -0.59584 | -0.5543 | NA | 0.470098 | 4 | -525.104 | 1058.255 | 0 | 0.433913 |
| **4.460571** | **NA** | **-0.59059** | **-0.48189** | **-0.03808** | **0.469091** | **5** | **-524.966** | **1060.003** | **1.748109** | **0.181053** |
| 4.722483 | 0.013908 | -0.61241 | -0.56113 | NA | 0.463647 | 5 | -525.062 | 1060.194 | 1.939047 | 0.164567 |
| 2.911628 | NA | -0.63037 | NA | -0.11546 | 0.477942 | 4 | -526.814 | 1061.676 | 3.420861 | 0.078446 |
| 4.532739 | 0.013672 | -0.60692 | -0.48899 | -0.0379 | 0.462763 | 6 | -524.926 | 1061.949 | 3.694643 | 0.06841 |
| 2.729579 | NA | -0.68079 | NA | NA | 0.487937 | 3 | -528.706 | 1063.44 | 5.185409 | 0.032464 |
| 2.928845 | 0.004815 | -0.63633 | NA | -0.11577 | 0.475786 | 5 | -526.809 | 1063.689 | 5.433993 | 0.02867 |
| 2.730654 | 0.000312 | -0.68118 | NA | NA | 0.487799 | 4 | -528.706 | 1065.459 | 7.204069 | 0.011832 |
| 3.44851 | NA | NA | -0.74213 | NA | 0.396921 | 3 | -533.665 | 1073.358 | 15.10329 | 0.000228 |
| 3.294233 | -0.05516 | NA | -0.69352 | NA | 0.426682 | 4 | -532.857 | 1073.76 | 15.50504 | 0.000186 |
| 3.191828 | NA | NA | -0.63595 | -0.05458 | 0.396056 | 4 | -533.37 | 1074.787 | 16.53201 | 0.000112 |
| 3.041124 | -0.0548 | NA | -0.58892 | -0.05378 | 0.425701 | 5 | -532.571 | 1075.212 | 16.95741 | 9.02E-05 |
| 0.996265 | -0.06954 | NA | NA | -0.14782 | 0.440269 | 4 | -535.368 | 1078.782 | 20.52716 | 1.51E-05 |
| 0.976146 | NA | NA | NA | -0.15769 | 0.404442 | 3 | -536.694 | 1079.417 | 21.16205 | 1.10E-05 |
| 0.559717 | -0.08136 | NA | NA | NA | 0.454257 | 3 | -538.666 | 1083.359 | 25.10441 | 1.53E-06 |
| 0.500804 | NA | NA | NA | NA | 0.414614 | 2 | -540.503 | 1085.021 | 26.7657 | 6.69E-07 |
| 5.024493 | 0.069916 | -0.52309 | -0.68128 | NA | NA | 4 | -539.909 | 1087.865 | 29.61065 | 1.61E-07 |
| 4.620948 | NA | -0.41321 | -0.65345 | NA | NA | 3 | -541.311 | 1088.651 | 30.39599 | 1.09E-07 |
| 4.807918 | 0.069466 | -0.51664 | -0.59875 | -0.04377 | NA | 5 | -539.724 | 1089.517 | 31.26242 | 7.06E-08 |
| 4.396644 | NA | -0.40727 | -0.56721 | -0.04571 | NA | 4 | -541.108 | 1090.262 | 32.00713 | 4.87E-08 |
| 2.841024 | 0.061904 | -0.55083 | NA | -0.13915 | NA | 4 | -542.632 | 1093.31 | 35.05557 | 1.06E-08 |
| 2.567309 | NA | -0.45163 | NA | -0.13653 | NA | 3 | -543.746 | 1093.52 | 35.26493 | 9.54E-09 |
| 3.777541 | NA | NA | -0.78745 | NA | NA | 2 | -545.917 | 1095.847 | 37.59249 | 2.98E-09 |
| 2.604686 | 0.05922 | -0.6044 | NA | NA | NA | 3 | -545.451 | 1096.93 | 38.67479 | 1.73E-09 |
| 2.347142 | NA | -0.50853 | NA | NA | NA | 2 | -546.475 | 1096.964 | 38.70908 | 1.71E-09 |
| 3.507683 | NA | NA | -0.67541 | -0.05801 | NA | 3 | -545.579 | 1097.187 | 38.93174 | 1.53E-09 |
| 3.796617 | 0.007671 | NA | -0.79427 | NA | NA | 3 | -545.896 | 1097.82 | 39.56481 | 1.11E-09 |
| 3.526945 | 0.007938 | NA | -0.68224 | -0.05814 | NA | 4 | -545.557 | 1099.161 | 40.90576 | 5.69E-10 |
| 1.158479 | NA | NA | NA | -0.16829 | NA | 2 | -549.422 | 1102.858 | 44.6029 | 8.95E-11 |
| 1.161762 | -0.00542 | NA | NA | -0.1675 | NA | 3 | -549.411 | 1104.85 | 46.59559 | 3.31E-11 |
| 0.653384 | NA | NA | NA | NA | NA | 1 | -553.837 | 1109.678 | 51.42303 | 2.96E-12 |
| 0.670144 | -0.01606 | NA | NA | NA | NA | 2 | -553.742 | 1111.498 | 53.24326 | 1.19E-12 |

#### **Table S27 – Model selection for forbs under control, expansion, mean**

| (Intercept) | I((log(size))^2) | log(Nf) | log(Ng) | log(Nl + exp(-1)) | log(size) | df | logLik | AICc | delta | weight |
| --- | --- | --- | --- | --- | --- | --- | --- | --- | --- | --- |
| **-0.416982209** | **0.101694** | **NA** | **0.236052** | **-0.05655** | **0.164809** | **6** | **-532.917** | **1077.985** | **0** | **0.413991** |
| -0.605014525 | 0.094735 | 0.071291 | 0.228364 | -0.05832 | 0.160956 | 7 | -532.109 | 1078.418 | 0.433405 | 0.333333 |
| -0.097964745 | 0.101192 | NA | 0.11294 | NA | 0.167557 | 5 | -535.67 | 1081.447 | 3.461912 | 0.073324 |
| -0.25628126 | 0.094989 | 0.063404 | 0.102663 | NA | 0.164207 | 6 | -535.035 | 1082.219 | 4.234458 | 0.04983 |
| 0.341808137 | 0.106746 | NA | NA | NA | 0.160451 | 4 | -537.185 | 1082.442 | 4.456902 | 0.044585 |
| 0.104954636 | 0.098735 | 0.075718 | NA | NA | 0.157222 | 5 | -536.266 | 1082.638 | 4.653755 | 0.040406 |
| 0.132471027 | 0.100009 | 0.082471 | NA | -0.01715 | 0.153751 | 6 | -535.856 | 1083.861 | 5.876428 | 0.021925 |
| 0.380019072 | 0.108308 | NA | NA | -0.01347 | 0.15795 | 5 | -536.928 | 1083.964 | 5.979174 | 0.020827 |
| -0.530874612 | 0.141802 | 0.086731 | 0.208513 | -0.06126 | NA | 6 | -539.34 | 1090.83 | 12.84513 | 0.000673 |
| -0.298755617 | 0.151689 | NA | 0.217333 | -0.05917 | NA | 5 | -540.512 | 1091.13 | 13.14566 | 0.000579 |
| 0.104541718 | 0.14434 | 0.087487 | NA | NA | NA | 4 | -543.152 | 1094.375 | 16.39066 | 0.000114 |
| 0.379079065 | 0.154711 | NA | NA | NA | NA | 3 | -544.353 | 1094.749 | 16.76473 | 9.47E-05 |
| 0.142090511 | 0.144704 | 0.096345 | NA | -0.0234 | NA | 5 | -542.4 | 1094.908 | 16.92318 | 8.75E-05 |
| 0.037386158 | 0.152037 | NA | 0.088082 | NA | NA | 4 | -543.447 | 1094.966 | 16.98085 | 8.50E-05 |
| -0.16265271 | 0.143068 | 0.078766 | 0.075932 | NA | NA | 5 | -542.488 | 1095.083 | 17.09833 | 8.02E-05 |
| 0.432928102 | 0.155877 | NA | NA | -0.01928 | NA | 4 | -543.837 | 1095.745 | 17.76046 | 5.76E-05 |
| -0.989443626 | NA | 0.149084 | 0.268905 | -0.05896 | 0.292279 | 6 | -543.889 | 1099.927 | 21.94261 | 7.12E-06 |
| -0.637919204 | NA | 0.141321 | 0.141932 | NA | 0.295922 | 5 | -546.758 | 1103.622 | 25.63729 | 1.12E-06 |
| -0.626159489 | NA | NA | 0.293153 | -0.05503 | 0.32297 | 5 | -547.557 | 1105.222 | 27.23693 | 5.04E-07 |
| -0.151982789 | NA | 0.162919 | NA | NA | 0.293408 | 4 | -549.046 | 1106.163 | 28.17784 | 3.15E-07 |
| -0.137816413 | NA | 0.167531 | NA | -0.01004 | 0.292404 | 5 | -548.911 | 1107.928 | 29.94361 | 1.30E-07 |
| -0.314649482 | NA | NA | 0.173051 | NA | 0.324884 | 4 | -550.035 | 1108.14 | 30.15553 | 1.17E-07 |
| 0.360697391 | NA | NA | NA | NA | 0.327297 | 3 | -553.506 | 1113.055 | 35.07051 | 1.00E-08 |
| 0.362292265 | NA | NA | NA | -0.00056 | 0.327294 | 4 | -553.506 | 1115.083 | 37.09821 | 3.64E-09 |
| -1.346274854 | NA | 0.314258 | 0.269791 | -0.06889 | NA | 5 | -580.395 | 1170.897 | 92.91194 | 2.76E-21 |
| -0.939900112 | NA | 0.307579 | 0.121138 | NA | NA | 4 | -583.842 | 1175.755 | 97.76999 | 2.44E-22 |
| -0.522448905 | NA | 0.324829 | NA | NA | NA | 3 | -585.308 | 1176.659 | 98.6742 | 1.55E-22 |
| -0.491994744 | NA | 0.332837 | NA | -0.01981 | NA | 4 | -584.844 | 1177.759 | 99.77394 | 8.94E-23 |
| -0.569063433 | NA | NA | 0.327855 | -0.062 | NA | 4 | -596.266 | 1200.603 | 122.6184 | 9.79E-28 |
| -0.217462392 | NA | NA | 0.192675 | NA | NA | 3 | -598.915 | 1203.873 | 125.8881 | 1.91E-28 |
| 0.535904101 | NA | NA | NA | NA | NA | 2 | -602.539 | 1209.1 | 131.1152 | 1.40E-29 |
| 0.539005176 | NA | NA | NA | -0.00109 | NA | 3 | -602.538 | 1211.119 | 133.1339 | 5.10E-30 |

#### **Table S28 – Model selection for forbs under control, expansion, s.d.**

| (Intercept) | I((log(size))^2) | log(Nf) | log(Ng) | log(Nl + exp(-1)) | log(size) | df | logLik | AICc | delta | weight |
| --- | --- | --- | --- | --- | --- | --- | --- | --- | --- | --- |
| **-0.20337** | **0.029425** | **0.085946** | **0.124428** | **-0.04128** | **0.04593** | **7** | **-198.21** | **410.6213** | **0** | **0.562968** |
| -0.18221 | 0.042855 | 0.090352 | 0.118764 | -0.04212 | NA | 6 | -200.14 | 412.4309 | 1.809582 | 0.227792 |
| 0.198464 | 0.032298 | 0.092037 | NA | -0.01885 | 0.042004 | 6 | -201.823 | 415.7963 | 5.175049 | 0.042339 |
| -0.32277 | NA | 0.110108 | 0.13702 | -0.04148 | 0.086719 | 6 | -201.953 | 416.0569 | 5.435597 | 0.037167 |
| 0.023317 | 0.037814 | NA | 0.133696 | -0.03914 | 0.050576 | 6 | -202.006 | 416.1617 | 5.540439 | 0.035269 |
| 0.201092 | 0.044509 | 0.095828 | NA | -0.02056 | NA | 5 | -203.424 | 416.955 | 6.333724 | 0.023721 |
| 0.168222 | 0.030898 | 0.084615 | NA | NA | 0.045819 | 5 | -203.429 | 416.9657 | 6.344371 | 0.023595 |
| 0.043477 | 0.029604 | 0.080363 | 0.035452 | NA | 0.048231 | 6 | -202.955 | 418.06 | 7.438761 | 0.013651 |
| 0.168102 | 0.044188 | 0.088045 | NA | NA | NA | 4 | -205.338 | 418.7474 | 8.126091 | 0.009681 |
| 0.059598 | 0.053156 | NA | 0.127952 | -0.03995 | NA | 5 | -204.325 | 418.7576 | 8.136298 | 0.009632 |
| 0.070978 | 0.043726 | 0.084875 | 0.027601 | NA | NA | 5 | -205.049 | 420.2058 | 9.584512 | 0.004669 |
| 0.432909 | 0.03985 | NA | NA | NA | 0.049427 | 4 | -207.126 | 422.3227 | 11.70142 | 0.00162 |
| 0.474727 | 0.04156 | NA | NA | -0.01475 | 0.046691 | 5 | -206.139 | 422.3857 | 11.76441 | 0.00157 |
| 0.24414 | 0.037466 | NA | 0.048478 | NA | 0.052478 | 5 | -206.232 | 422.5716 | 11.95031 | 0.001431 |
| 0.111174 | NA | 0.119508 | NA | -0.01655 | 0.086782 | 5 | -206.314 | 422.7355 | 12.1142 | 0.001318 |
| 0.087817 | NA | 0.111904 | NA | NA | 0.088436 | 4 | -207.541 | 423.1526 | 12.53134 | 0.00107 |
| -0.07546 | NA | 0.104647 | 0.047691 | NA | 0.089281 | 5 | -206.681 | 423.4693 | 12.84806 | 0.000913 |
| 0.490367 | 0.055621 | NA | NA | -0.01646 | NA | 4 | -208.095 | 424.2617 | 13.64043 | 0.000614 |
| 0.44439 | 0.054626 | NA | NA | NA | NA | 3 | -209.324 | 424.6908 | 14.06956 | 0.000496 |
| 0.286531 | 0.05339 | NA | 0.040693 | NA | NA | 4 | -208.694 | 425.4589 | 14.83758 | 0.000338 |
| -0.05446 | NA | NA | 0.154929 | -0.03858 | 0.109385 | 5 | -208.603 | 427.3133 | 16.69207 | 0.000134 |
| 0.163913 | NA | NA | 0.070735 | NA | 0.110728 | 4 | -212.616 | 433.304 | 22.68268 | 6.68E-06 |
| 0.439961 | NA | NA | NA | NA | 0.111714 | 3 | -214.529 | 435.1 | 24.4787 | 2.72E-06 |
| 0.467925 | NA | NA | NA | -0.00979 | 0.111671 | 4 | -214.1 | 436.2703 | 25.64906 | 1.52E-06 |
| -0.42864 | NA | 0.159115 | 0.137283 | -0.04443 | NA | 5 | -213.185 | 436.4769 | 25.85557 | 1.37E-06 |
| 0.006058 | NA | 0.168569 | NA | -0.01945 | NA | 4 | -217.394 | 442.8584 | 32.23713 | 5.63E-08 |
| -0.02385 | NA | 0.160705 | NA | NA | NA | 3 | -219.027 | 444.0963 | 33.47505 | 3.03E-08 |
| -0.16657 | NA | 0.154808 | 0.041417 | NA | NA | 4 | -218.404 | 444.8788 | 34.25752 | 2.05E-08 |
| -0.03512 | NA | NA | 0.166682 | -0.04094 | NA | 4 | -228.07 | 464.211 | 53.58975 | 1.30E-12 |
| 0.197037 | NA | NA | 0.077423 | NA | NA | 3 | -232.291 | 470.6244 | 60.00314 | 5.26E-14 |
| 0.499762 | NA | NA | NA | NA | NA | 2 | -234.429 | 472.8798 | 62.25848 | 1.70E-14 |
| 0.528218 | NA | NA | NA | -0.00997 | NA | 3 | -234.014 | 474.0714 | 63.45011 | 9.39E-15 |

#### **Table S29 – Model selection for forbs under control, colonisation, mean**

| (Intercept) | log(Nf) | log(Ng) | log(Nl + exp(-1)) | df | logLik | AICc | delta | weight |
| --- | --- | --- | --- | --- | --- | --- | --- | --- |
| -1.20115 | 0.16742 | 0.230141 | NA | 4 | -167.31 | 342.7867 | 0 | 0.63623 |
| **-1.25773** | **0.170522** | **0.249966** | **-0.011** | **5** | **-167.215** | **344.6791** | **1.892403** | **0.246993** |
| -0.34533 | 0.18402 | NA | NA | 3 | -170.524 | 347.1466 | 4.359885 | 0.071925 |
| -0.37172 | 0.177042 | NA | 0.016942 | 4 | -170.246 | 348.6586 | 5.871924 | 0.033771 |
| -0.81192 | NA | 0.269142 | NA | 3 | -172.753 | 351.6044 | 8.817758 | 0.007742 |
| -0.80822 | NA | 0.267603 | 0.000824 | 4 | -172.752 | 353.6702 | 10.88351 | 0.002756 |
| 0.251939 | NA | NA | NA | 2 | -177.011 | 358.0709 | 15.28427 | 0.000305 |
| 0.161321 | NA | NA | 0.031306 | 3 | -176.079 | 358.2568 | 15.47009 | 0.000278 |

#### **Table S30 – Model selection for forbs under control, colonisation, s.d.**

| (Intercept) | log(Nf) | log(Ng) | log(Nl + exp(-1)) | df | logLik | AICc | delta | weight |
| --- | --- | --- | --- | --- | --- | --- | --- | --- |
| -1.06334 | **0.184727** | **0.207695** | **NA** | **4** | **-39.2382** | **86.64235** | **0** | **0.73006** |
| -1.08409 | 0.185865 | 0.214968 | -0.00404 | 5 | -39.2017 | 88.65335 | 2.011003 | 0.267101 |
| -0.32213 | 0.191472 | NA | 0.019996 | 4 | -45.4668 | 99.09959 | 12.45724 | 0.00144 |
| -0.29098 | 0.199708 | NA | NA | 3 | -46.529 | 99.15721 | 12.51486 | 0.001399 |
| -0.63388 | NA | 0.250728 | NA | 3 | -57.0706 | 120.2404 | 33.59803 | 3.70E-08 |
| -0.59414 | NA | 0.234191 | 0.008854 | 4 | -56.9155 | 121.9969 | 35.35457 | 1.54E-08 |
| 0.254353 | NA | NA | 0.035531 | 3 | -63.3664 | 132.8319 | 46.18958 | 6.81E-11 |
| 0.357199 | NA | NA | NA | 2 | -66.3429 | 136.7352 | 50.09281 | 9.68E-12 |

#### **Table S31 – Model selection for grasses under drought, persistence**

| (Intercept) | I((log(size))^2) | log(Nf) | log(Ng) | log(Nl + exp(-1)) | log(size) | df | logLik | AICc | delta | weight |
| --- | --- | --- | --- | --- | --- | --- | --- | --- | --- | --- |
| 4.650234 | NA | -0.59584 | -0.5543 | NA | 0.470098 | 4 | -525.104 | 1058.255 | 0 | 0.433913 |
| **4.460571** | **NA** | **-0.59059** | **-0.48189** | **-0.03808** | **0.469091** | **5** | **-524.966** | **1060.003** | **1.748109** | **0.181053** |
| 4.722483 | 0.013908 | -0.61241 | -0.56113 | NA | 0.463647 | 5 | -525.062 | 1060.194 | 1.939047 | 0.164567 |
| 2.911628 | NA | -0.63037 | NA | -0.11546 | 0.477942 | 4 | -526.814 | 1061.676 | 3.420861 | 0.078446 |
| 4.532739 | 0.013672 | -0.60692 | -0.48899 | -0.0379 | 0.462763 | 6 | -524.926 | 1061.949 | 3.694643 | 0.06841 |
| 2.729579 | NA | -0.68079 | NA | NA | 0.487937 | 3 | -528.706 | 1063.44 | 5.185409 | 0.032464 |
| 2.928845 | 0.004815 | -0.63633 | NA | -0.11577 | 0.475786 | 5 | -526.809 | 1063.689 | 5.433993 | 0.02867 |
| 2.730654 | 0.000312 | -0.68118 | NA | NA | 0.487799 | 4 | -528.706 | 1065.459 | 7.204069 | 0.011832 |
| 3.44851 | NA | NA | -0.74213 | NA | 0.396921 | 3 | -533.665 | 1073.358 | 15.10329 | 0.000228 |
| 3.294233 | -0.05516 | NA | -0.69352 | NA | 0.426682 | 4 | -532.857 | 1073.76 | 15.50504 | 0.000186 |
| 3.191828 | NA | NA | -0.63595 | -0.05458 | 0.396056 | 4 | -533.37 | 1074.787 | 16.53201 | 0.000112 |
| 3.041124 | -0.0548 | NA | -0.58892 | -0.05378 | 0.425701 | 5 | -532.571 | 1075.212 | 16.95741 | 9.02E-05 |
| 0.996265 | -0.06954 | NA | NA | -0.14782 | 0.440269 | 4 | -535.368 | 1078.782 | 20.52716 | 1.51E-05 |
| 0.976146 | NA | NA | NA | -0.15769 | 0.404442 | 3 | -536.694 | 1079.417 | 21.16205 | 1.10E-05 |
| 0.559717 | -0.08136 | NA | NA | NA | 0.454257 | 3 | -538.666 | 1083.359 | 25.10441 | 1.53E-06 |
| 0.500804 | NA | NA | NA | NA | 0.414614 | 2 | -540.503 | 1085.021 | 26.7657 | 6.69E-07 |
| 5.024493 | 0.069916 | -0.52309 | -0.68128 | NA | NA | 4 | -539.909 | 1087.865 | 29.61065 | 1.61E-07 |
| 4.620948 | NA | -0.41321 | -0.65345 | NA | NA | 3 | -541.311 | 1088.651 | 30.39599 | 1.09E-07 |
| 4.807918 | 0.069466 | -0.51664 | -0.59875 | -0.04377 | NA | 5 | -539.724 | 1089.517 | 31.26242 | 7.06E-08 |
| 4.396644 | NA | -0.40727 | -0.56721 | -0.04571 | NA | 4 | -541.108 | 1090.262 | 32.00713 | 4.87E-08 |
| 2.841024 | 0.061904 | -0.55083 | NA | -0.13915 | NA | 4 | -542.632 | 1093.31 | 35.05557 | 1.06E-08 |
| 2.567309 | NA | -0.45163 | NA | -0.13653 | NA | 3 | -543.746 | 1093.52 | 35.26493 | 9.54E-09 |
| 3.777541 | NA | NA | -0.78745 | NA | NA | 2 | -545.917 | 1095.847 | 37.59249 | 2.98E-09 |
| 2.604686 | 0.05922 | -0.6044 | NA | NA | NA | 3 | -545.451 | 1096.93 | 38.67479 | 1.73E-09 |
| 2.347142 | NA | -0.50853 | NA | NA | NA | 2 | -546.475 | 1096.964 | 38.70908 | 1.71E-09 |
| 3.507683 | NA | NA | -0.67541 | -0.05801 | NA | 3 | -545.579 | 1097.187 | 38.93174 | 1.53E-09 |
| 3.796617 | 0.007671 | NA | -0.79427 | NA | NA | 3 | -545.896 | 1097.82 | 39.56481 | 1.11E-09 |
| 3.526945 | 0.007938 | NA | -0.68224 | -0.05814 | NA | 4 | -545.557 | 1099.161 | 40.90576 | 5.69E-10 |
| 1.158479 | NA | NA | NA | -0.16829 | NA | 2 | -549.422 | 1102.858 | 44.6029 | 8.95E-11 |
| 1.161762 | -0.00542 | NA | NA | -0.1675 | NA | 3 | -549.411 | 1104.85 | 46.59559 | 3.31E-11 |
| 0.653384 | NA | NA | NA | NA | NA | 1 | -553.837 | 1109.678 | 51.42303 | 2.96E-12 |
| 0.670144 | -0.01606 | NA | NA | NA | NA | 2 | -553.742 | 1111.498 | 53.24326 | 1.19E-12 |

#### **Table S32 – Model selection for grasses under drought, expansion, mean**

| (Intercept) | I((log(size))^2) | log(Nf) | log(Ng) | log(Nl + exp(-1)) | log(size) | df | logLik | AICc | delta | weight |
| --- | --- | --- | --- | --- | --- | --- | --- | --- | --- | --- |
| **-0.41698** | **0.101694** | **NA** | **0.236052** | **-0.05655** | **0.164809** | **6** | **-532.917** | **1077.985** | **0** | **0.413991** |
| -0.60501 | 0.094735 | 0.071291 | 0.228364 | -0.05832 | 0.160956 | 7 | -532.109 | 1078.418 | 0.433405 | 0.333333 |
| -0.09796 | 0.101192 | NA | 0.11294 | NA | 0.167557 | 5 | -535.67 | 1081.447 | 3.461912 | 0.073324 |
| -0.25628 | 0.094989 | 0.063404 | 0.102663 | NA | 0.164207 | 6 | -535.035 | 1082.219 | 4.234458 | 0.04983 |
| 0.341808 | 0.106746 | NA | NA | NA | 0.160451 | 4 | -537.185 | 1082.442 | 4.456902 | 0.044585 |
| 0.104955 | 0.098735 | 0.075718 | NA | NA | 0.157222 | 5 | -536.266 | 1082.638 | 4.653755 | 0.040406 |
| 0.132471 | 0.100009 | 0.082471 | NA | -0.01715 | 0.153751 | 6 | -535.856 | 1083.861 | 5.876428 | 0.021925 |
| 0.380019 | 0.108308 | NA | NA | -0.01347 | 0.15795 | 5 | -536.928 | 1083.964 | 5.979174 | 0.020827 |
| -0.53087 | 0.141802 | 0.086731 | 0.208513 | -0.06126 | NA | 6 | -539.34 | 1090.83 | 12.84513 | 0.000673 |
| -0.29876 | 0.151689 | NA | 0.217333 | -0.05917 | NA | 5 | -540.512 | 1091.13 | 13.14566 | 0.000579 |
| 0.104542 | 0.14434 | 0.087487 | NA | NA | NA | 4 | -543.152 | 1094.375 | 16.39066 | 0.000114 |
| 0.379079 | 0.154711 | NA | NA | NA | NA | 3 | -544.353 | 1094.749 | 16.76473 | 9.47E-05 |
| 0.142091 | 0.144704 | 0.096345 | NA | -0.0234 | NA | 5 | -542.4 | 1094.908 | 16.92318 | 8.75E-05 |
| 0.037386 | 0.152037 | NA | 0.088082 | NA | NA | 4 | -543.447 | 1094.966 | 16.98085 | 8.50E-05 |
| -0.16265 | 0.143068 | 0.078766 | 0.075932 | NA | NA | 5 | -542.488 | 1095.083 | 17.09833 | 8.02E-05 |
| 0.432928 | 0.155877 | NA | NA | -0.01928 | NA | 4 | -543.837 | 1095.745 | 17.76046 | 5.76E-05 |
| -0.98944 | NA | 0.149084 | 0.268905 | -0.05896 | 0.292279 | 6 | -543.889 | 1099.927 | 21.94261 | 7.12E-06 |
| -0.63792 | NA | 0.141321 | 0.141932 | NA | 0.295922 | 5 | -546.758 | 1103.622 | 25.63729 | 1.12E-06 |
| -0.62616 | NA | NA | 0.293153 | -0.05503 | 0.32297 | 5 | -547.557 | 1105.222 | 27.23693 | 5.04E-07 |
| -0.15198 | NA | 0.162919 | NA | NA | 0.293408 | 4 | -549.046 | 1106.163 | 28.17784 | 3.15E-07 |
| -0.13782 | NA | 0.167531 | NA | -0.01004 | 0.292404 | 5 | -548.911 | 1107.928 | 29.94361 | 1.30E-07 |
| -0.31465 | NA | NA | 0.173051 | NA | 0.324884 | 4 | -550.035 | 1108.14 | 30.15553 | 1.17E-07 |
| 0.360697 | NA | NA | NA | NA | 0.327297 | 3 | -553.506 | 1113.055 | 35.07051 | 1.00E-08 |
| 0.362292 | NA | NA | NA | -0.00056 | 0.327294 | 4 | -553.506 | 1115.083 | 37.09821 | 3.64E-09 |
| -1.34627 | NA | 0.314258 | 0.269791 | -0.06889 | NA | 5 | -580.395 | 1170.897 | 92.91194 | 2.76E-21 |
| -0.9399 | NA | 0.307579 | 0.121138 | NA | NA | 4 | -583.842 | 1175.755 | 97.76999 | 2.44E-22 |
| -0.52245 | NA | 0.324829 | NA | NA | NA | 3 | -585.308 | 1176.659 | 98.6742 | 1.55E-22 |
| -0.49199 | NA | 0.332837 | NA | -0.01981 | NA | 4 | -584.844 | 1177.759 | 99.77394 | 8.94E-23 |
| -0.56906 | NA | NA | 0.327855 | -0.062 | NA | 4 | -596.266 | 1200.603 | 122.6184 | 9.79E-28 |
| -0.21746 | NA | NA | 0.192675 | NA | NA | 3 | -598.915 | 1203.873 | 125.8881 | 1.91E-28 |
| 0.535904 | NA | NA | NA | NA | NA | 2 | -602.539 | 1209.1 | 131.1152 | 1.40E-29 |
| 0.539005 | NA | NA | NA | -0.00109 | NA | 3 | -602.538 | 1211.119 | 133.1339 | 5.10E-30 |

#### **Table S33 – Model selection for grasses under drought, expansion, s.d.**

| (Intercept) | I((log(size))^2) | log(Nf) | log(Ng) | log(Nl + exp(-1)) | log(size) | df | logLik | AICc | delta | weight |
| --- | --- | --- | --- | --- | --- | --- | --- | --- | --- | --- |
| **1.657017** | **NA** | **NA** | **-0.27601** | **NA** | **0.113981** | **4** | **-145.128** | **298.4714** | **0** | **0.326858** |
| 1.53731 | NA | 0.040507 | -0.2831 | NA | 0.112076 | 5 | -144.938 | 300.2 | 1.728648 | 0.137717 |
| 1.666376 | NA | NA | -0.27286 | -0.00746 | 0.112347 | 5 | -145.094 | 300.5127 | 2.041318 | 0.117786 |
| 1.647564 | -0.00568 | NA | -0.2745 | NA | 0.129751 | 5 | -145.095 | 300.5134 | 2.04206 | 0.117742 |
| 1.703659 | 0.030422 | NA | -0.27273 | NA | NA | 4 | -146.837 | 301.8884 | 3.41706 | 0.059204 |
| 1.538347 | NA | 0.044736 | -0.27929 | -0.01079 | 0.109513 | 6 | -144.869 | 302.1939 | 3.722567 | 0.050818 |
| 1.52753 | -0.00574 | 0.040583 | -0.28159 | NA | 0.128009 | 6 | -144.903 | 302.2634 | 3.792063 | 0.049082 |
| 1.656925 | -0.00561 | NA | -0.2714 | -0.00737 | 0.127946 | 6 | -145.061 | 302.5793 | 4.107899 | 0.041912 |
| 1.568165 | 0.029811 | 0.045525 | -0.2807 | NA | NA | 5 | -146.6 | 303.5246 | 5.053199 | 0.026126 |
| 1.718413 | 0.029676 | NA | -0.26745 | -0.01267 | NA | 5 | -146.739 | 303.803 | 5.331644 | 0.02273 |
| 1.528717 | -0.00565 | 0.044777 | -0.27783 | -0.0107 | 0.125209 | 7 | -144.835 | 304.2828 | 5.811456 | 0.017882 |
| 1.568622 | 0.028762 | 0.051784 | -0.27497 | -0.01639 | NA | 6 | -146.441 | 305.339 | 6.867658 | 0.010545 |
| 0.589091 | NA | NA | NA | NA | 0.097241 | 3 | -150.466 | 307.0613 | 8.589893 | 0.004457 |
| 1.641562 | NA | NA | -0.23197 | NA | NA | 3 | -150.914 | 307.9568 | 9.485473 | 0.002848 |
| 0.64913 | NA | NA | NA | -0.02078 | 0.09322 | 4 | -150.211 | 308.6381 | 10.16668 | 0.002026 |
| 0.582309 | -0.01072 | NA | NA | NA | 0.127175 | 4 | -150.353 | 308.9204 | 10.44903 | 0.001759 |
| 0.551817 | NA | 0.010261 | NA | NA | 0.09665 | 4 | -150.455 | 309.1245 | 10.65309 | 0.001589 |
| 1.461041 | NA | 0.061217 | -0.2438 | NA | NA | 4 | -150.501 | 309.2172 | 10.74582 | 0.001517 |
| 1.674087 | NA | NA | -0.22342 | -0.02532 | NA | 4 | -150.532 | 309.2783 | 10.80694 | 0.001471 |
| 0.644038 | 0.024703 | NA | NA | NA | NA | 3 | -151.938 | 310.0047 | 11.53334 | 0.001023 |
| 1.468756 | NA | 0.071637 | -0.2357 | -0.02993 | NA | 5 | -149.976 | 310.276 | 11.80464 | 0.000893 |
| 0.641713 | -0.01037 | NA | NA | -0.02049 | 0.12224 | 5 | -150.105 | 310.5338 | 12.06244 | 0.000785 |
| 0.581557 | NA | 0.01989 | NA | -0.0224 | 0.09176 | 5 | -150.169 | 310.6618 | 12.19044 | 0.000737 |
| 0.543375 | -0.01077 | 0.010709 | NA | NA | 0.126698 | 5 | -150.34 | 311.004 | 12.53266 | 0.000621 |
| 0.714645 | 0.023432 | NA | NA | -0.02537 | NA | 4 | -151.56 | 311.3355 | 12.86417 | 0.000526 |
| 0.586648 | 0.024435 | 0.015693 | NA | NA | NA | 4 | -151.911 | 312.0372 | 13.56581 | 0.00037 |
| 0.573014 | -0.01044 | 0.020208 | NA | -0.02213 | 0.120944 | 6 | -150.061 | 312.5777 | 14.10629 | 0.000283 |
| 0.724452 | NA | NA | NA | NA | NA | 2 | -154.581 | 313.225 | 14.75364 | 0.000204 |
| 0.621058 | 0.022859 | 0.027222 | NA | -0.02752 | NA | 5 | -151.481 | 313.2863 | 14.81492 | 0.000198 |
| 0.813546 | NA | NA | NA | -0.03401 | NA | 3 | -153.907 | 313.9415 | 15.47014 | 0.000143 |
| 0.605046 | NA | 0.032161 | NA | NA | NA | 3 | -154.468 | 315.0652 | 16.59385 | 8.15E-05 |
| 0.650083 | NA | 0.046349 | NA | -0.0373 | NA | 4 | -153.678 | 315.5714 | 17.10007 | 6.33E-05 |

#### **Table S34 – Model selection for grasses under drought, colonisation, mean**

| (Intercept) | log(Nf) | log(Ng) | log(Nl + exp(-1)) | df | logLik | AICc | delta | weight |
| --- | --- | --- | --- | --- | --- | --- | --- | --- |
| -1.20115 | 0.16742 | 0.230141 | NA | 4 | -167.31 | 342.7867 | 0 | 0.63623 |
| -1.25773 | 0.170522 | 0.249966 | -0.011 | 5 | -167.215 | 344.6791 | 1.892403 | 0.246993 |
| -0.34533 | 0.18402 | NA | NA | 3 | -170.524 | 347.1466 | 4.359885 | 0.071925 |
| -0.37172 | 0.177042 | NA | 0.016942 | 4 | -170.246 | 348.6586 | 5.871924 | 0.033771 |
| -0.81192 | NA | 0.269142 | NA | 3 | -172.753 | 351.6044 | 8.817758 | 0.007742 |
| -0.80822 | NA | 0.267603 | 0.000824 | 4 | -172.752 | 353.6702 | 10.88351 | 0.002756 |
| 0.251939 | NA | NA | NA | 2 | -177.011 | 358.0709 | 15.28427 | 0.000305 |
| 0.161321 | NA | NA | 0.031306 | 3 | -176.079 | 358.2568 | 15.47009 | 0.000278 |

#### **Table S35 – Model selection for grasses under drought, colonisation, s.d.**

| (Intercept) | log(Nf) | log(Ng) | log(Nl + exp(-1)) | df | logLik | AICc | delta | weight |
| --- | --- | --- | --- | --- | --- | --- | --- | --- |
| **0.76469** | **NA** | **NA** | **NA** | **2** | **-46.4661** | **97.17216** | **0** | **0.313883** |
| 1.519376 | -0.20374 | NA | NA | 3 | -45.5688 | 97.62742 | 0.455259 | 0.249983 |
| 1.039492 | NA | -0.07409 | NA | 3 | -46.3508 | 99.1914 | 2.019241 | 0.114366 |
| 0.793456 | NA | NA | -0.01322 | 3 | -46.4201 | 99.33005 | 2.157896 | 0.106706 |
| 1.584339 | -0.19783 | -0.02342 | NA | 4 | -45.5577 | 99.94871 | 2.776548 | 0.078316 |
| 1.519165 | -0.20352 | NA | -0.00029 | 4 | -45.5688 | 99.97091 | 2.798754 | 0.077451 |
| 1.034316 | NA | -0.06726 | -0.00926 | 4 | -46.3291 | 101.4916 | 4.319436 | 0.036209 |
| 1.586166 | -0.19833 | -0.02387 | 0.000786 | 5 | -45.5575 | 102.3917 | 5.2195 | 0.023087 |

#### **Table S36 – Model selection for legumes under drought, persistence**

| (Intercept) | I((log(size))^2) | log(Nf) | log(Ng) | log(Nl + exp(-1)) | log(size) | df | logLik | AICc | delta | weight |
| --- | --- | --- | --- | --- | --- | --- | --- | --- | --- | --- |
| 6.694131 | NA | NA | -1.49037 | NA | NA | 2 | -70.457 | 145.0193 | 0 | 0.155425 |
| **5.699385** | **NA** | **0.496715** | **-1.70306** | **NA** | **NA** | **3** | **-69.492** | **145.1963** | **0.177006** | **0.14226** |
| 7.116145 | NA | 0.541325 | -1.85027 | -0.31827 | NA | 4 | -68.8482 | 146.0535 | 1.034162 | 0.092673 |
| 7.955481 | NA | NA | -1.5899 | -0.27778 | NA | 3 | -69.9639 | 146.1402 | 1.120904 | 0.08874 |
| 5.883208 | -0.05704 | 0.546837 | -1.75675 | NA | NA | 4 | -69.0066 | 146.3703 | 1.35106 | 0.079094 |
| 6.916535 | -0.04548 | NA | -1.51522 | NA | NA | 3 | -70.1442 | 146.5008 | 1.481557 | 0.074098 |
| 6.802476 | NA | NA | -1.49125 | NA | -0.08962 | 3 | -70.3335 | 146.8794 | 1.86015 | 0.061319 |
| 5.777075 | NA | 0.522031 | -1.71153 | NA | -0.11828 | 4 | -69.2823 | 146.9217 | 1.902397 | 0.060037 |
| 6.924032 | -0.03734 | 0.565739 | -1.85268 | -0.2504 | NA | 5 | -68.6688 | 147.8782 | 2.858933 | 0.037214 |
| 7.849848 | -0.0271 | NA | -1.58516 | -0.22652 | NA | 4 | -69.8691 | 148.0954 | 3.076064 | 0.033386 |
| 7.026496 | NA | 0.547909 | -1.84104 | -0.29192 | -0.04661 | 5 | -68.8198 | 148.1802 | 3.160947 | 0.031998 |
| 7.920061 | NA | NA | -1.58497 | -0.2638 | -0.02384 | 4 | -69.9564 | 148.2699 | 3.250592 | 0.030596 |
| 5.895835 | -0.09732 | 0.55131 | -1.77946 | NA | 0.145886 | 5 | -68.9249 | 148.3903 | 3.371014 | 0.028808 |
| 6.925145 | -0.08289 | NA | -1.5319 | NA | 0.135087 | 4 | -70.0751 | 148.5074 | 3.48808 | 0.02717 |
| 7.003704 | -0.08591 | 0.572475 | -1.88694 | -0.26539 | 0.180283 | 6 | -68.5461 | 149.8558 | 4.836557 | 0.013844 |
| 7.915397 | -0.07207 | NA | -1.60956 | -0.24012 | 0.166326 | 5 | -69.7661 | 150.0727 | 5.053427 | 0.012422 |
| 0.693147 | NA | NA | NA | NA | NA | 1 | -74.4722 | 150.9791 | 5.959809 | 0.007895 |
| -0.2816 | NA | 0.260575 | NA | NA | NA | 2 | -74.1859 | 152.477 | 7.45773 | 0.003733 |
| 0.765359 | -0.02597 | NA | NA | NA | NA | 2 | -74.351 | 152.8072 | 7.787944 | 0.003165 |
| 1.055642 | NA | NA | NA | -0.11655 | NA | 2 | -74.3599 | 152.825 | 7.805704 | 0.003137 |
| 0.765838 | NA | NA | NA | NA | -0.06056 | 2 | -74.4091 | 152.9234 | 7.904121 | 0.002986 |
| -0.26865 | -0.02998 | 0.279369 | NA | NA | NA | 3 | -74.0254 | 154.2632 | 9.243876 | 0.001528 |
| 0.078484 | NA | 0.264788 | NA | -0.12088 | NA | 3 | -74.0641 | 154.3407 | 9.321369 | 0.00147 |
| -0.24704 | NA | 0.275115 | NA | NA | -0.07412 | 3 | -74.0931 | 154.3986 | 9.379295 | 0.001428 |
| 0.999639 | -0.01901 | NA | NA | -0.08155 | NA | 3 | -74.3048 | 154.8219 | 9.802642 | 0.001156 |
| 0.740216 | -0.04151 | NA | NA | NA | 0.056952 | 3 | -74.3379 | 154.8881 | 9.868811 | 0.001118 |
| 1.036564 | NA | NA | NA | -0.09737 | -0.0338 | 3 | -74.3432 | 154.8987 | 9.879423 | 0.001112 |
| -0.03794 | -0.02328 | 0.278139 | NA | -0.0787 | NA | 4 | -73.9818 | 156.3208 | 11.30155 | 0.000546 |
| -0.28854 | -0.04422 | 0.278509 | NA | NA | 0.052281 | 4 | -74.0147 | 156.3865 | 11.36722 | 0.000529 |
| 0.020688 | NA | 0.273474 | NA | -0.09416 | -0.04813 | 4 | -74.0308 | 156.4187 | 11.39946 | 0.00052 |
| 0.985381 | -0.03755 | NA | NA | -0.08731 | 0.069734 | 4 | -74.2853 | 156.9278 | 11.90854 | 0.000403 |
| -0.04733 | -0.0404 | 0.2769 | NA | -0.08374 | 0.064393 | 5 | -73.9658 | 158.4722 | 13.45287 | 0.000186 |

#### **Table S37 – Model selection for legumes under drought, expansion, mean**

| (Intercept) | I((log(size))^2) | log(Nf) | log(Ng) | log(Nl + exp(-1)) | log(size) | df | logLik | AICc | delta | weight |
| --- | --- | --- | --- | --- | --- | --- | --- | --- | --- | --- |
| 0.725033 | 0.194524 | NA | NA | NA | NA | 3 | -96.356 | 199.0363 | 0 | 0.148441 |
| -0.10803 | 0.192919 | 0.221306 | NA | NA | NA | 4 | -95.4309 | 199.4098 | 0.373417 | 0.12316 |
| 0.60469 | 0.121688 | NA | NA | NA | 0.267402 | 4 | -95.687 | 199.922 | 0.885632 | 0.095333 |
| -0.0523 | 0.201486 | NA | 0.194873 | NA | NA | 4 | -95.8624 | 200.2727 | 1.236316 | 0.08 |
| 0.471343 | NA | NA | NA | NA | 0.655802 | 3 | -97.291 | 200.9064 | 1.870043 | 0.058275 |
| **-0.07027** | **0.137617** | **0.186893** | **NA** | **NA** | **0.203943** | **5** | **-95.0567** | **200.9467** | **1.910324** | **0.057113** |
| 0.916458 | 0.200144 | NA | NA | -0.06708 | NA | 4 | -96.2066 | 200.9612 | 1.924866 | 0.056699 |
| -0.61483 | 0.198431 | 0.197537 | 0.149483 | NA | NA | 5 | -95.1451 | 201.1235 | 2.087177 | 0.052279 |
| 0.083895 | 0.198587 | 0.221639 | NA | -0.06769 | NA | 5 | -95.2751 | 201.3836 | 2.347225 | 0.045905 |
| -0.04584 | 0.135164 | NA | 0.166208 | NA | 0.239726 | 5 | -95.3296 | 201.4925 | 2.456184 | 0.043471 |
| 0.809703 | 0.126325 | NA | NA | -0.07268 | 0.272734 | 5 | -95.5088 | 201.851 | 2.814618 | 0.036339 |
| 0.137236 | 0.205147 | NA | 0.182421 | -0.04901 | NA | 5 | -95.7837 | 202.4007 | 3.364401 | 0.027605 |
| 0.035826 | NA | 0.117552 | NA | NA | 0.647866 | 4 | -97.0422 | 202.6323 | 3.595969 | 0.024587 |
| -0.52658 | 0.146876 | 0.168324 | 0.133718 | NA | 0.187982 | 6 | -94.8282 | 202.8396 | 3.803229 | 0.022166 |
| 0.134582 | 0.142155 | 0.186324 | NA | -0.0719 | 0.209411 | 6 | -94.8795 | 202.942 | 3.905667 | 0.02106 |
| 0.162552 | NA | NA | 0.077144 | NA | 0.662919 | 4 | -97.2132 | 202.9744 | 3.938092 | 0.020721 |
| 0.599407 | NA | NA | NA | -0.04655 | 0.668696 | 4 | -97.22 | 202.988 | 3.951645 | 0.020581 |
| -0.4123 | 0.202444 | 0.200089 | 0.135113 | -0.05425 | NA | 6 | -95.047 | 203.2772 | 4.240813 | 0.01781 |
| 0.175541 | 0.137569 | NA | 0.150867 | -0.0572 | 0.246477 | 6 | -95.2214 | 203.6259 | 4.589515 | 0.014961 |
| 0.163432 | NA | 0.115803 | NA | -0.04403 | 0.66018 | 5 | -96.9783 | 204.79 | 5.753618 | 0.008359 |
| -0.13612 | NA | 0.108657 | 0.051188 | NA | 0.653189 | 5 | -97.0092 | 204.8517 | 5.81532 | 0.008105 |
| -0.29979 | 0.149522 | 0.17013 | 0.117296 | -0.05993 | 0.1945 | 7 | -94.7079 | 205.0159 | 5.979552 | 0.007466 |
| 0.315258 | NA | NA | 0.065661 | -0.0388 | 0.672608 | 5 | -97.1656 | 205.1644 | 6.128098 | 0.006932 |
| 0.018117 | NA | 0.10914 | 0.039376 | -0.03953 | 0.663016 | 6 | -96.9594 | 207.1019 | 8.065588 | 0.002631 |
| 0.338511 | NA | NA | NA | 0.290928 | NA | 3 | -117.572 | 241.4673 | 42.431 | 9.07E-11 |
| -0.68671 | NA | 0.274407 | NA | 0.28672 | NA | 4 | -116.745 | 242.0389 | 43.00254 | 6.82E-11 |
| 1.23233 | NA | NA | NA | NA | NA | 2 | -119.441 | 243.0417 | 44.00533 | 4.13E-11 |
| 0.864361 | NA | NA | -0.12086 | 0.273028 | NA | 4 | -117.46 | 243.4681 | 44.43174 | 3.34E-11 |
| 0.159527 | NA | 0.283568 | NA | NA | NA | 3 | -118.599 | 243.5232 | 44.48684 | 3.25E-11 |
| 0.018456 | NA | 0.302723 | -0.18639 | 0.25868 | NA | 5 | -116.483 | 243.7996 | 44.7633 | 2.83E-11 |
| 2.14242 | NA | NA | -0.23361 | NA | NA | 3 | -119.016 | 244.3572 | 45.32087 | 2.14E-11 |
| 1.154868 | NA | 0.327417 | -0.29808 | NA | NA | 4 | -117.912 | 244.3711 | 45.33472 | 2.12E-11 |

#### **Table S38 – Model selection for legumes under drought, expansion, s.d.**

| (Intercept) | I((log(size))^2) | log(Nf) | log(Ng) | log(Nl + exp(-1)) | log(size) | df | logLik | AICc | delta | weight |
| --- | --- | --- | --- | --- | --- | --- | --- | --- | --- | --- |
| -0.34308 | NA | NA | 0.230221 | NA | 0.097414 | 4 | -48.9458 | 106.4395 | 0 | 0.142448 |
| -0.25721 | -0.0557 | NA | 0.19352 | NA | 0.271803 | 5 | -47.8541 | 106.5416 | 0.102145 | 0.135355 |
| **0.500219** | **-0.07139** | **NA** | **NA** | **NA** | **0.304026** | **4** | **-49.4648** | **107.4775** | **1.037995** | **0.084773** |
| -0.05961 | -0.06051 | -0.06919 | 0.206874 | NA | 0.293072 | 6 | -47.5688 | 108.3207 | 1.881203 | 0.055611 |
| -0.05215 | NA | NA | 0.184556 | NA | NA | 3 | -51.0619 | 108.4481 | 2.008637 | 0.052178 |
| -0.22048 | NA | -0.0446 | 0.240876 | NA | 0.101409 | 5 | -48.8273 | 108.488 | 2.048527 | 0.051147 |
| -0.31167 | NA | NA | 0.227859 | -0.00798 | 0.099407 | 5 | -48.9388 | 108.711 | 2.271483 | 0.045752 |
| -0.26453 | 0.019498 | NA | 0.226021 | NA | NA | 4 | -50.1167 | 108.7813 | 2.341831 | 0.044171 |
| 0.578447 | NA | NA | NA | NA | 0.076172 | 3 | -51.266 | 108.8564 | 2.416905 | 0.042543 |
| -0.25513 | -0.05568 | NA | 0.193375 | -0.00054 | 0.271867 | 6 | -47.8541 | 108.8913 | 2.451846 | 0.041806 |
| 0.666836 | NA | NA | NA | NA | NA | 2 | -52.554 | 109.268 | 2.828473 | 0.034631 |
| 0.646336 | -0.07484 | -0.04046 | NA | NA | 0.317764 | 5 | -49.3688 | 109.5709 | 3.131427 | 0.029763 |
| 0.557716 | -0.07009 | NA | NA | -0.02038 | 0.305522 | 5 | -49.419 | 109.6713 | 3.231835 | 0.028306 |
| -0.23052 | NA | NA | 0.200292 | 0.038105 | NA | 4 | -50.8918 | 110.3315 | 3.892063 | 0.020347 |
| -0.02005 | NA | -0.01064 | 0.186652 | NA | NA | 4 | -51.0553 | 110.6586 | 4.219112 | 0.017278 |
| -0.06178 | -0.06054 | -0.0692 | 0.207031 | 0.000572 | 0.293009 | 7 | -47.5688 | 110.7375 | 4.298031 | 0.016609 |
| 0.637044 | 0.011424 | NA | NA | NA | NA | 3 | -52.2186 | 110.7616 | 4.32211 | 0.01641 |
| 0.674393 | NA | NA | NA | -0.03488 | 0.085833 | 4 | -51.1362 | 110.8204 | 4.380877 | 0.015935 |
| -0.19049 | NA | -0.04451 | 0.238579 | -0.00769 | 0.103319 | 6 | -48.8209 | 110.8248 | 4.385361 | 0.0159 |
| -0.1972 | 0.019863 | -0.02364 | 0.231454 | NA | NA | 5 | -50.0838 | 111.0009 | 4.561387 | 0.01456 |
| -0.29738 | 0.018863 | NA | 0.228179 | 0.008494 | NA | 5 | -50.1091 | 111.0514 | 4.611965 | 0.014196 |
| 0.588639 | NA | -0.00275 | NA | NA | 0.076358 | 4 | -51.2656 | 111.0791 | 4.639642 | 0.014001 |
| 0.603218 | NA | 0.016816 | NA | NA | NA | 3 | -52.5377 | 111.3997 | 4.960258 | 0.011927 |
| 0.640904 | NA | NA | NA | 0.008441 | NA | 3 | -52.5454 | 111.4152 | 4.97572 | 0.011836 |
| 0.704902 | -0.07354 | -0.04062 | NA | -0.02055 | 0.319327 | 6 | -49.3221 | 111.8274 | 5.387875 | 0.009631 |
| -0.19043 | NA | -0.01434 | 0.203397 | 0.038785 | NA | 5 | -50.8799 | 112.593 | 6.153568 | 0.006568 |
| 0.677306 | 0.012606 | NA | NA | -0.01411 | NA | 4 | -52.1982 | 112.9443 | 6.504817 | 0.00551 |
| 0.587507 | 0.011328 | 0.01316 | NA | NA | NA | 4 | -52.2086 | 112.9652 | 6.525679 | 0.005453 |
| 0.689978 | NA | -0.00414 | NA | -0.03497 | 0.086137 | 5 | -51.1352 | 113.1037 | 6.664258 | 0.005088 |
| -0.23126 | 0.019189 | -0.02407 | 0.233871 | 0.009124 | NA | 6 | -50.075 | 113.333 | 6.893555 | 0.004537 |
| 0.579056 | NA | 0.016554 | NA | 0.008187 | NA | 4 | -52.5297 | 113.6073 | 7.167815 | 0.003955 |
| 0.627611 | 0.012513 | 0.013229 | NA | -0.01414 | NA | 5 | -52.188 | 115.2094 | 8.76993 | 0.001775 |

#### **Table S39 – Model selection for legumes under drought, colonisation, mean**

| (Intercept) | log(Nf) | log(Ng) | log(Nl + exp(-1)) | df | logLik | AICc | delta | weight |
| --- | --- | --- | --- | --- | --- | --- | --- | --- |
| 0.808066 | NA | NA | -0.14503 | 3 | -24.6456 | 56.55444 | 0 | 0.238435 |
| **-0.57963** | **NA** | **0.380077** | **-0.14611** | **4** | **-23.4161** | **57.05448** | **0.500032** | **0.18569** |
| 0.514641 | NA | NA | NA | 2 | -26.3062 | 57.21246 | 0.658012 | 0.171587 |
| -0.85934 | NA | 0.375732 | NA | 3 | -25.2749 | 57.81298 | 1.258541 | 0.127081 |
| -0.77815 | 0.440984 | NA | -0.16737 | 4 | -23.8289 | 57.88001 | 1.325571 | 0.122893 |
| -1.66965 | 0.351333 | 0.332498 | -0.16377 | 5 | -22.8666 | 59.26268 | 2.70824 | 0.061558 |
| -0.49316 | 0.272416 | NA | NA | 3 | -26.0285 | 59.32013 | 2.765684 | 0.059815 |
| -1.44037 | 0.181641 | 0.350862 | NA | 4 | -25.1455 | 60.51315 | 3.958707 | 0.032942 |

#### **Table S40 – Model selection for legumes under drought, colonisation, s.d.**

| (Intercept) | log(Nf) | log(Ng) | log(Nl + exp(-1)) | df | logLik | AICc | delta | weight |
| --- | --- | --- | --- | --- | --- | --- | --- | --- |
| 0.779809 | NA | NA | -0.10969 | 3 | -5.31827 | 17.8997 | 0 | 0.495113 |
| 0.334969 | NA | 0.121838 | -0.11004 | 4 | -4.65624 | 19.5347 | 1.635005 | 0.218609 |
| **0.230013** | **0.152849** | **NA** | **-0.11744** | **4** | **-4.79814** | **19.8185** | **1.918799** | **0.189689** |
| -0.05144 | 0.124546 | 0.104971 | -0.1163 | 5 | -4.30638 | 22.14217 | 4.242467 | 0.059356 |
| 0.557885 | NA | NA | NA | 2 | -9.80415 | 24.2083 | 6.308606 | 0.021126 |
| 0.124315 | NA | 0.118565 | NA | 3 | -9.38408 | 26.03132 | 8.131618 | 0.008491 |
| 0.429972 | 0.034576 | NA | NA | 3 | -9.78557 | 26.8343 | 8.934604 | 0.005683 |
| 0.111379 | 0.004044 | 0.118011 | NA | 4 | -9.38383 | 28.98987 | 11.09017 | 0.001934 |

#### **Table S41 – Model selection for forbs under drought, persistence**

| (Intercept) | I((log(size))^2) | log(Nf) | log(Ng) | log(Nl + exp(-1)) | log(size) | df | logLik | AICc | delta | weight |
| --- | --- | --- | --- | --- | --- | --- | --- | --- | --- | --- |
| 3.821004 | 0.108906 | -0.829216197 | NA | -0.269851238 | 0.191941 | 5 | -251.085 | 512.3181 | 0 | 0.168396 |
| 2.292898 | NA | -0.798952725 | 0.406474 | -0.328069746 | 0.276588 | 5 | -251.086 | 512.3206 | 0.002515 | 0.168184 |
| 2.778272 | 0.096427 | -0.920293787 | 0.360219 | -0.318481809 | 0.222769 | 6 | -250.17 | 512.548 | 0.22989 | 0.15011 |
| **3.42127** | **NA** | **-0.677721031** | **NA** | **-0.273677524** | **0.250657** | **4** | **-252.283** | **512.6652** | **0.347152** | **0.141562** |
| 3.800954 | 0.138702 | -0.812236661 | NA | -0.264170764 | NA | 4 | -252.284 | 512.6677 | 0.349632 | 0.141387 |
| 2.997597 | 0.132809 | -0.879872919 | 0.275839 | -0.299984819 | NA | 5 | -251.731 | 513.6102 | 1.292103 | 0.088258 |
| 3.223914 | NA | -0.587916815 | NA | -0.268163561 | NA | 3 | -254.503 | 515.0648 | 2.74671 | 0.042647 |
| 2.316704 | NA | -0.677219314 | 0.320458 | -0.309291782 | NA | 4 | -253.74 | 515.5794 | 3.261351 | 0.032971 |
| 3.363025 | 0.113288 | -0.919976919 | NA | NA | 0.18176 | 4 | -254.451 | 517.0006 | 4.682535 | 0.016201 |
| 3.354631 | 0.141712 | -0.902447368 | NA | NA | NA | 3 | -255.532 | 517.1225 | 4.804403 | 0.015243 |
| 2.94552 | NA | -0.765415254 | NA | NA | 0.244221 | 3 | -255.758 | 517.5754 | 5.257292 | 0.012154 |
| 3.072429 | 0.110037 | -0.947748178 | 0.093273 | NA | 0.189141 | 5 | -254.374 | 518.8974 | 6.579354 | 0.006275 |
| 3.235141 | 0.140854 | -0.913540244 | 0.038284 | NA | NA | 4 | -255.518 | 519.136 | 6.817882 | 0.00557 |
| 2.537237 | NA | -0.812234927 | 0.136391 | NA | 0.251987 | 4 | -255.59 | 519.2792 | 6.961131 | 0.005185 |
| 2.760953 | NA | -0.675333034 | NA | NA | NA | 2 | -257.872 | 519.7742 | 7.456144 | 0.004048 |
| 2.524513 | NA | -0.70063646 | 0.077973 | NA | NA | 3 | -257.817 | 521.6927 | 9.374657 | 0.001551 |
| 1.524412 | NA | NA | NA | -0.347790198 | NA | 2 | -261.821 | 527.6724 | 15.35428 | 7.80E-05 |
| 1.494988 | NA | NA | NA | -0.355930796 | 0.120855 | 3 | -261.259 | 528.5776 | 16.25951 | 4.96E-05 |
| 1.900435 | NA | NA | -0.10139 | -0.333081345 | NA | 3 | -261.733 | 529.5262 | 17.20809 | 3.09E-05 |
| 1.536244 | -0.02156 | NA | NA | -0.343967539 | NA | 3 | -261.741 | 529.5418 | 17.22374 | 3.06E-05 |
| 1.513285 | -0.05106 | NA | NA | -0.349713852 | 0.160187 | 4 | -260.879 | 529.8566 | 17.53856 | 2.62E-05 |
| 1.843533 | NA | NA | -0.09385 | -0.342233371 | 0.119169 | 4 | -261.184 | 530.4671 | 18.14902 | 1.93E-05 |
| 1.828846 | -0.01632 | NA | -0.07972 | -0.333270511 | NA | 4 | -261.691 | 531.482 | 19.16386 | 1.16E-05 |
| 1.609941 | -0.04892 | NA | -0.02629 | -0.346077142 | 0.158082 | 5 | -260.874 | 531.896 | 19.57789 | 9.44E-06 |
| 2.039769 | NA | NA | -0.36147 | NA | NA | 2 | -266.794 | 537.618 | 25.29988 | 5.40E-07 |
| 0.556735 | NA | NA | NA | NA | NA | 1 | -268.244 | 538.4984 | 26.18029 | 3.48E-07 |
| 2.003249 | NA | NA | -0.36176 | NA | 0.08745 | 3 | -266.491 | 539.0416 | 26.72354 | 2.65E-07 |
| 1.973557 | -0.01626 | NA | -0.34126 | NA | NA | 3 | -266.752 | 539.5628 | 27.24473 | 2.04E-07 |
| 0.60053 | -0.04242 | NA | NA | NA | NA | 2 | -267.927 | 539.8833 | 27.5652 | 1.74E-07 |
| 0.519089 | NA | NA | NA | NA | 0.087901 | 2 | -267.944 | 539.9184 | 27.60035 | 1.71E-07 |
| 0.566224 | -0.06887 | NA | NA | NA | 0.141953 | 3 | -267.237 | 540.5339 | 28.21584 | 1.26E-07 |
| 1.825052 | -0.04067 | NA | -0.31162 | NA | 0.119607 | 4 | -266.272 | 540.6421 | 28.32396 | 1.19E-07 |

#### **Table S42 – Model selection for forbs under drought, expansion, mean**

| (Intercept) | I((log(size))^2) | log(Nf) | log(Ng) | log(Nl + exp(-1)) | log(size) | df | logLik | AICc | delta | weight |
| --- | --- | --- | --- | --- | --- | --- | --- | --- | --- | --- |
| **-0.09385** | **0.116875** | **NA** | **0.162862** | **-0.07483** | **0.218547** | **6** | **-358.809** | **729.8689** | **0** | **0.442293** |
| 0.194516 | 0.126714 | -0.10147 | 0.179365 | -0.06993 | 0.213184 | 7 | -358.08 | 730.497 | 0.628141 | 0.323081 |
| 0.498875 | 0.12271 | NA | NA | -0.05984 | 0.20802 | 5 | -361.218 | 732.6141 | 2.745255 | 0.112095 |
| 0.721816 | 0.129384 | -0.06493 | NA | -0.05574 | 0.203906 | 6 | -360.914 | 734.0791 | 4.210243 | 0.053885 |
| 0.36303 | 0.128929 | -0.14039 | 0.128417 | NA | 0.212877 | 6 | -362.033 | 736.3168 | 6.447978 | 0.017601 |
| 0.355384 | 0.119268 | NA | NA | NA | 0.213216 | 4 | -364.39 | 736.8984 | 7.02953 | 0.01316 |
| -0.03017 | 0.115161 | NA | 0.099883 | NA | 0.220471 | 5 | -363.43 | 737.0386 | 7.169748 | 0.012269 |
| 0.737664 | 0.130616 | -0.10662 | NA | NA | 0.205876 | 5 | -363.544 | 737.2681 | 7.399198 | 0.010939 |
| 0.040226 | 0.185321 | NA | 0.144818 | -0.07561 | NA | 5 | -364.299 | 738.7777 | 8.908849 | 0.005143 |
| 0.377448 | 0.194974 | -0.12003 | 0.164863 | -0.0698 | NA | 6 | -363.307 | 738.8657 | 8.996853 | 0.004921 |
| 0.564338 | 0.18759 | NA | NA | -0.06218 | NA | 4 | -366.156 | 740.432 | 10.56315 | 0.002249 |
| 0.856485 | 0.194696 | -0.08558 | NA | -0.05671 | NA | 5 | -365.64 | 741.4597 | 11.59082 | 0.001345 |
| 0.545378 | 0.197086 | -0.15885 | 0.114033 | NA | NA | 5 | -367.127 | 744.4331 | 14.5642 | 0.000304 |
| 0.873935 | 0.196592 | -0.12821 | NA | NA | NA | 4 | -368.289 | 744.698 | 14.82912 | 0.000266 |
| 0.416792 | 0.185696 | NA | NA | NA | NA | 3 | -369.485 | 745.0421 | 15.17328 | 0.000224 |
| 0.105772 | 0.184198 | NA | 0.081012 | NA | NA | 4 | -368.869 | 745.8578 | 15.98892 | 0.000149 |
| -0.24298 | NA | NA | 0.198131 | -0.07236 | 0.459904 | 5 | -369.019 | 748.2176 | 18.34875 | 4.58E-05 |
| -0.29773 | NA | 0.020142 | 0.194266 | -0.07338 | 0.456935 | 6 | -368.99 | 750.2305 | 20.36166 | 1.68E-05 |
| 0.476837 | NA | NA | NA | -0.05378 | 0.461777 | 4 | -372.404 | 752.9272 | 23.05839 | 4.35E-06 |
| -0.17925 | NA | NA | 0.136698 | NA | 0.45834 | 4 | -373.098 | 754.3152 | 24.44638 | 2.17E-06 |
| 0.26313 | NA | 0.062574 | NA | -0.05805 | 0.452441 | 5 | -372.11 | 754.3994 | 24.53056 | 2.08E-06 |
| 0.34807 | NA | NA | NA | NA | 0.460045 | 3 | -374.816 | 755.7026 | 25.83379 | 1.09E-06 |
| -0.1298 | NA | -0.0185 | 0.141037 | NA | 0.461086 | 5 | -373.073 | 756.325 | 26.45614 | 7.96E-07 |
| 0.275092 | NA | 0.020392 | NA | NA | 0.456958 | 4 | -374.783 | 757.6858 | 27.8169 | 4.03E-07 |
| -0.75663 | NA | 0.282165 | 0.158135 | -0.08298 | NA | 5 | -420.067 | 850.3122 | 120.4433 | 3.10E-27 |
| -0.29557 | NA | 0.314667 | NA | -0.07041 | NA | 4 | -421.609 | 851.3371 | 121.4682 | 1.86E-27 |
| -0.28781 | NA | 0.266457 | NA | NA | NA | 3 | -424.554 | 855.1798 | 125.3109 | 2.72E-28 |
| -0.57121 | NA | 0.241112 | 0.097494 | NA | NA | 4 | -423.944 | 856.0074 | 126.1385 | 1.80E-28 |
| 0.035501 | NA | NA | 0.213688 | -0.06844 | NA | 4 | -424.728 | 857.5747 | 127.7058 | 8.22E-29 |
| 0.094885 | NA | NA | 0.155528 | NA | NA | 3 | -427.371 | 860.8127 | 130.9438 | 1.63E-29 |
| 0.813306 | NA | NA | NA | -0.04837 | NA | 3 | -427.573 | 861.2174 | 131.3485 | 1.33E-29 |
| 0.696313 | NA | NA | NA | NA | NA | 2 | -428.989 | 862.0141 | 132.1452 | 8.93E-30 |

#### **Table S43 – Model selection for forbs under drought, expansion, s.d.**

| (Intercept) | I((log(size))^2) | log(Nf) | log(Ng) | log(Nl + exp(-1)) | log(size) | df | logLik | AICc | delta | weight |
| --- | --- | --- | --- | --- | --- | --- | --- | --- | --- | --- |
| 0.530842 | NA | NA | NA | -0.02008 | 0.090307 | 4 | -180.254 | 368.627 | 0 | 0.14819 |
| 0.482772 | NA | NA | NA | NA | 0.08966 | 3 | -181.295 | 368.662 | 0.035057 | 0.145615 |
| 0.330261 | NA | 0.058731 | NA | -0.02409 | 0.081544 | 5 | -179.454 | 369.0863 | 0.459301 | 0.117783 |
| **0.335224** | **NA** | **0.041229** | **NA** | **NA** | **0.083418** | **4** | **-180.883** | **369.8858** | **1.258851** | **0.07897** |
| 0.452125 | NA | NA | 0.021667 | -0.02211 | 0.090102 | 5 | -180.13 | 370.4395 | 1.812563 | 0.059872 |
| 0.531765 | 0.005134 | NA | NA | -0.02033 | 0.079689 | 5 | -180.195 | 370.57 | 1.943029 | 0.056091 |
| 0.483016 | 0.003965 | NA | NA | NA | 0.081455 | 4 | -181.261 | 370.6403 | 2.01333 | 0.054154 |
| 0.471596 | NA | NA | 0.002897 | NA | 0.089624 | 4 | -181.293 | 370.7051 | 2.078111 | 0.052428 |
| 0.298931 | NA | 0.056361 | 0.010852 | -0.02494 | 0.081795 | 6 | -179.424 | 371.0989 | 2.471951 | 0.043057 |
| 0.3267 | -0.001 | 0.059721 | NA | -0.02411 | 0.083473 | 6 | -179.452 | 371.1546 | 2.52765 | 0.041874 |
| 0.356018 | NA | 0.043226 | -0.00724 | NA | 0.083206 | 5 | -180.869 | 371.9179 | 3.290913 | 0.02859 |
| 0.333553 | -0.00047 | 0.041688 | NA | NA | 0.084325 | 5 | -180.883 | 371.945 | 3.318023 | 0.028205 |
| 0.457746 | 0.004406 | NA | 0.020338 | -0.0222 | 0.081004 | 6 | -180.088 | 372.4266 | 3.799662 | 0.022168 |
| 0.476641 | 0.003897 | NA | 0.001652 | NA | 0.081575 | 5 | -181.26 | 372.6988 | 4.071854 | 0.019348 |
| 0.556842 | 0.029989 | NA | NA | -0.02123 | NA | 4 | -182.309 | 372.7363 | 4.109326 | 0.018988 |
| 0.506475 | 0.029342 | NA | NA | NA | NA | 3 | -183.456 | 372.9839 | 4.356907 | 0.016778 |
| 0.294393 | -0.00117 | 0.057482 | 0.010989 | -0.02498 | 0.084042 | 7 | -179.421 | 373.1783 | 4.55133 | 0.015223 |
| 0.381829 | 0.025732 | 0.051267 | NA | -0.0245 | NA | 5 | -181.764 | 373.7071 | 5.080165 | 0.011686 |
| 0.354576 | -0.00038 | 0.043583 | -0.00721 | NA | 0.083932 | 6 | -180.869 | 373.9897 | 5.362741 | 0.010146 |
| 0.389369 | 0.026551 | 0.032845 | NA | NA | NA | 4 | -183.223 | 374.5656 | 5.938678 | 0.007608 |
| 0.507442 | 0.029775 | NA | 0.01365 | -0.02249 | NA | 5 | -182.26 | 374.6998 | 6.072893 | 0.007114 |
| 0.526941 | 0.029441 | NA | -0.00533 | NA | NA | 4 | -183.448 | 375.0158 | 6.388881 | 0.006074 |
| 0.366509 | 0.025741 | 0.050165 | 0.005272 | -0.02492 | NA | 6 | -181.757 | 375.7656 | 7.138621 | 0.004175 |
| 0.426471 | 0.026495 | 0.036305 | -0.01288 | NA | NA | 5 | -183.18 | 376.5382 | 7.911241 | 0.002837 |
| 0.229566 | NA | 0.104166 | NA | -0.02632 | NA | 4 | -185.133 | 378.3848 | 9.757869 | 0.001127 |
| 0.232466 | NA | 0.086148 | NA | NA | NA | 3 | -186.786 | 379.6428 | 11.01588 | 0.000601 |
| 0.216783 | NA | 0.103265 | 0.004384 | -0.02666 | NA | 5 | -185.128 | 380.4354 | 11.80848 | 0.000404 |
| 0.550643 | NA | NA | NA | NA | NA | 2 | -188.67 | 381.3747 | 12.74773 | 0.000253 |
| 0.276362 | NA | 0.090074 | -0.0151 | NA | NA | 4 | -186.727 | 381.5728 | 12.94583 | 0.000229 |
| 0.596644 | NA | NA | NA | -0.01902 | NA | 3 | -187.775 | 381.6205 | 12.99351 | 0.000224 |
| 0.506683 | NA | NA | 0.024715 | -0.02134 | NA | 4 | -187.62 | 383.36 | 14.73309 | 9.37E-05 |
| 0.5252 | NA | NA | 0.006579 | NA | NA | 3 | -188.658 | 383.3868 | 14.75983 | 9.24E-05 |

#### **Table S44 – Model selection for forbs under drought, colonisation, mean**

| (Intercept) | log(Nf) | log(Ng) | log(Nl + exp(-1)) | df | logLik | AICc | delta | weight |
| --- | --- | --- | --- | --- | --- | --- | --- | --- |
| -0.97488 | 0.358523 | NA | NA | 3 | -111.234 | 228.6906 | 0 | 0.426188 |
| **-1.40088** | **0.341071** | **0.130061** | **NA** | **4** | **-110.509** | **229.3916** | **0.70099** | **0.300181** |
| -0.97784 | 0.356305 | NA | 0.00426 | 4 | -111.228 | 230.8308 | 2.140226 | 0.146169 |
| -1.41922 | 0.335259 | 0.133477 | 0.010283 | 5 | -110.476 | 231.5176 | 2.826994 | 0.103688 |
| -0.29758 | NA | 0.173886 | NA | 3 | -115.121 | 236.4644 | 7.773788 | 0.008741 |
| 0.35875 | NA | NA | NA | 2 | -116.331 | 236.7715 | 8.080933 | 0.007496 |
| -0.42142 | NA | 0.182787 | 0.034292 | 4 | -114.767 | 237.9073 | 9.216729 | 0.004248 |
| 0.28517 | NA | NA | 0.027961 | 3 | -116.098 | 238.4189 | 9.728295 | 0.003289 |

#### **Table S45 – Model selection for forbs under drought, colonisation, s.d.**

| (Intercept) | log(Nf) | log(Ng) | log(Nl + exp(-1)) | df | logLik | AICc | delta | weight |
| --- | --- | --- | --- | --- | --- | --- | --- | --- |
| -0.5861 | 0.279935 | NA | NA | 3 | -65.7019 | 137.6259 | 0 | 0.496858 |
| **-0.78721** | **0.271697** | **0.0614** | **NA** | **4** | **-65.3385** | **139.0509** | **1.424998** | **0.243668** |
| -0.58192 | 0.283065 | NA | -0.00601 | 4 | -65.6763 | 139.7264 | 2.100502 | 0.173826 |
| -0.78134 | 0.273556 | 0.060307 | -0.00329 | 5 | -65.331 | 141.228 | 3.602031 | 0.082047 |
| 0.455204 | NA | NA | NA | 2 | -72.5945 | 149.299 | 11.67307 | 0.00145 |
| 0.091682 | NA | 0.09631 | NA | 3 | -71.787 | 149.7963 | 12.17035 | 0.001131 |
| 0.421472 | NA | NA | 0.012819 | 3 | -72.4879 | 151.1981 | 13.57214 | 0.000561 |
| 0.032814 | NA | 0.100541 | 0.016301 | 4 | -71.6137 | 151.6013 | 13.97535 | 0.000459 |

### **Details of model implementation to find overall abundance equilibrium**

We ran our IPMs for 300 time-steps, since the values of *N_g_*, *N_l_*, and *N_f_* for each functional group (and therefore, their corresponding IPMs) did not change significantly when running for further time-steps (changes in overall abundance and the value of the dominant eigenvalue were <10^-6^ units for the last 10 timesteps for each functional group $\times$ treatment). The percentage-cover stage-structure of each functional group *k* at time-step *t* was represented as a 200-entry vector *V_k, t,_* where the *i-*th entry is the number of species of functional group *k* at time *t* with cover between i/2 – 0.5% and *i/2%*. For each treatment, *V_k,0_* was calculated from the distribution of percentage covers of species in each replicate block and taking the element-wise mean. At *t*, to extract the overall abundance values *N_k, t_* from the percentage-cover distribution vectors *V_k, t_* we summed all entries of *V_k, t_*, with each entry weighted by the corresponding midpoint. Then, we used the values of *N_g, t_*, *N_l, t_* and *N_f, t_* to generate *K_t_*, the projection kernel that operated from time *t* to *t+1*. Finally, the new percentage-cover distributions *V_k, t + 1_* were calculated as *V_k, t +1_* = *K_t_* ⋅ *V_k, t_* (Ellner et al. 2016), thus starting the next time-step.

### **Construction of the overall community matrices for each treatment**

The discretized form of equation 8 in the main text is

$$\begin{aligned} n_{k,y}\left( t+1 \right)=\sum_{x} K_{k,yx}\left( t \right)n_{k,x}\left( t \right),\#\left( S1 \right) \end{aligned}$$

where *n _k, x_* (*t*) is the number of species from functional group *k* with percentage-cover *x* at time *t*, and *K_k, yx_* (*t*) is the (y, x)-element of the discretized IPM kernel for group *k* at time-step *t*. To obtain the community matrix, we compute the Jacobian matrix and evaluate it at equilibrium. For a given treatment, the entries of the Jacobian *J* are defined as

$$\begin{aligned} J_{\left( k,y \right),\left( j,z \right)}=\frac{\partial n_{k,y}\left( t+1 \right)}{\partial n_{j,z}\left( t \right)}.\#\left( S2 \right) \end{aligned}$$

Substituting *n_k, y_* (*t* + 1) from equation S1into equation S2 gives

$$\begin{aligned} J_{\left( k,y \right),\left( j,z \right)}=\frac{\partial}{\partial n_{j,z}\left( t \right)}\sum_{x} K_{k,yx}\left( t \right) n_{k,x}\left( t \right)=\sum_{x} \frac{\partial K_{k,yx}\left( t \right)}{\partial n_{j,z}\left( t \right)}n_{k,x}\left( t \right)+\sum_{x} K_{k,yx}\left( t \right)\delta_{kj}\delta_{xz},\#\left( S3 \right) \end{aligned}$$

where *ẟ_kj_* is the Kronecker delta (equal to 1 if *k = j*, and equal to 0 if *k ≠ j*). Calculating the second sum in equation S3, we then obtain

$$\begin{aligned} J_{\left( k,y \right),\left( j,z \right)}=\sum_{x} \left[ {\frac{\partial K_{k,yx}\left( t \right)}{\partial n_{j,z}\left( t \right)}n}_{k,x}\left( t \right) \right]+K_{k,yz}\left( t \right)\delta_{kj}\#\left( S4 \right) \end{aligned}$$

Then, at equilibrium, substituting *n_j, z_(t) = n_j, z_* and *K_k, yz_* *(t) = K_k, yz_* into equation S4 leads us to the community matrix *C _(k, y), (j, z)_* for the treatment in question, given by

$$\begin{aligned} C_{\left( k,y \right),\left( j,z \right)}=\sum_{x} \left[ {\frac{\partial K_{k,yx}}{\partial n_{j,z}}n}_{k,x} \right]+K_{k,yz}\delta_{kj},\#\left( S5 \right) \end{aligned}$$

where all quantities are understood to be evaluated at equilibrium. We now seek the analytic form of *∂K_k, yx_/∂_nj, z_*. For any functional group *k*, by definition, at equilibrium

$$\begin{aligned} N_{k}=\log\left( \sum_{x} \left[ e^{x}n_{k,x} \right] + {e^{-1}\delta}_{kl} \right)\#\left( S6 \right) \end{aligned}$$

Here, the Kronecker delta term represents the *e^-^*^1^ augmentation applied to the overall abundance for legumes. Then,

$$\begin{aligned} \frac{\partial N_{k}}{\partial n_{j,z}}=\frac{\partial}{\partial n_{j,z}}\left( \sum_{x} \left[ e^{x}n_{k,x} \right] + {e^{-1}\delta}_{kl} \right)\frac{1}{\sum_{x} \left[ e^{x}n_{k,x} \right] + {e^{-1}\delta}_{kl}}=\frac{1}{e^{N_{k}}}\sum_{x} e^{x}\delta_{kj}\delta_{xz}=\frac{e^{z}}{e^{N_{k}}}\delta_{kj}\#\left( S7 \right) \end{aligned}$$

By using the generalised chain rule and substituting equation S7 into S5, we obtain

$$\begin{aligned} C_{\left( k,y \right),\left( j,z \right)}=\sum_{x} \left[ {\frac{\partial K_{k,yx}}{\partial N_{j}}\frac{\partial N_{j}}{\partial n_{j,z}}n}_{k,x} \right]+K_{k,yz}\delta_{kj}=\sum_{x} \left[ {\frac{\partial K_{k,yx}}{\partial N_{j}}\frac{e^{z}}{e^{N_{j}}}n}_{k,x} \right]+K_{k,yz}\delta_{kj}\#\left( S8 \right) \end{aligned}$$

Then, using the expressions for *s_k_*(*x*), *g_k_*(*x, y*) and *f_k_*(*y*) from equations (1–4) in the main text, we obtain the following partial derivatives

$$\begin{aligned} \frac{\partial s_{k}\left( x \right)}{\partial N_{j}}=\beta_{{k, s, N}_{j}}s_{k}\left( x \right)\left[ 1-s_{k}\left( x \right) \right]\#\left( S9 \right) \end{aligned}$$

$$\frac{\partial g_{k}\left( x,y \right)}{\partial N_{j}} =exp\left[ \frac{-\left( y-\mu_{g, k}\left( x \right) \right)^{2}}{2{\sigma_{g, k}\left( x \right)}^{2}} \right] \frac{\partial}{\partial N_{j}}\frac{1}{\sigma_{g, k}\left( x \right)\sqrt{2\pi}}+ \frac{1}{\sigma_{g, k}\left( x \right)\sqrt{2\pi}} \frac{\partial}{\partial N_{j}}exp\left[ \frac{-\left( y-\mu_{g, k}\left( x \right) \right)^{2}}{2{\sigma_{g, k}\left( x \right)}^{2}} \right]=-\frac{\beta_{k, \sigma_{g} ,N_{j}}}{{\sigma_{g, k}\left( x \right)}^{2}\sqrt{2\pi}}exp\left[ \frac{-\left( y-\mu_{g, k}\left( x \right) \right)^{2}}{2{\sigma_{g, k}\left( x \right)}^{2}} \right]+\frac{1}{\sigma_{g, k}\left( x \right)\sqrt{2\pi}} exp\left[ \frac{-\left( y-\mu_{g, k}\left( x \right) \right)^{2}}{2{\sigma_{g, k}\left( x \right)}^{2}} \right]\frac{\partial}{\partial N_{j}}\left[ \frac{-\left( y-\mu_{g, k}\left( x \right) \right)^{2}}{2{\sigma_{g, k}\left( x \right)}^{2}} \right]=- \frac{\beta_{k, \sigma_{g} ,N_{j}}}{\sigma_{g, k}\left( x \right)}g_{k}\left( x,y \right) + g_{k}\left( x,y \right) \frac{\partial}{\partial N_{j}}\left[ \frac{-\left( y-\mu_{g, k}\left( x \right) \right)^{2}}{2{\sigma_{g, k}\left( x \right)}^{2}} \right]$$

$$\begin{aligned} =g_{k}\left( x,y \right)\left[ - \frac{\beta_{k, \sigma_{g} ,N_{j}}}{\sigma_{g, k}\left( x \right)} +\frac{\beta_{k, \mu_{g} ,N_{j}}\sigma_{g, k}\left( x \right)\left( y-\mu_{g, k}\left( x \right) \right)+\beta_{k, \sigma_{g} ,N_{j}}\left( y-\mu_{g, k}\left( x \right) \right)^{2}}{{\sigma_{g, k}\left( x \right)}^{3}} \right]\#\left( S10 \right) \end{aligned}$$

By equations (5–7), and the same reasoning as S10, it follows that

$$\begin{aligned} \frac{\partial f_{k}\left( y \right)}{\partial N_{j}}=f_{k}\left( y \right)\left[ - \frac{\beta_{k, \sigma_{f} ,N_{j}}}{\sigma_{f, k}} +\frac{\beta_{k, \mu_{f} ,N_{j}}\sigma_{f, k}\left( y-\mu_{f, k} \right)+\beta_{k, \sigma_{f} ,N_{j}}\left( y-\mu_{f, k} \right)^{2}}{{\sigma_{f, k}}^{3}} \right]\#\left( S11 \right) \end{aligned}$$

Finally, by the product rule,

$$\begin{aligned} \frac{\partial K_{k,yx}}{\partial N_{j}}=\frac{\partial}{\partial N_{j}}\left[ s_{k}\left( x \right)g_{k}\left( x,y \right)+f_{k}\left( y \right) \right]=\frac{\partial s_{k}\left( x \right)}{\partial N_{j}}g_{k}\left( x,y \right)+s_{k}\left( x \right)\frac{\partial g_{k}\left( x,y \right)}{\partial N_{j}}+\frac{\partial f_{k}\left( y \right)}{\partial N_{j}}\#\left( S12 \right) \end{aligned}$$

Finally, substituting equations S9–S11 into S12, and then substituting that into S8, we obtain the analytic form of *C _(k, y), (j, z)_* for every treatment. The resulting matrix is a 600 × 600 matrix whose 200⋅ (*k* – 1) + *y*-th row and 200⋅ (*j* – 1) + *z*-th column is *C _(k, y), (j, z)_,* where *k* = 1 and *j* = 1 correspond to grasses, *k* = 2 and *j* = 2 to legumes, and *k* = 3 and *j* = 3 to forbs. Hence, for example, for a given treatment *K_1_* is the (equilibrium, and hence time-independent) kernel surface for grasses, and *n_2, z_* is the equilibrium number of legume species of cover *z*. Figure 5j-l in the main text show the eigenvalues and pseudospectra of the resulting community matrices for each treatment.

**Figure S1. Values of *T_ϵ_* for ϵ = 10^-4^ to 10^-1^ for   each functional group × treatment, indicating degree of transient instability.** Values of *T_ϵ_* < 0 are not shown as they are uninformative. For every functional group × treatment (a – c), and for each treatment overall (d – f), sup*_ϵ_*_≥0_ *T_ϵ_* > 1, since all points like above the dashed line representing *T_ϵ_* = 1. This implies every functional group × treatment, and every treatment overall, is transiently unstable — *i.e.*, they all transiently amplify small perturbations. Then, since all functional group × treatments, and all treatments overall, are asymptotically stable (Figure 5), as *t* → ∞*,* the perturbation is eventually suppressed. (a, b) Under Irrigation and control, Legumes are more transiently unstable than forbs, which in turn are more transiently unstable than grasses. (c) Under drought, grasses are more transiently unstable than forbs, which in turn are more transiently unstable than legumes. (d – f) Each treatment overall has a significantly greater degree of transient instability than individual functional groups as their values of sup*_ϵ_*_≥0_ *T_ϵ_* are much larger, especially the drought treatment (f). This potentially indicates that individual functional-group transient instabilities can amplify each other transiently to produce a much greater transient instability in the community as a whole.

**Figure S1**

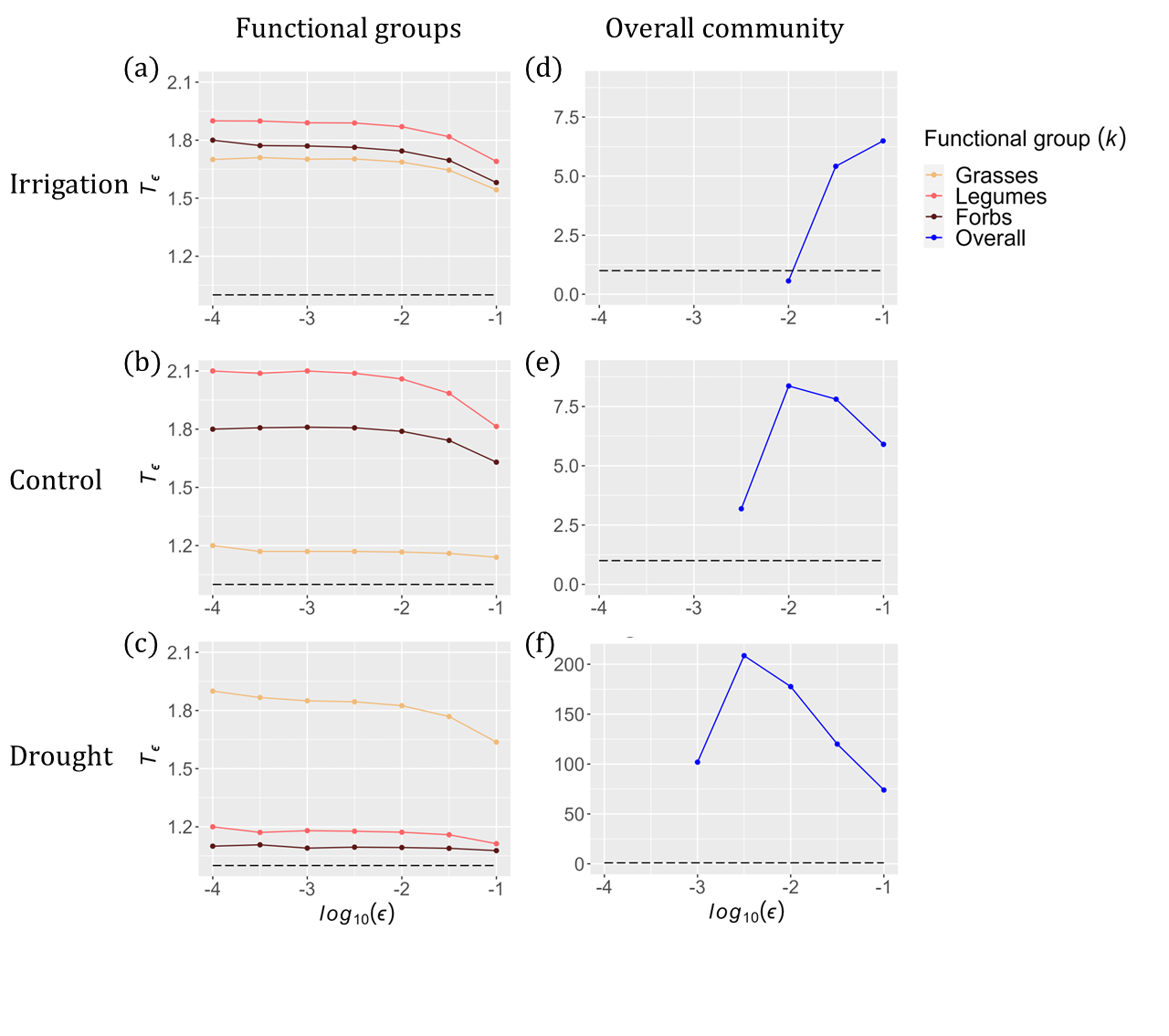
